# Towards phage therapy for acne vulgaris: Topical application in a mouse model

**DOI:** 10.1101/2022.02.19.481124

**Authors:** Amit Rimon, Chani Rakov, Vanda Lerer, Sivan Sheffer-Levi, Sivan Alkalky-Oren, Tehila Shlomov, Lihi Shasha, Ruthi Lubin, Shunit Coppenhagen-Glazer, Vered Molho-Pessach, Ronen Hazan

**Author notes:** These authors contributed equally (Amit Rimon, Chani Rakov, Vered Molho-Pessach, Ronen Hazan).

## Abstract

Acne vulgaris is a common neutrophile-driven inflammatory skin disorder in which *Cutibacterium acnes* (*C. acnes*) bacteria play a significant role. Until now, antibiotics have been widely used to treat acne vulgaris, with the inevitable increase in bacterial antibiotic resistance. Phage therapy is a promising solution to the rising problem of antibiotic-resistant bacteria, utilizing viruses that specifically lyse bacteria.

Here, we explored the feasibility of phage therapy against *C. acnes*. By combining eight novel phages we had isolated, together with commonly used antibiotics, we achieved 100% eradication of clinically isolated *C. acnes* strains. Using topical phage therapy in an acne mouse model resulted in significantly superior clinical scores, as well as a reduction in neutrophil infiltration compared to the control group. These results demonstrate the potential of phage therapy in acne vulgaris treatment, especially when antibiotic-resistant strains are involved.

## INTRODUCTION

*Cutibacterium acnes* (*C. acnes*, previously termed *Propionibacterium acnes*) is a gram-positive, lipophilic, anaerobic bacterium that is a skin microbiome resident (O’Neill and Gallo, 2018). *C. acnes* plays a significant role in the pathogenesis of acne vulgaris, a common chronic inflammatory disorder of the pilosebaceous unit (Dessinioti and Katsambas, 2010), affecting 80% of the population during adolescence (Rzany and Kahl, 2006) as well as some adults (Zaenglein, 2018). Although strains of *C. acnes* associated with healthy skin have been identified (phylotypes II and III) (Lomholt and Kilian, 2010), other strains (phylotype IA) have been linked to acne (Lomholt and Kilian, 2010).

The complex pathogenesis of acne involves androgen-mediated stimulation of sebaceous glands, follicular hyperkeratinization, dysbiosis within the pilosebaceous microbiome, and innate and cellular immune responses (O’Neill and Gallo, 2018).

*C. acnes* activates the innate immune response to produce proinflammatory interleukin (IL)-1 by activating the nod-like receptor P3 (NLRP3) inflammasome in human sebocytes and monocytes (Li et al., 2014). Moreover, *C. acnes* activates Toll-like receptor-2 in monocytes and triggers the secretion of the proinflammatory cytokines IL-12 and IL-8. IL-8 attracts neutrophils and leads to the release of lysosomal enzymes. These neutrophil-derived enzymes result in the rupture of the follicular epithelium and further inflammation (Kim et al., 2002).

Acne has been treated for decades with topical and oral antibiotics, such as tetracycline (TET), doxycycline (DOX), minocycline (MC), erythromycin (EM), and clindamycin (CM), aimed at *C. acnes* (Muhammad and Rosen, 2013). Topical treatment modality is preferred when possible with level A recommendation strength (Zaenglein et al., 2016), due to its local effect and lack of systemic side effects (Zaenglein et al., 2016). However, an alarming global increase in antibiotic-resistant *C. acnes* strains has occurred during the last two dozen years (Swanson, 2003). For example, we (Sheffer-Levi et al., 2020) examined the sensitivity profile of 36 clinical isolates of *C. acnes*, representing the verity of strains in Israel, to the abovementioned commonly used antibiotics and found that the antibiotic resistance in this collection was 30.6% for at least one of these antibiotics. These results correlate with the worldwide reported data of 20%–60% of resistant strains (Karadag et al., 2020; Sheffer-Levi et al., 2020), indicating the need for other treatment strategies aimed at *C. acnes*.

Bacteriophage (phage) therapy is evolving as one of the most promising solutions for emerging antibiotic resistance. Phages are bacterial viruses widely distributed in the environment that replicate within bacteria and specifically kill their bacterial targets without harming other flora members. Therefore, they can be used as living drugs for various bacterial infections, including acne (Jassim and Limoges, 2014).

The first *C. acnes* phage was isolated in 1964 by Brzin (BRZIN, 1964), but only in recent years has phage therapy become a potential treatment approach for acne vulgaris, as reflected by the increase in academic and industrial performance publications. Nevertheless, research on this topic is lacking, as *in vivo* phage therapy in an acne mouse model has been tested only by intralesional injections of *C. acnes* phages, and the efficacy of topical application, has not yet been shown (Kim et al., 2019; Lam et al., 2021; Nelson et al., 2012).

Here, we present a direct topical application of *C. acnes* phages in an *in vivo* mouse model of acne vulgaris as proof of the concept of phage therapy for acne. We tested our collection of *C. acnes* strains for their *in vitro* susceptibility to eight novel phages we had isolated. Using an acne mouse model, we then performed an *in vivo* evaluation of the efficacy and safety of topical application of phages. To the best of our knowledge, this is the first demonstration of phage therapy for *C. acnes* using a topical application.

## RESULTS

### Phage isolation

As part of the routine work of the Israeli Phage Bank (Yerushalmy et al., 2020), a screen for potential phages targeting *C. acnes* was performed. Eight phages targeting *C. acnes* were isolated from acne patients’ saliva and Skin samples (Table 1). The phages were characterized as follows.

**Table 1.**
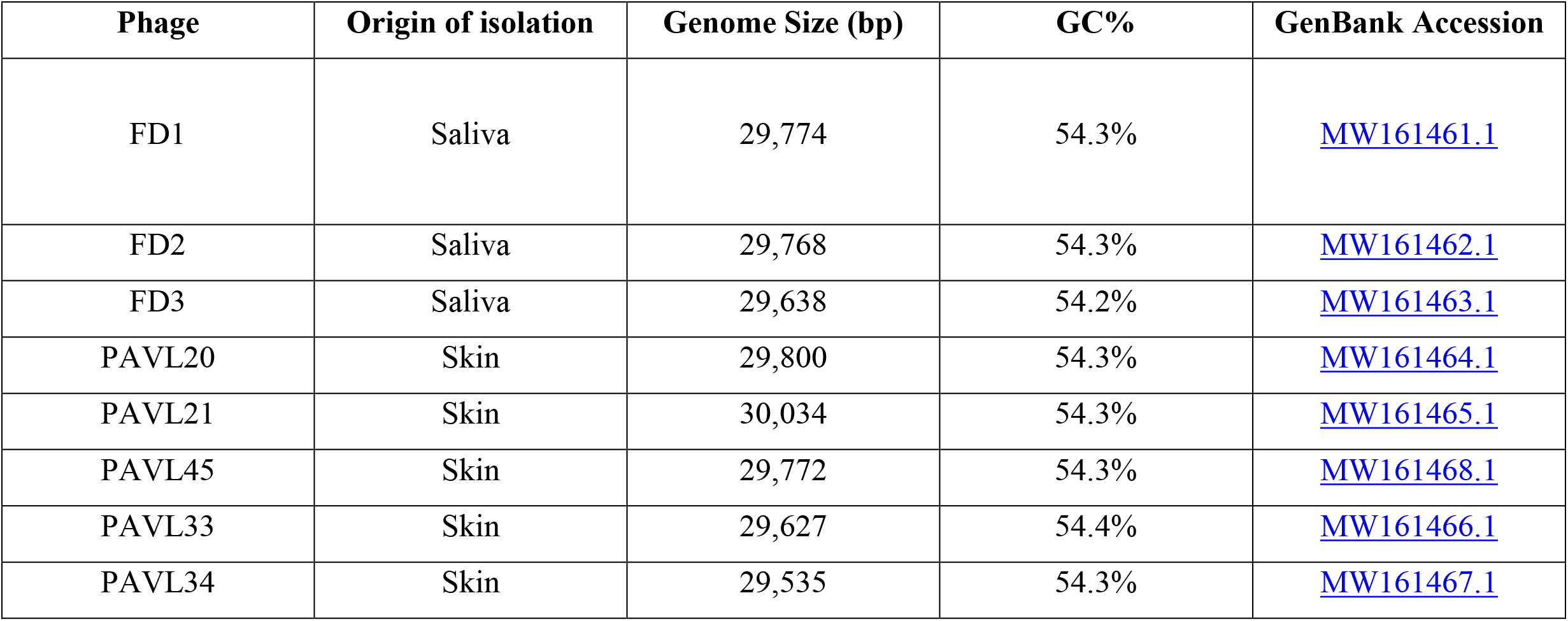
Characterization of eight novel *C. acnes* phages.

| Phage | Origin of isolation | Genome Size (bp) | GC% | GenBank Accession |
| --- | --- | --- | --- | --- |
| FD1 | Saliva | 29,774 | 54.3% | MW161461.1 |
| FD2 | Saliva | 29,768 | 54.3% | MW161462.1 |
| FD3 | Saliva | 29,638 | 54.2% | MW161463.1 |
| PAVL20 | Skin | 29,800 | 54.3% | MW161464.1 |
| PAVL21 | Skin | 30,034 | 54.3% | MW161465.1 |
| PAVL45 | Skin | 29,772 | 54.3% | MW161468.1 |
| PAVL33 | Skin | 29,627 | 54.4% | MW161466.1 |
| PAVL34 | Skin | 29,535 | 54.3% | MW161467.1 |

### Genome sequencing and analysis

The genome of each phage was fully sequenced and analyzed (Figure 1, Table 1 and Supplamental table S1). All phages were similar, with a genome size range of 29,535–30,034 bp. Based on the absence of these phage sequences in bacterial genomes and the lack of typical lysogenic genes, such as repressors, integrases, or other hallmarks of lysogens, we assumed that they have a lytic lifecycle. The absence of repeat sequences indicates that their genomes have a linear topology. Phylogeny analysis showed that they belong to the Pahexavirus genera of the Siphoviridae family. BLAST alignment of the genomes revealed a high similarity between them and the genomes of many other published *Cutibacterium* phages from the Siphoviridae family.

**Figure 1.**
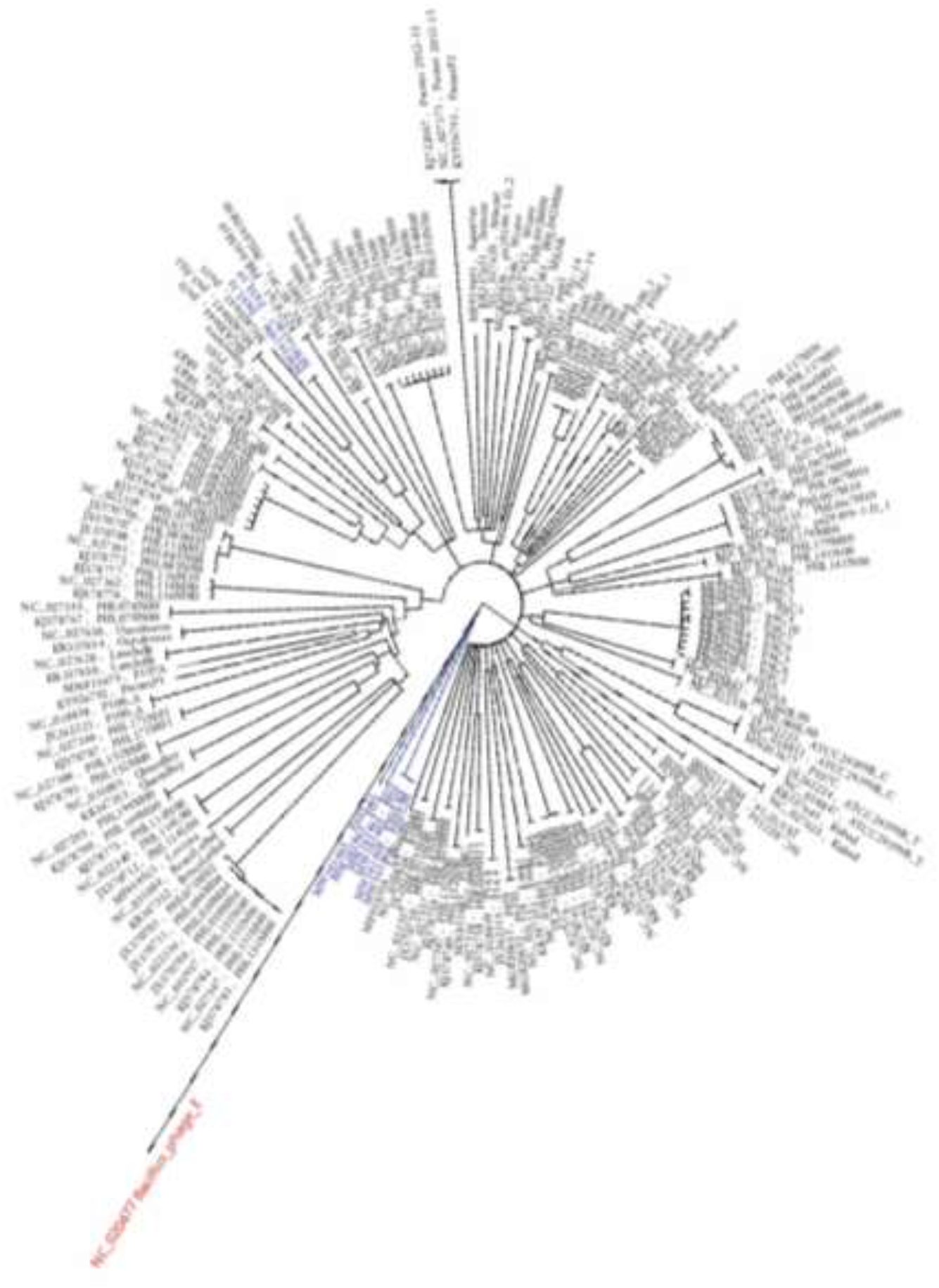
Phylogenetic tree. novel isolated *C. acnes* targeting phages (blue) in comparison to all *C. acnes* targeting phages found in blast (black). all results are relative to A *Bacillus* phage that is an out group. For more details and accession numbers see Supplamental table S1.

Even though the phages were isolated separately, they showed a very high similarity of 87%–99%, with a coverage of 95%–98%. Phages PAVL33 and PAVL34 differed in a few point mutations and short insertions/deletions, mainly hypothetical or unrecognized phage proteins, and phylogenetically they seem to differ from all other phages (Figure 1, Table 1 and Supplamental table S1). As they were isolated from different samples, we considered them to be different. However, they could also be considered variants of the same strain.

The genome of the phages was found to be free from known harmful virulence factors or antibiotic resistance elements (data not shown), indicating that they were likely safe for phage therapy.

### Phage visualization

The geometric structure and morphological characteristics of the *C. acnes* phages were visualized using transmission electron microscopy (TEM). As expected from their genome similarities, we did not observe any differences. They all had a similar capsid geometrical structure of an icosahedron, with a capsid diameter of 66 nm (Figure 2A), and a long noncontractile tail, with a length of 144 nm (Figure 2A).

**Figure 2.**
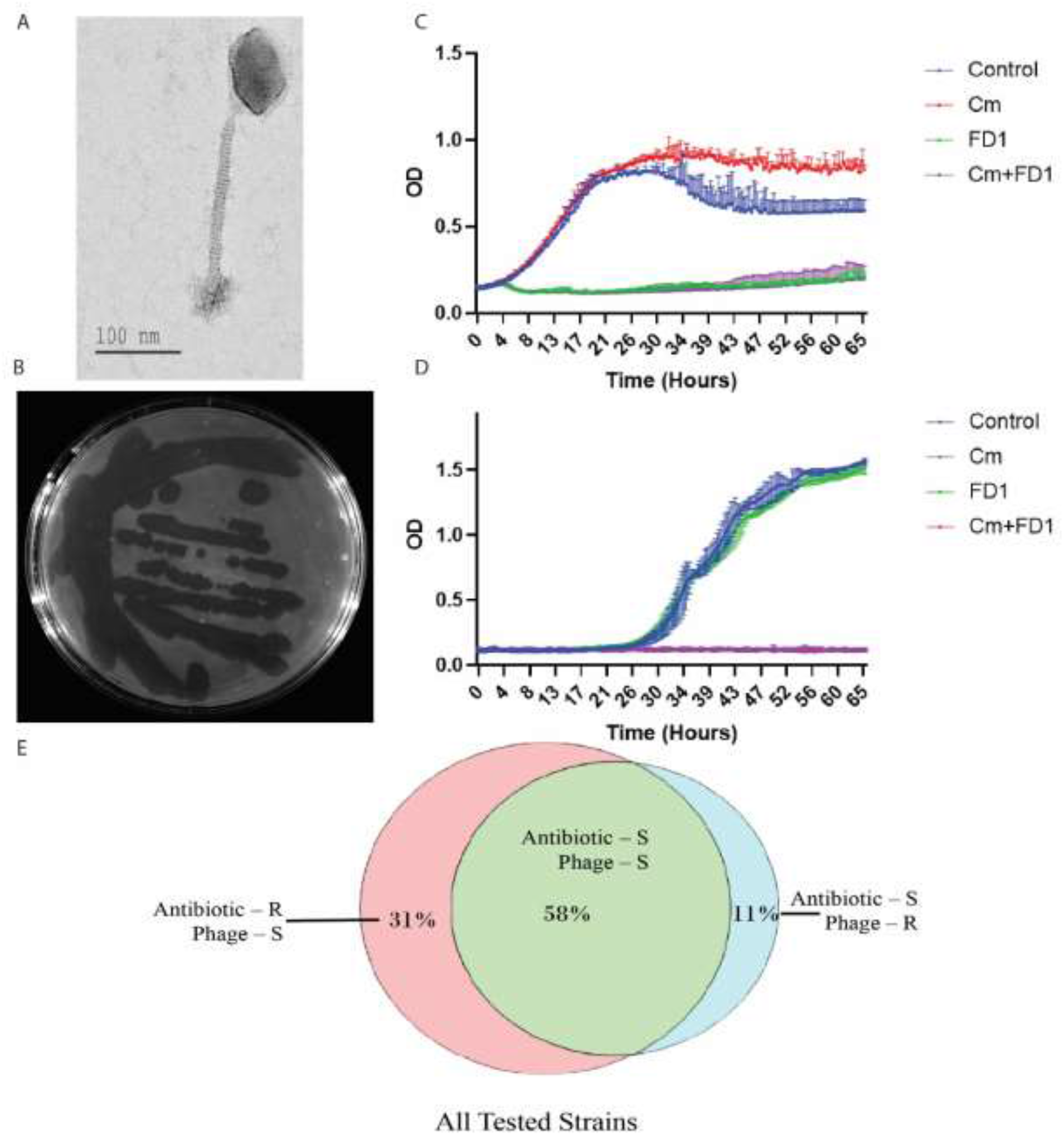
*In vitro* phage activity and coverage. (A) Transmission electron microscopy of FD1 phage. FD1 plaques on strain 27. Note the various sizes of plaques, despite the fact that they originated from a single plaque, and no lysogen was detected when the lytic phage was sequenced. (B) CM-resistant strain 28, growth with CM, FD1 phage, and their combination. The results are the average of triplicates, presented as mean ± SD. (C) FD1-resistant strain 21 growth with CM, FD1 phage, and their combination. Note that the two lower curves, CM and CM + FD1, overlap. The results are the average of triplicates, presented as mean ± SD. (E) Venn diagram of the phage and antibiotic susceptibility, S – susceptible to all tested antibiotics or phages, R – resistant.

### Host range coverage of phages *in vitro*

We tested the efficacy of the phages against *C. acnes* using solid media (agar plates) and liquid cultures (Figure 2B–D and Supllamental Figure S2). Interestingly, despite the fact that the lysate originated from a single plaque, we observed various sizes of clear plaques on the plates (Figure 2B). In the liquid culture, significant growth inhibition was observed with all phages (Figure 2C, D) in the sensitive strains (Table 2). The CM-resistant strain 28 showed improved growth with CM, but its growth was inhibited entirely by phage FD1 in the first 45 h of the experiment, followed by regrowth afterward. Notably, the combination of phages and CM achieved complete growth inhibition throughout the 65 h of the experiment (Figure 2C). Strain 21 showed the opposite effect as it was resistant to the phages but sensitive to CM or to the combination of phage and CM (Figure 2D). Thus, the combination provided 100% inhibition of all tested strains.

**Table 2.** Antibiotic and phage sensitivity of *C. acnes* strains. Erythromycin (EM), Clindamycin (CM), Tetracycline (TET), Doxycycline (DOX), and Minocycline (MC). Strains were either sensitive (S), intermediate (I), or resistant (R) to antibiotics. All strains were either sensitive (S) or resistant (R) to all eight phages tested in the phage column (P).

| # | P | EM | EM+P | CM | CM + P | TET | TET+P | DOX | DOX + P | MC | MC+P | All AB | All AB+P |
| --- | --- | --- | --- | --- | --- | --- | --- | --- | --- | --- | --- | --- | --- |
| 1 | S | S | S | S | S | S | S | S | S | S | S | S | S |
| 2 | S | R | S | R | S | I | S | R | S | I | S | S | S |
| 4 | R | S | S | S | S | S | S | S | S | S | S | S | S |
| 5 | S | R | S | R | S | R | S | R | S | S | S | S | S |
| 6 | S | S | S | S | S | S | S | S | S | S | S | S | S |
| 7 | S | R | S | I | S | I | S | I | S | S | S | S | S |
| 8 | S | S | S | S | S | S | S | S | S | S | S | S | S |
| 9 | S | S | S | S | S | S | S | S | S | S | S | S | S |
| 10 | S | S | S | S | S | S | S | S | S | R | S | S | S |
| 11 | S | S | S | S | S | S | S | S | S | S | S | S | S |
| 13 | S | S | S | S | S | I | S | S | S | S | S | S | S |
| 14 | S | R | S | R | S | R | S | R | S | R | S | R | S |
| 15 | S | S | S | S | S | S | S | S | S | S | S | S | S |
| 18 | S | S | S | S | S | S | S | S | S | S | S | S | S |
| 19 | S | S | S | S | S | S | S | I | S | S | S | S | S |
| 21 | R | S | S | S | S | S | S | S | S | S | S | S | S |
| 22 | S | S | S | S | S | S | S | S | S | S | S | S | S |
| 23 | S | S | S | S | S | S | S | S | S | R | S | S | S |
| 24 | S | R | S | I | S | S | S | S | S | S | S | S | S |
| 25 | S | R | S | I | S | I | S | R | S | I | S | I | S |
| 27 | S | S | S | S | S | S | S | S | S | S | S | S | S |
| 28 | S | R | S | R | S | I | S | R | S | S | S | S | S |
| 30 | S | S | S | S | S | S | S | S | S | S | S | S | S |
| 31 | S | R | S | R | S | I | S | R | S | I | S | I | S |
| 32 | S | S | S | S | S | S | S | S | S | S | S | S | S |
| 33 | R | S | S | S | S | S | S | S | S | S | S | S | S |
| 35 | S | R | S | R | S | R | S | R | S | R | S | R | S |
| 36 | S | S | S | S | S | S | S | S | S | S | S | S | S |
| 37 | S | S | S | S | S | S | S | S | S | S | S | S | S |
| 40 | S | S | S | S | S | S | S | S | S | S | S | S | S |
| 41 | S | S | S | S | S | S | S | S | S | S | S | S | S |
| 43 | S | S | S | S | S | S | S | S | S | S | S | S | S |
| 44 | S | S | S | S | S | S | S | S | S | S | S | S | S |
| 47 | S | S | S | S | S | S | S | S | S | S | S | S | S |
| 48 | R | S | S | S | S | S | S | S | S | S | S | S | S |
| 49 | S | S | S | S | S | S | S | S | S | S | S | S | S |

We tested the susceptibility profile of the *C. acnes* strains from our collection (8) to phages and compared it to the previously tested susceptibility to various antibiotics (8). Of the 36 strains tested, 11 were resistant to at least one antibiotic (30.6%). We found that 32 of 36 *C. acnes* isolates (88%) were phage susceptible. These 32 strains include all the above-mentioned strains resistant to at least one antibiotic (34.4%), or resistant to all antibiotics (6.3%). Moreover, the four strains which were phage-resistant were susceptible to all five tested antibiotics (Tables 2, Supplamental Table S2, and Figure 2E).

### Acne mouse model of phage therapy *in vivo*

We assessed the potential of phages for *C. acnes* infection using our isolated phages. To this end, we infected mice with *C. acnes* strain 27, a clinical isolate from a patient with severe acne vulgaris. We used FD3 as the representative phage, which showed high *in vitro* efficacy against strain 27 (Figure 2B) and was stable for a week in carbopol gel preparation (2.5%) at room temperature and at 4°C without any significant reduction of its titer (data not shown).

The infection was carried out by two intradermal injections of strain 27 on two consecutive days to the back of 34 Balb/c (ICR) mice (days one and two, respectively), followed by daily topical application of artificial human sebum (Kolar et al., 2019) to the injection site. This treatment yielded significant inflammatory acne lesions at the injection site by day 3 (Figure 3). Once inflammatory lesions were established, the mice were randomized into two groups of 17 mice each. One group was treated with FD3 phage in carbopol gel applied daily for five consecutive days, and the other group was treated with carbopol gel only. Photographs of the lesions were taken, and the lesions’ diameter, elevation/papulation of lesions, and the presence of eschar over lesions were assessed daily. A clinical score was defined to assess the severity of inflammatory lesions based on a modified revised acne Leeds grading system (O’brien et al., 1998) and other methods previously described for acne vulgaris (Agnew et al., 2016; Qin et al., 2015). Mice were sacrificed and biopsies were taken on day 10. Biopsies underwent histopathological assessment, and were analyzed for the presence of bacteria and phages at the lesion site by polymerase chain reaction (PCR), colony-forming unit (CFU), and plaque-forming unit (PFU) determination in a homogenized tissue biopsy, respectively. For more details, refer to the Assessment of Phage Clinical Efficacy section in Materials and Methods and Figure 7.

**Figure 3.**
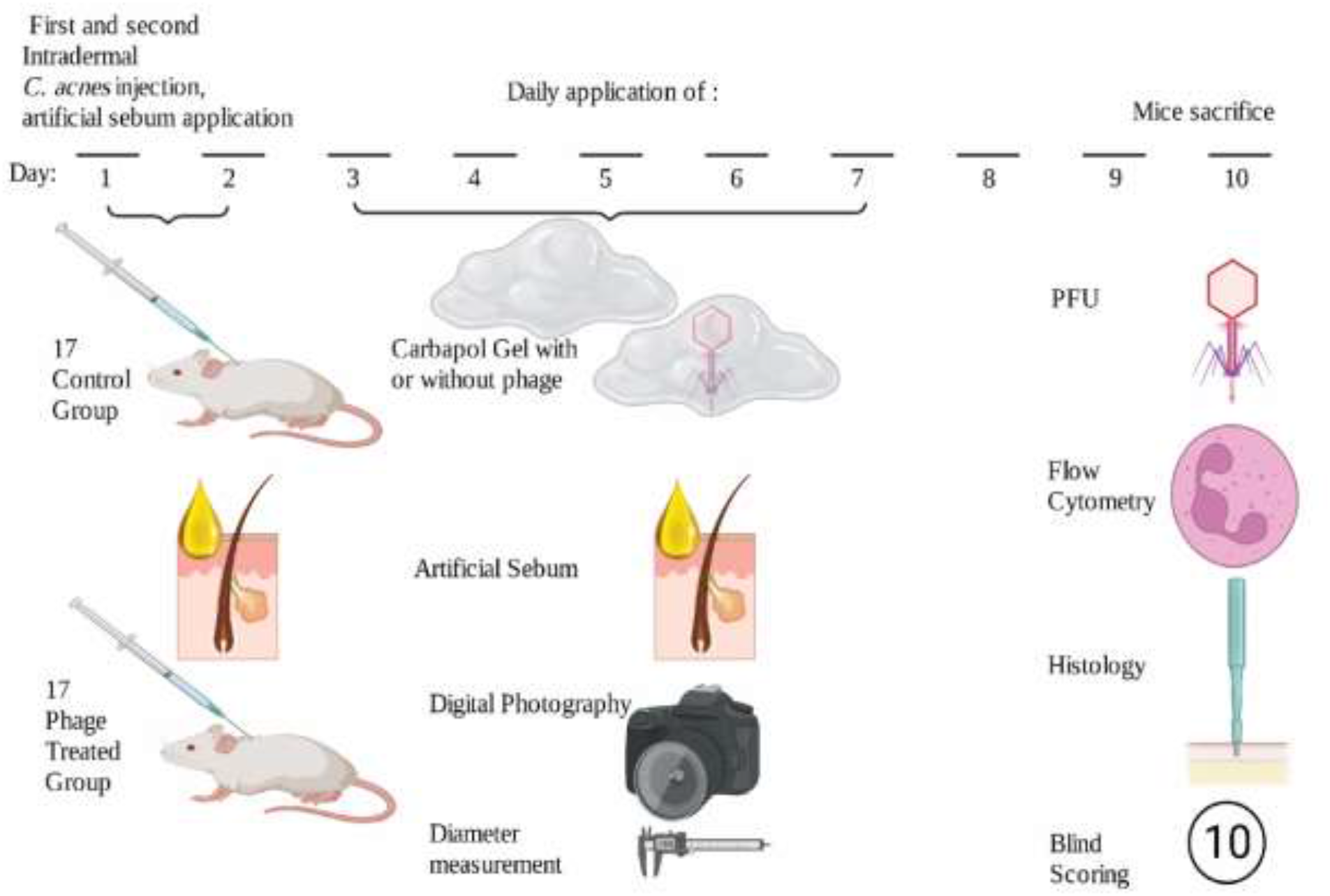
Schematic description of phage therapy *in vivo* in an acne mouse model. Mice were injected intradermally on days one and two. Throughout days 3–7, mice were treated with topical carbopol gel for the control group and phage-containing carbopol gel for the phage-treated group. Artificial sebum was applied daily starting from day 1. Mice were sacrificed on day 10 and analyzed as described (This Figureure was made with biorender.com)

The plaques of FD3 were isolated from a skin biopsy taken on day 10, three days after the last administration of the 2.5% phage-containing carbopol gel.

Mice treated with phage and mice treated with vehicle did not develop any adverse events during this experiment. The daily application of FD3 for five consecutive days resulted in improvement in three clinical parameters (diameter of inflammatory lesions, papulation/elevation of lesions, and presence and severity of eschar over lesions) compared to mice treated with vehicle only (Figure 4A - F and Supplamental Table S3).

**Figure 4.**
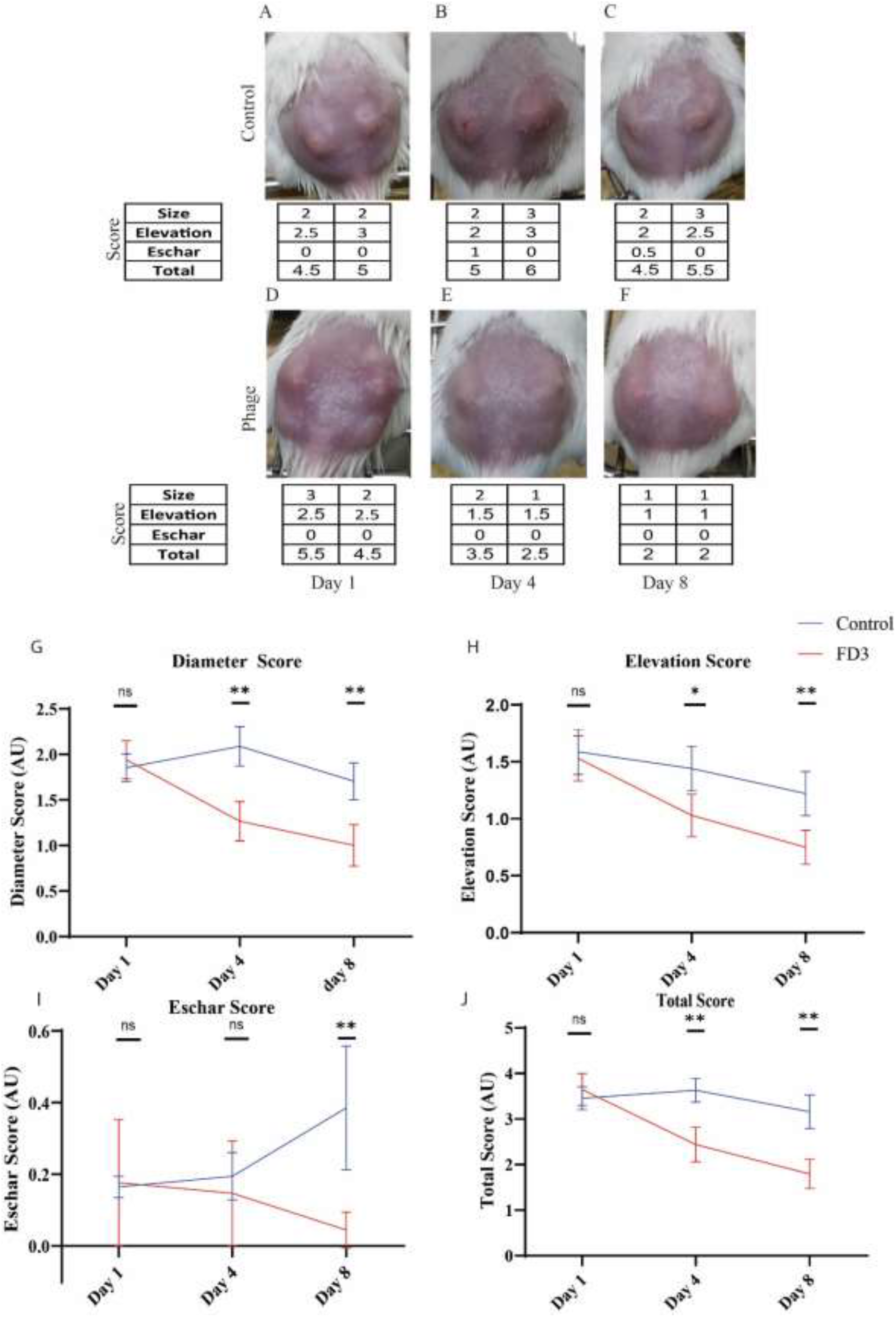
Clinical scoring of inflammatory lesions. Mice # 11 (from the control group) and mice # 12 (from the treated group) are shown as representative examples. (A-C) Photographs taken on days one, four, and eight and the clinical scores are presented. The diameter of the lesions, degree of elevation (papulation), and presence and severity of eschar were evaluated using a scoring system. For more details, refer to the Materials and Methods section. Control mice lesions. (D-F) phage-treated mice lesions. (G-J) The scores of the diameter, degree of elevation, eschar, and combined scores are shown accordingly, in blue (control group) and red (phage group) at three timepoints. Data is presented as mean ± SD. Student’s *t-test* two-tailed unpaired p-values between the control group and the FD3-treated group. *p-value < 0.05, **p-value < 0.001. Eschar score is corrected to the same value at day 1. For details on all mice see Supplemental Table S3.

The differences in the lesion diameters were significant starting from day four, with a score of 2.09 arbitrary units (AU) (5.2 mm) in the control group versus 1.27 (3.6 mm) in the treated group (p-value < 0.001, Figure 4G). The phage-treated group showed a continuous reduction in the lesion diameter, from an average score of 1.93 AU (4.6 mm on day one) to 1.00 AU (2.9 mm on day eight) (p-value < 0.001, Figure 4G). Conversely, the control group’s average lesion diameter changed in a non-continuous manner, increasing from an average score of 1.82 (4.5 mm on day one) to 2.09 AU (5.2 mm on day four) in the early days of the experiment, followed by a decrease to 1.7 AU (4.5 mm on day eight) at the end of the experiment (Figure 4G). A spontaneous reduction was observed in the control group toward the end of the experiment (Figure 4G).

Another clinical parameter assessed was the degree of elevation (papulation) of the inflammatory lesions (Figure 7). Significant differences were observed between the treated and control groups on the latter days of the experiment. The lesions’ average elevation score on day zero was 1.57 AU (average of 34 lesions) in control group and 1.52 AU in phage-treated group (non-significant). However, it decreased to 0.75 AU in the treated group and was 1.2 AU on day 8 in the control group (p-value < 0.001, Figure 4H).

The third clinical parameter was the presence of eschar in the inflammatory lesions (Figure 7). The number and severity of eschar in mice increased significantly in the control group, whereas the average eschar score in the phage-treated mice was lower (p-value < 0.01, Figure 4I). The comparison of the combined scores of the three clinical parameters between the two groups (Figure 7) was significant on days 4–8 (p-value < 0.001 at both timepoints, Figure 4J).

### Histopathological evaluation of inflammatory lesions

Histopathological evaluation was performed on the biopsies taken from both groups’ inflammatory lesions on day eight of the experiment (Figure 5).

**5.**
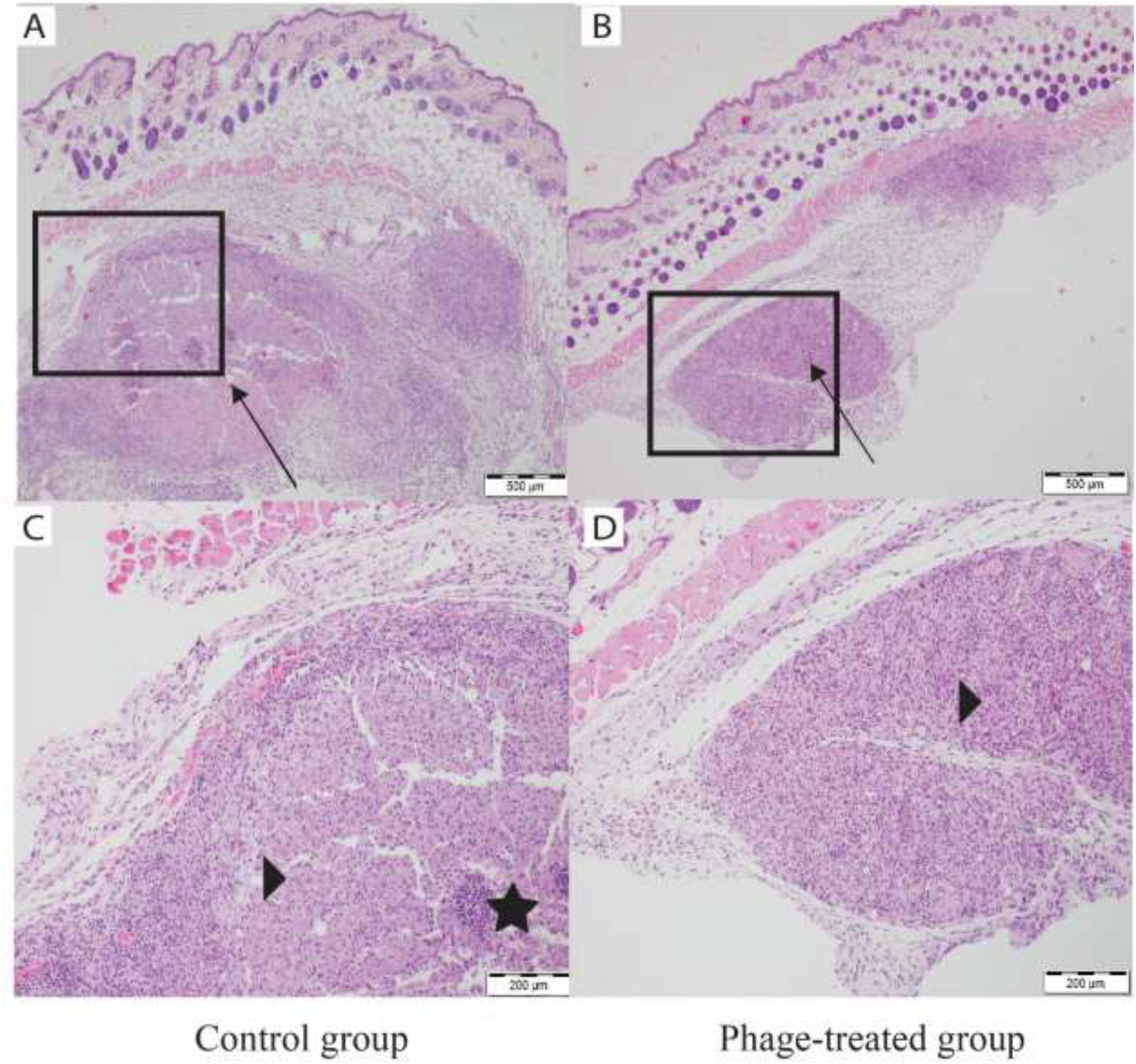
Histology of inflammatory lesions. (A) Nodular inflammation (arrow) involving the dermis and subcutaneous tissue at 4× magnification in the control group. (B) nodular inflammation (arrow) involving only the subcutaneous tissue below the panniculus carnosus at 4× magnification in the phage-treated group. (C) A nodular infiltrate with an area of necrosis with small numbers of interspersed neutrophils (stars) surrounded by macrophages (arrowhead) and rare lymphocytes at 10× magnification in the control group. (D) A nodular infiltrate of macrophages (arrowhead), many with vacuolated cytoplasm mixed with a minimal number of neutrophils at 10× magnification in the phage-treated group.

The skin of the control mice showed evidence of a more severe and acute pyogranulomatous inflammatory process. In some cases, the inflammation involved the dermis and subcutaneous tissue (Figure 5A). A nodular infiltrate, with an area of necrosis with small numbers of interspersed neutrophils surrounded by macrophages and rare lymphocytes, was observed (Figure 5C). By contrast, inflammation was less severe in the phage-treated group, with mostly chronic granulomatous infiltration, involving only subcutaneous tissue below the panniculus carnosus mixed with minimal numbers of neutrophils (Figure 5B, D).

### Evaluation of neutrophilic inflammatory reaction

Three mice were sacrificed at different time points: day three (after establishing inflammatory lesions, before commencing treatment with carbopol or phage), day five, and day seven of the treatment with carbopol alone or phage. Three uninfected mice were sacrificed as the negative control (uninfected). Skin specimens underwent evaluation by flow cytometry)Materials and Methods). Antibodies for Ly6G, CD64, CD11b, and CD45 were used to identify different cell populations. As acne is a neutrophil-driven process, we decided to specifically examine the neutrophilic infiltrate in the specimens collected using the neutrophilic marker Ly6G + (Swamydas et al., 2015). On day three, the average neutrophil percentage (%NT) was 48.63% in *C. acnes*-injected mice (see a representative example in Figure 6A). The skin specimen of the control unifected mice had 11.21%NT (See representative example in Figure 6B,). On day five, the average %NT was 40.6% in the infected untreated control group and 25.12% in the infected phage-treated group. On day seven of the experiment, the average %NT was 35.22% in the infected untreated control group and 17.6% in the infected phage-treated group. Bacteria-injected mice on day three, before the group allocation were significantly different from the phage-treated group on day seven (p-value = 0.02), and the placebo-treated group was different from the phage-treated group (p-value = 0.02) (Figure 6C). The lesions of phage-treated mice had normalized to the uninfected control on days five and seven (p-value = 0.22, 0.28 accordingly), whereas the control-treated group did not normalize (p-value < 0.05) (Figure 6D).

**Figure 6.**
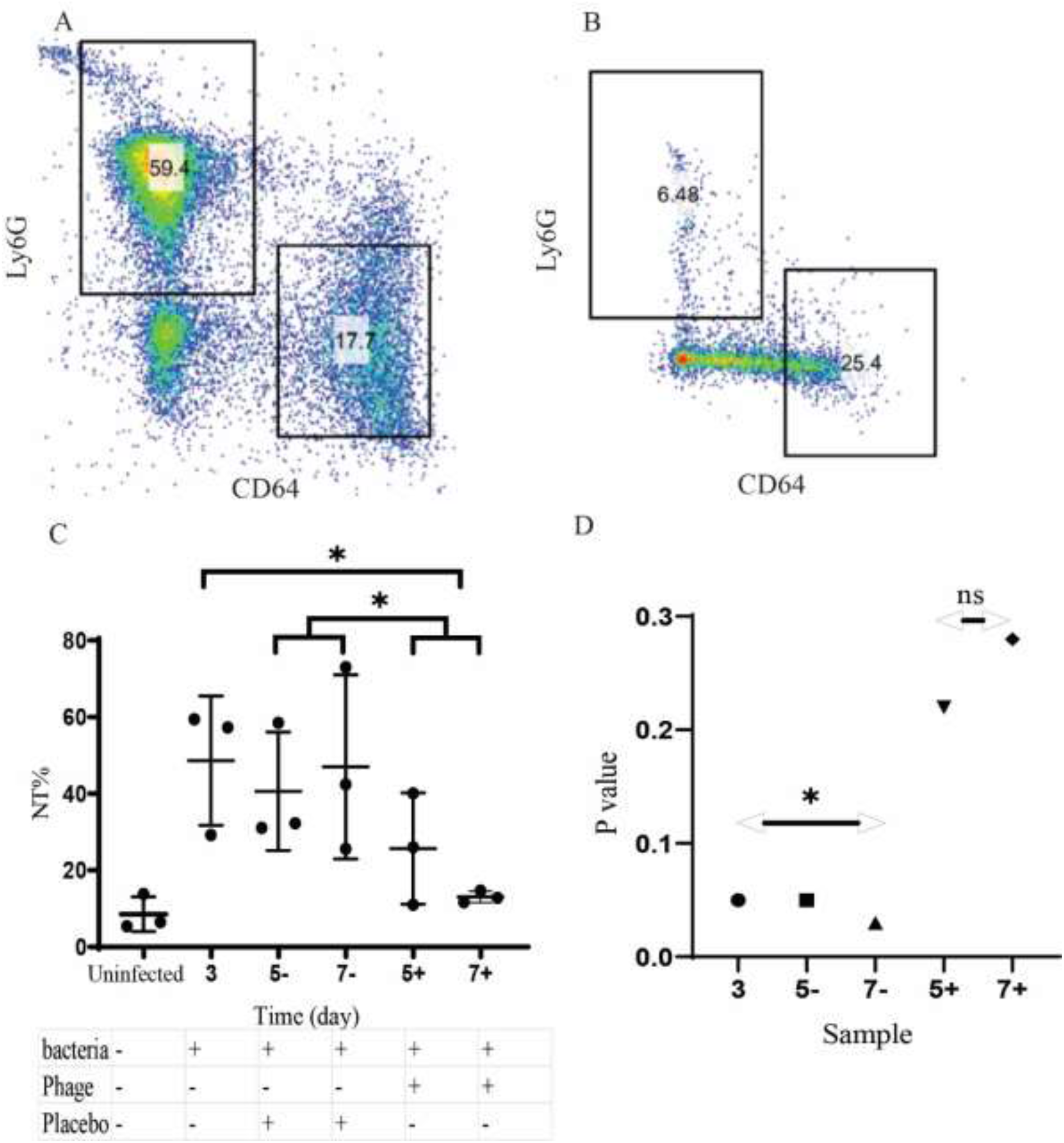
Flow cytometry at various timepoints. (A) *C. acnes* infected mice on day three of the experiment, 59% of the myeloid cells (%NT) (CD64 Negative, Ly6 G positive). (B) uninfected control, 6%NT. (C) Flow cytometry analysis. A significant difference was found between day three and day seven in the phage group and between the phage group and the control group. Data is presented as mean ± SD. (D) The p-value of each group compared to the uninfected group. The Mann–Whitney U test was used. **\*** denotes p-value < 0.05 and **ns** “not significant”.

Monocytes were also examined (CD64+) (Genel et al., 2012), without significant dynamics in this model.

## DISCUSSION

This study presents a phage-based topical treatment in an acne mouse model. The results provide *in vivo* support for the efficacy and safety of transdermal phage therapy in treating acne vulgaris and shed light on the cytological modification involved in the response to phage therapy. Antibiotic resistance of *C. acnes* has been reported worldwide. Regional differences and dynamic changes over time are remarkable, and an increase in resistance rates has been reported (Coates et al., 2002; Karadag et al., 2020; Sheffer-Levi et al., 2020). Studies on phage susceptibility have also been performed (Brüggemann and Lood, 2013; Castillo et al., 2019; Jong et al., 1975; Liu et al., 2015; Marinelli et al., 2012), demonstrating an average of 88% susceptibility to *C. acnes*, consistent with our findings of an 88.8% susceptibility rate (Brüggemann and Lood, 2013; Castillo et al., 2019).

Nevertheless, to the best of our knowledge, the current study is the first to demonstrate full treatment coverage of *C. acnes* strains using a combination of antibiotics and phages (Figure 2E). As previously shown on *Acinetobacter baumannii* (28), this may indicate that resistance to one modality of treatment interferes with resistance to the other. Further research is needed to clarify whether resistance mechanisms induce the re-sensitization of the other.

Among the *C. acnes* strains resistant to a given antibiotic, the phage–antibiotic combination caused effective inhibition, indicating that phage therapy could serve as an optional treatment for antibiotic-resistant acne. In all investigated strains, whether they were resistant to a specific antibiotic or phage, a combination of phage and antibiotic showed a non-inferior response to treatment with antibiotics alone. This shows that a combination of phages and any given antibiotic may be an effective empiric regimen (Figure 2, Table 2) (Katsambas et al., 2004; Zaenglein et al., 2016).

Therefore, we suggest that phage therapy should be evaluated for the treatment of acne vulgaris, especially in antibiotic-resistant cases, perhaps in combination with conventional therapy. This approach may give superior results *in vivo* and lead to a decrease in the development of antibiotic-resistant strains. *In vivo* and clinical data are needed to confirm this assumption.

*C. acnes* strains showed an “all or none” susceptibility to all phages; bacterial strains that were resistant to a single phage, with no exceptions, showed resistance to all eight phages. Bacteria that showed susceptibility to one phage showed susceptibility to all phages (Table 2 and Supplamental Table S2).

*C. acnes* phages are known to demonstrate small variability, to which most isolated *C. acnes* strains are susceptible, with little genomic variance (Castillo et al., 2019; Marinelli et al., 2012). The two possible theories explaining the small variety are as follows: 1) small niche theory: the fact that the life niche of *C. acnes* is only in the pilosebaceous unit makes it difficult for phages to go through different varieties of bacteria (Castillo et al., 2019); 2) bottleneck theory: evolutionary limitations have made it possible for only one family of phages to survive (Castillo et al., 2019). Phage isolation from saliva reflects the presence of *C. acnes* phages and bacteria in this environment, tipping the scale away from the “small niche theory.” The isolated phages show limited evolutionary space and are all part of the same phylotype (Abedon, 2009). One possible theory of an evolutionary force involves prokaryotic innate immunity in the form of repeating palindromic nucleotides that correlate with resistance to several phages (Marinelli et al., 2012). For example, unlike *C. acnes* phages, *Propionibacterium freudenreichii* phages are more diverse (Cheng et al., 2018). Perhaps the evolutionary forces differ between species. The latter is not part of the human microbiome and is an important part of the cheese manufacturing process and a possible probiotic (Thierry et al., 2011).

Moreover, the high genomic similarity of our *C. acnes* phages, together with the high similarity observed in other known *C. acnes* phages, shows that a more precise taxonomic method is required to determine whether the given phage is new. One option is to compare only the specific gene/s and not entire genomes.

Previous studies have assessed the activity and efficacy of injected acne phages in an acne mouse model (Kim et al., 2019; Lam et al., 2021; Nelson et al., 2012). Such a method of phage delivery is not clinically applicable. Our work is the first *in vivo* demonstration of the efficacy and safety of topically applied phages in an acne mouse model, providing further support for the potential role of phage therapy in acne vulgaris.

The establishment of an acne mouse model is not trivial, as acne is solely a human disease (Plewig et al., 2019). The use of artificial sebum and two consecutive injections of a clinically isolated *C. acnes* strain enabled us to induce inflammatory lesions, simulating inflammatory papules and nodules of acne vulgaris (Figure 4 and Supplamental Table S3). Our model showed a self-resolving effect, with spontaneous recovery of the inflammatory lesions over time (Figure 4 and Supplamental Table S3). Therefore, the treatment period was limited to five days, whereas any treatment for acne vulgaris in humans, including antibiotics, was given for weeks and months (Zaenglein et al., 2016). Despite the concise treatment period provided in our acne mouse model, we observed significant and fast improvement in the inflammatory lesions in the FD3-treated group compared to the control group (Figures. 4-6).

The significant earlier and faster clinical improvement in the FD3-treated group suggests a positive effect of FD3, which is further supported by evidence of reduced neutrophilic infiltration, both in histopathology, and flow cytometry analysis. Neutrophil-mediated inflammation is an essential part of acne vulgaris pathogenesis. The described reduced percentage of neutrophils in the lesions could be due to fewer bacteria in the tissue. Another possible mechanism is phage–innate immunity interactions, which have been described mainly *in vitro* (Van Belleghem et al., 2019) (Figures. 5, 6). Further testing of immunological *in vivo* data is needed to assess this hypothesis. Based on our results, early in the course of the inflammatory lesions, there seemed to be a dominant neutrophilic infiltration, but phage therapy induced faster neutrophilic clearance or inhibited neutrophilic migration later in the course of the lesions. Moreover, bacterial infected phage-treated mice were not significantly different from the uninfected control skin, whereas the bacterial infected untreated control mice were significantly different from the negative control.

Although one of our aims was to show the efficacy of topical application of the phage, we were concerned that the phage might not penetrate the skin. However, we were able to show phage penetration by isolating the phages from the lesions three days post-administration. Therefore, we assume that phages penetrate the lesion and multiply inside it, using the target bacterium. In this study we did not examine the efficacy of a combination of phages and antibiotics in our mouse model. The isolated strain used in this model was not antibiotic resistant, and we focused on investigating phage efficacy alone. Phages that have been proven efficient should be assessed for synergism with antibiotics in another mouse model and in clinical trials.

According to the collected data and in light of the harmless expected effect of phages on the skin microbiome (Barnard et al., 2016), we hypothesize that phage therapy is a promising treatment modality for acne vulgaris. Nevertheless, this should be further tested in clinical trials. If anti-*C. acnes* phage treatment is found to be safe and effective in humans, it is expected to be used in the future to treat acne vulgaris and reduce the widespread use of antibiotics in this common skin condition.

## MATERIALS AND METHODS

### Bacterial clinical isolates

This study used a collection of 36 *C. acnes* clinical isolates with various ribotype single-locus sequence typing (SLST) types (Figure S1) obtained from the Department of Clinical Microbiology and Infectious Diseases of Hadassah Medical Center, as we previously described (Sheffer-Levi et al., 2020). Unless mentioned otherwise, the *C. acne*s isolates were grown in Wilkins–Chalgren broth (Difco, Sparks, MD, USA) at 37°C under anaerobic conditions and stored at −80°C in glycerol (25%) until use. Bacterial concentrations were evaluated using 10 µl of 10-fold serial dilutions plated on Wilkins agar plates under anaerobic conditions. Colonies were counted after 48 h at 37°C, and the number of CFU/mL was calculated.

### Antibiotic susceptibility

Following bacterial identification, *C. acnes* isolates were sub-cultured in Wilkins broth (Oxoid, Basingstoke, UK) and suspended at a density of 1.0 McFarland. Bacterial lawns were prepared on anaerobic blood plates (Novamed, Jerusalem, Israel) and dried. Antibiotic susceptibility was assessed by determining a minimal inhibitory concentration (MIC) using an epsilometer test (ETEST® bioMérieux, St. Louis, MO, USA). The MIC was determined following 48 h of incubation under anaerobic conditions as the point on the scale at which the ellipse of growth inhibition intercepts the plastic strip. The antibiotics used tetracycline (TET), doxycycline (DOX), minocycline (MC), erythromycin (EM), and clindamycin (CM),. The breakpoints used to define susceptibility or resistance to CM and TET followed the recommendations set by the Clinical and Laboratory Standards Institute (Wayne, 2017). Resistance to CM was defined at an MIC above 2 µg/ml and to TET at an MIC above 4 µg/ml. As no standards exist for the breakpoints of EM, DOX, and MC, those with an MIC of ≥ 0.5 µg/ml (for EM) and ≥ 1 µg/ml (for DOX and MC) were defined as resistant according to the definitions used in previous studies (Toyne et al., 2012). CM (1.5 ug/ml) and EM (15 ug/ml) were the antibiotics used when assessing phage antibiotic synergism.

### Phage isolation and propagation

The phages were isolated using the standard double-layered agar method, as previously described (Adams, 1959). Briefly, 1–5 ml of saliva or skin samples were mixed with 5 ml of phage buffer (150 mM NaCl, 40 mM Tris-Cl, pH 7.4, 10 mM MgSO4) centrifuged on the following day (centrifuge 5430R, rotor FA-45-24-11HS; Eppendorf, Hamburg, Germany) at 10,000 g for 10 min. The supernatant was filtered first through filters with a 0.45-μm pore size (Merck Millipore, Ltd., Ireland) and then through filters with a 0.22-μm pore size (Merck Millipore Ltd., Ireland). Exponentially grown bacterial cultures were inoculated with filtered skin or saliva effluent for 24 h at 37°C in an anaerobic jar. The cultures were filtered again and added to 5 ml of Wilkins– Chalgren agar (Oxoid Basingstoke, United Kingdom) containing 0.5 ml of overnight-grown *C. acnes* after centrifugation. The supernatant was filtered as described and plated using soft agar (0.6%) overlaid with the test strain and then incubated overnight at 37°C in an anaerobic jar, as described above. Clear plaques were observed and transferred into a broth tube using a sterile Pasteur pipette. The phage stocks were inoculated with bacterial cultures to collect high titer lysates, which were then stored in Wilkins at 4°C.

### Determination of phage concentration

The phage solution was inoculated into 5 ml of pre-warmed Wilkins soft agar (0.6%). A 0.1 ml portion of an overnight culture of *C. acnes* was added to the tube and placed on a Wilkins agar plate. The plates were incubated anaerobically for 48 h, and the appearance of plaques on the plates was used to determine whether the bacteria were phage-susceptible. The phage concentration was determined according to the standard PFU method. Lysates were serially diluted 10-fold into 5 ml of pre-warmed Wilkins soft agar (0.6%). A 0.1 ml portion of an overnight culture of *C. acnes* was added to the tube, placed on a Wilkins agar plate, and grown anaerobically for 48 h. The number of plaques was counted, and the initial concentration of PFU/mL was calculated (Adams, 1959). If not specified otherwise, phages were grown to an initial concentration of 10^8^PFU/ml.

### Growth kinetics

Logarithmic (10^7^ CFU/ml) *C. acnes* cultures with phages and antibiotic concentrations, as described, were put in a total volume of 200 µl in triplicates. The growth kinetics of the cultures were recorded at 37°C anaerobic conditions sealing the conetents of an anerobic bag (Thermo Fisher Scientific, Waltham MA, USA) on the outer cells of a 96 well plate, with 5 s shaking every 20 min in a 96-well plate reader (Synergy; BioTek, Winooski, VT) at 600 nm. The mean and 95% confidence intervals are shown.

### Genome sequencing and analysis

The DNA of the phages was extracted using a phage DNA isolation kit (Norgen Biotek, Thorold, Canada) (Summer, 2009). Libraries were prepared using an Illumina Nextera XT DNA kit (San Diego, CA, USA). Normalization, pooling, and tagging were performed using a flow cell with 1 × 150 bp paired-end reads, which were used for sequencing with the Illumina NextSeq 500 platform. Sequencing was performed in the sequencing unit of the Hebrew University of Jerusalem at the Hadassah Campus. Trimming, quality control, reads assembly, and analyses were performed using Geneious Prime 2021.2.2 and its plugins (https://www.geneious.com). Assembly was performed using the SPAdes plugin of Geneious Prime. Annotation was performed using RAST version 2 (https://rast.nmpdr.org/rast.cgi, accessed on March 1, 2021), PHAge Search Tool Enhanced Release (PHASTER) (https://phaster.ca, accessed on March 1, 2021), and the BLAST server. The phages were scanned for resistance genes and virulence factors using ABRicate (Seemann T, Abricate, https://github.com/tseemann/abricate) based on several databases: NCBI, CARD, Resfinder, ARG-ANNOT, EcOH, MEGARES, PlasmidFinder, Ecoli_VF, and VFDB.

### TEM

A lysate sample (1 ml) with 10^8^ *PFU*/*ml* was centrifuged at 20,000 g (centrifuge 5430R, rotor FA-45-24-11HS; Eppendorf, Hamburg Germany) for 2 h at room temperature. The supernatant was discarded, and the pellet was resuspended in 200 µl of 5 Mm MgSO_4_. This sample was spotted on a carbon-coated copper grid, with an addition of 2% uranyl acetate and incubated for 1 min; the excess was removed (Yazdi et al., 2020). The sample was visualized using a transmission electron microscope (TEM 1400 plus Joel, Tokyo, Japan), and a charge-coupled device camera (Gatan Orius 600) was used to capture images in the microscopy department of the intradepartmental unit of Hebrew University.

### Acne vulgaris mouse model

For the induction of acne vulgaris lesions, we used the clinically highly virulent *C. acnes* strain S.27 (Sheffer-Levi et al., 2020). Bacteria were grown for three days and brought to a concentration of 10^9^ *CFU*/*ml*, and 50 µl of bacteria were injected intradermally into the right and left sides of the back of 34 ICR eight-week-old albino mice, for a total of 68 lesions (Kolar et al., 2019). A second intradermal bacterial infection was performed again after 24 h in the same region, as the lesions were not sufficient after one injection. Based on a previous report showing that synthetic sebum enhances inflammatory acne vulgaris lesions in an acne vulgaris mouse model (Kolar et al., 2019), 20 µlof artificial sebum (prepared by mixing 17% fatty acid, oleic acid, triolein, 25% jojoba oil, and 13% squalene) was applied immediately, following the first intradermal injection, and reapplied daily for the duration of the experiment. Mice were sacrificed three days post-last gel administration. Chlorhexidine cleaning of the lesion and three days without treatment assured that the phages isolated *ex vivo* were not the phages recently applied and not absorbed. For flow cytometry analysis, 15 mice were sacrificed at three timepoints. Three mice on day three before receiving any treatment. On day five, three phage-treated mice and three control-treated mice were sacrificed; on day seven, three phage-treated mice and three control-treated mice were sacrificed. From several mice, uninfected skin was taken as a negative control.

### Topical phage treatment

Mice were randomly assigned to two groups of 17 mice, with each group treated daily for five days. The control group received a topical 2.5% carbopol gel (Super-Pharm, Herzliya, Israel), and the treated group received a 2.5% carbopol gel containing a *C. acnes* phage, FD3, at a concertation of 10^9^ *PFU*/*Ml*. The groups were separated to prevent the transfer of phages between them.

### Assessment of phage clinical efficacy

Photography was performed daily. The degree of clinical inflammatory changes was recorded daily using three clinical parameters: 1) measurement of the lesion diameter using an electronic caliper (Winstar), 2) degree of elevation/papulation of inflammatory lesions, and 3) presence and severity of eschar within inflammatory lesions. A clinical scoring system was developed and evaluated (Figure 7) by combining the three abovementioned clinical parameters. The scoring system was developed based on various clinical acne vulgaris scoring methods (Agnew et al., 2016; Qin et al., 2015), including the revised acne Leeds grading system (O’brien et al., 1998). To avoid bias, the score was evaluated blindly by two independent investigators. The unit of the score was AU. The eschar score was corrected to the same initial score because of uneven mouse allocation of this clinical parameter.

**Figure 7.**
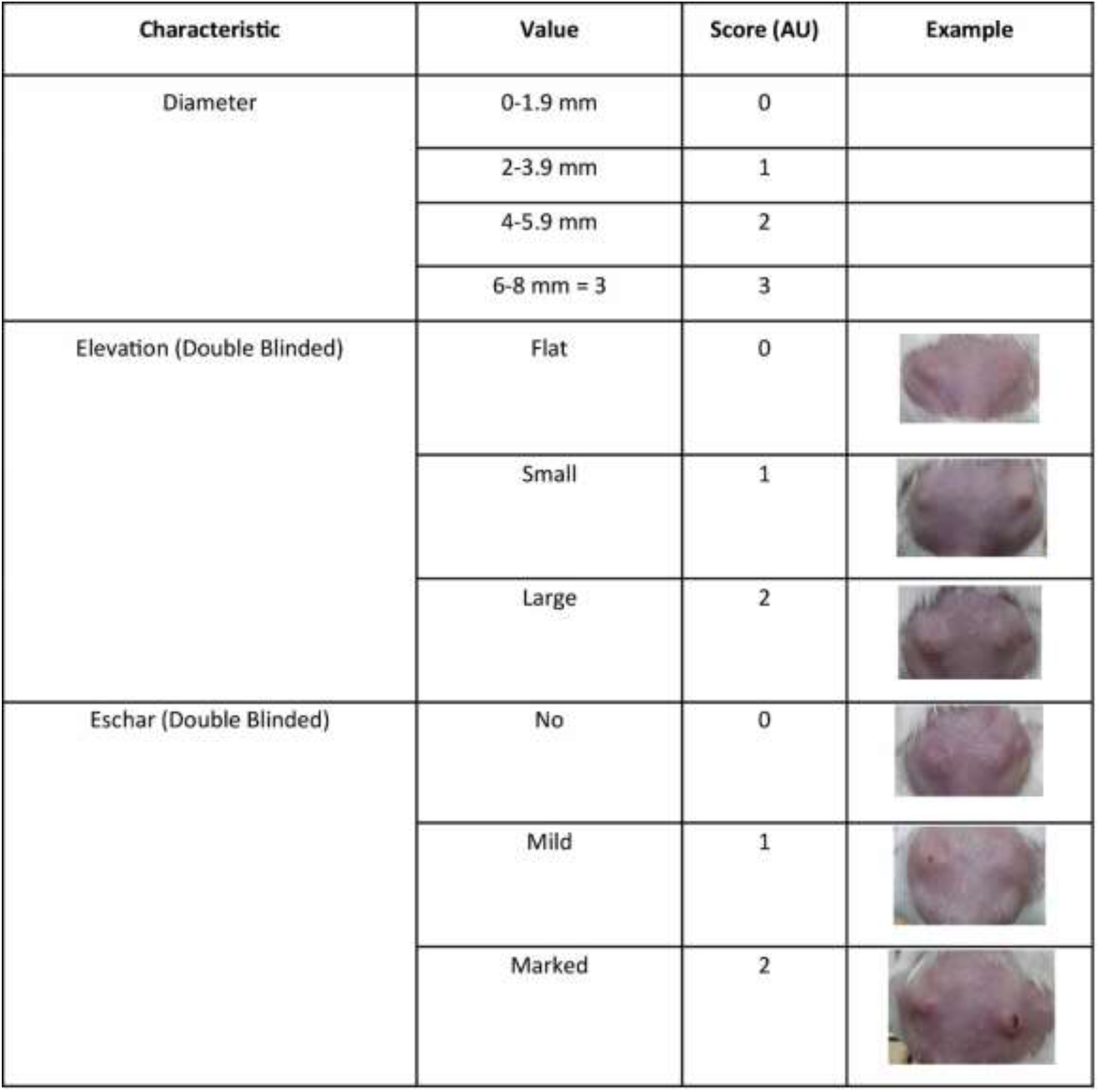
Definition of the clinical scoring of mice. The following parameters were used to score the severity of the lesions based on a modified revised acne Leeds grading system (O’brien et al., 1998) and other methods previously described for the clinical scoring of acne vulgaris (Agnew et al., 2016; Qin et al., 2015). The total score, with a range of 0–7, was calculated by summing up the parameters. Refer to the assessment of phage clinical efficacy subsection in the Materials and Methods section for more details.

### Histopathological evaluation

Punch skin biopsies of 3 mm were obtained from inflammatory lesions at the end of the treatment. Several representative biopsies were used for histopathological evaluation, and the other tissue samples were homogenized in sterile phosphate buffered saline (PBS) using stainless steel beads and a bullet blender tissue homogenizer (Next Advance) for bacterial and phage assays. The presence of bacteria at the lesion’s site was validated by PCR using the following specific primers: the SLST primer, forward primer 5’-CAGCGGCGCTGCTAAGAACTT-3’, and reverse primer 5’-CCGGCTGGCAAATGAGGCAT-3’. The presence of the phage was assessed using the specific forward primers 5’TGATGCTGTAGGTGGCTGTG-3’ and reverse primer 5’– CCGAGACGAAATGACCACCA-3’. For phage recognition, a CFU count was performed for a quantitative assessment of viable bacteria on selective media supplemented with furazolidone (0.5 μg/mL) to inhibit the growth of staphylococci and live phages using a PFU count on the *C. acnes* bacterial lawn.

Pathological skin evaluation of *C. acnes*-injected lesions was performed using tissue biopsies soaked in formalin for fixation, trimmed, embedded in paraffin, and sectioned, followed by staining with hematoxylin and eosin. The histological processing of the biopsies, including embedding, sectioning of tissues, and preparation of slides, was performed in the Histology Laboratory of the Animal Facilities of the Faculty of Medicine of Hebrew University.

### Flow cytometry

Extraction of immune cells from skin biopsies was performed using the protocol of Lou *et al*. (Lou et al., 2020), with some modifications to the Dispase II concentration and incubation time. Briefly, 1 cm × 1 cm of skin was taken from the inflammatory lesions of sacrificed mice. The skin was washed in Hanks’ Balanced Salt Solution (HBSS) three times, cut into four pieces diagonally, and incubated with 8 mg/ml Dispase II (Merck, Kenilworth, New Jersey) for 12 h. Following dermis and epidermis separation, the dermis was cut and put in 3.5 ml of Dermis Dissociation Buffer composed of 100 μg/mL of DNASE I (Merck) and 1 mg/ml collagenase P (Merck, Kenilworth New Jersey) in Dulbecco’s Modified Eagle Medium (DMEM/high glucose) (Merck, Kenilworth New Jersey, USA) for 1 h. Suspensions were passed through a 40-µm strainer into a 50 ml tube and rinsed again in 12 ml of DMEM with 10% Fetal bovine serum (FBS) (Merck, Kenilworth New Jersey) in 15 ml tubes, centrifuged at 400 g for 5 min at 4°C, supplied with 2 ml of staining buffer (Lou et al., 2020) composed of PBS with 2% Fetal calf serum (FBS (Merck, Kenilworth, New Jersey, USA), and fixated with 250 µlBD Cytofix™ (BD Biosciences, Franklin Lakes, New Jersey, USA).

Single-cell suspensions were incubated and labeled with the following antibodies obtained from BioLegend (San Diego, CA, USA) at a 1:100 dilution: CD115 (AFS98), CD45 (30-f11), CD64 (X54-5/7.1), CD11b (M1/70), LY6C (HK1.4), LY6G (1A8), and Zombie UV™ for dead cell exclusion. Following membrane staining, the cells were fixed using a Fixation/Permeabilization Solution Kit (BD) according to the manufacturer’s instructions. Flow cytometry was performed using Cytek® Aurora (Cytek, Fremont, CA, USA), and data were analyzed offline using FlowJo 10.7.2 (BD Biosciences, Franklin Lakes, New Jersey, USA).

### Statistical analysis

GraphPad Prism 8.0.2 (42) was used to perform the statistical analysis and graph formation. Significance was calculated using the Student’s t-test two-tailed unpaired *p-values* and the Mann– Whitney U-test (significance level: p < 0.05). Pearson r and r square regression models were used. The results were the mean of at least three independent experiments.

### Study approval

This study was approved by the Authority for Biological and Biomedical Models at our institution (Approval number: MD1815519\3).

## Acknowledgments

We would like to thank Simon Yona for advising with flow cytometry, and suppling the materials for flow analysis. We would also like to thank the Core Research Facility of Hebrew University, Ein Karem Campus, Abed Nasereddin and Idit Shiff for the deep sequencing, and Eduardo Berenshtein for the TEM.

We are grateful to Mariana Scherem for helping us prepare our *in vivo* experiments.

## Funding

United States–Israel Binational Science Foundation Grant #2017123

Israel Science Foundation IPMP Grant #ISF1349/20

Rosetrees Trust Grant A2232

Milgrom Family Support Program

Gishur Fund of Hadassah Medical Center

George and Linda Hiltzick’s donation to Hadassah Medical Center

## Authors’ contributions

**Conceptualization**: AR, CR, VMP, RH

**Methodology**: AR, CR, SCG, VMP RH

**Experiment and analysis**: AR, CR, VL, SSL, SAO, TS, LS, RL

**Funding**: VMP, RH

**Project administration**: SCG, VMP, RH

**Supervision**: SCG, VMP, RH

**Writing (original draft)**: AR, CR, SAO, VMP, RH

**Writing (review and editing)**: SCG

## Data and materials availability

All data, codes, and materials used in the analysis must be available in some form to any researcher for the purpose of reproducing or extending the analysis. Include a note explaining any restrictions on materials, such as material transfer agreements (MTAs). Note accession numbers to any data related to the paper and deposited in a public database; include a brief description of the data set or model with the number. If all data are in the paper and supplementary materials, include the sentence, “All data are available in the main text or the supplementary materials.”

