## Supplemental Files for "Towards phage therapy for acne vulgaris: Topical application in a mouse model"

**Supplementary Materials**

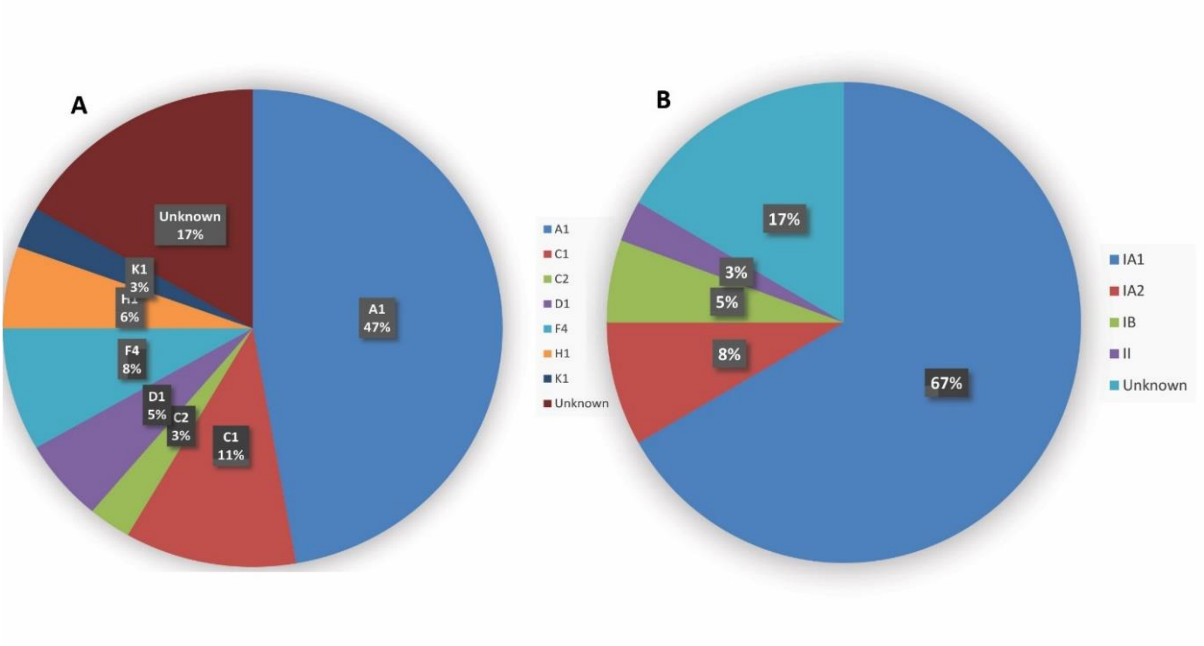


**Figure S1.** *C. acnes* molecular typing in 36 clinical isolates

SLST types (A) Phylotypes (traditional typing) (B)

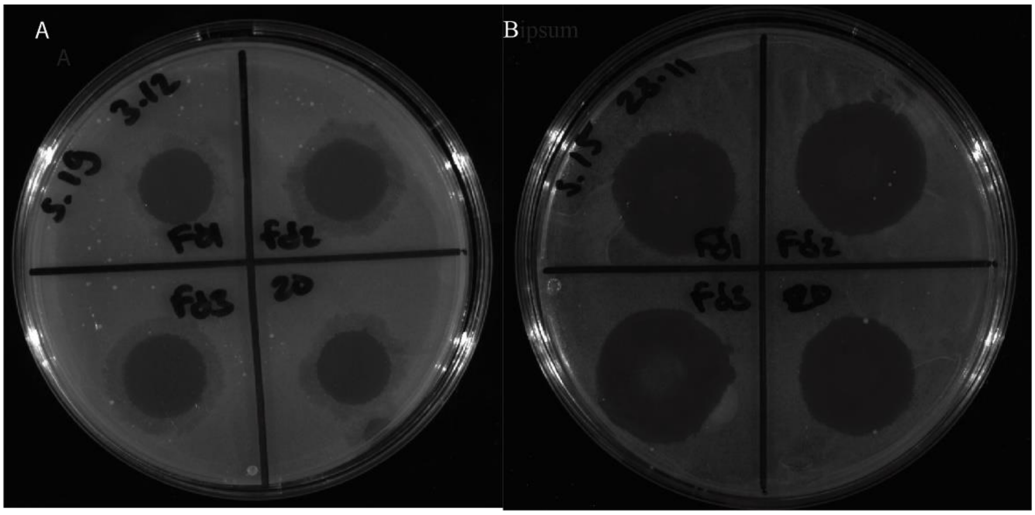


**Figure S2. -** Example of a phage susceptibility assay on strain 19 **(A).** Strain 15 **(B)**.

**Supplementary table legends**

**Table S1.**

The genome sequence details of the phages used to constract the phylogenetic tree presented in Fig. 1.

**Table S2.**

Clinical isolate number, phage, and antibiotic susceptibility. S – Susceptible, I – Intermediate, R – Resistant

**Table S3.**

Daily scoring and photography of 17 control and 17 phage-treated mice for 8 days. L- left lesion, R- right lesion. Diameter score was cacluted based on diameter measured using an electronic caliper. Elevation and eschar scores were assessed by two different blinded scorers. Total score was summed. For more details, refer to the Assessment of Phage Clinical Efficacy section in Materials and Methods and Fig. 6.

| Accession | Description | Sequence Length | Taxonomy |
|---|---|---|---|
| DQ431235 | Propionibacterium phage PA6, complete genome | 29739 | Viruses; Duplodnaviria; Heunggongvirae; Uroviricota; Caudoviricetes; Caudovirales; Siphoviridae; Pahexavirus |
| FJ706171 | Propionibacterium phage PAD20, complete genome | 29074 | Viruses; Duplodnaviria; Heunggongvirae; Uroviricota; Caudoviricetes; Caudovirales; Siphoviridae; Pahexavirus |
| FJ706172 | Propionibacterium phage PAS50, complete genome | 29017 | Viruses; Duplodnaviria; Heunggongvirae; Uroviricota; Caudoviricetes; Caudovirales; Siphoviridae; Pahexavirus |
| JX262215 | Propionibacterium phage P9.1, complete genome | 29214 | Viruses; Duplodnaviria; Heunggongvirae; Uroviricota; Caudoviricetes; Caudovirales; Siphoviridae; Pahexavirus |
| JX262216 | Propionibacterium phage P14.4, complete genome | 29729 | Viruses; Duplodnaviria; Heunggongvirae; Uroviricota; Caudoviricetes; Caudovirales; Siphoviridae; Pahexavirus |
| JX262217 | Propionibacterium phage P101A, complete genome | 29574 | Viruses; Duplodnaviria; Heunggongvirae; Uroviricota; Caudoviricetes; Caudovirales; Siphoviridae; Pahexavirus |
| JX262218 | Propionibacterium phage P104A, complete genome | 29371 | Viruses; Duplodnaviria; Heunggongvirae; Uroviricota; Caudoviricetes; Caudovirales; Siphoviridae; Pahexavirus |
| JX262219 | Propionibacterium phage P105, complete genome | 29202 | Viruses; Duplodnaviria; Heunggongvirae; Uroviricota; Caudoviricetes; Caudovirales; Siphoviridae; Pahexavirus |
| JX262220 | Propionibacterium phage P100D, complete genome | 29506 | Viruses; Duplodnaviria; Heunggongvirae; Uroviricota; Caudoviricetes; Caudovirales; Siphoviridae; Pahexavirus |
| JX262221 | Propionibacterium phage P100_A, complete genome | 29505 | Viruses; Duplodnaviria; Heunggongvirae; Uroviricota; Caudoviricetes; Caudovirales; Siphoviridae; Pahexavirus |
| JX262222 | Propionibacterium phage P100_1, complete genome | 29612 | Viruses; Duplodnaviria; Heunggongvirae; Uroviricota; Caudoviricetes; Caudovirales; Siphoviridae; Pahexavirus |
| JX262223 | Propionibacterium phage P1.1, complete genome | 29348 | Viruses; Duplodnaviria; Heunggongvirae; Uroviricota; Caudoviricetes; Caudovirales; Siphoviridae; Pahexavirus |
| JX262224 | Propionibacterium phage ATCC29399B_T, complete genome | 29516 | Viruses; Duplodnaviria; Heunggongvirae; Uroviricota; Caudoviricetes; Caudovirales; Siphoviridae; Pahexavirus |
| JX262225 | Propionibacterium phage ATCC29399B_C, complete genome | 29516 | Viruses; Duplodnaviria; Heunggongvirae; Uroviricota; Caudoviricetes; Caudovirales; Siphoviridae; Pahexavirus |
| JX570702 | Propionibacterium phage PHL111M01, complete genome | 29140 | Viruses; Duplodnaviria; Heunggongvirae; Uroviricota; Caudoviricetes; Caudovirales; Siphoviridae; Pahexavirus |
| JX570703 | Propionibacterium phage PHL073M02, complete genome | 29503 | Viruses; Duplodnaviria; Heunggongvirae; Uroviricota; Caudoviricetes; Caudovirales; Siphoviridae; Pahexavirus; unclassified Pahexavirus |
| JX570704 | Propionibacterium phage PHL010M04, complete genome | 29511 | Viruses; Duplodnaviria; Heunggongvirae; Uroviricota; Caudoviricetes; Caudovirales; Siphoviridae; Pahexavirus; unclassified Pahexavirus |
| JX570705 | Propionibacterium phage PHL060L00, complete genome | 29514 | Viruses; Duplodnaviria; Heunggongvirae; Uroviricota; Caudoviricetes; Caudovirales; Siphoviridae; Pahexavirus |
| JX570706 | Propionibacterium phage PHL037M02, complete genome | 29443 | Viruses; Duplodnaviria; Heunggongvirae; Uroviricota; Caudoviricetes; Caudovirales; Siphoviridae; Pahexavirus |
| JX570707 | Propionibacterium phage PHL085M01, complete genome | 29451 | Viruses; Duplodnaviria; Heunggongvirae; Uroviricota; Caudoviricetes; Caudovirales; Siphoviridae; Pahexavirus; unclassified Pahexavirus |
| JX570708 | Propionibacterium phage PHL115M02, complete genome | 29453 | Viruses; Duplodnaviria; Heunggongvirae; Uroviricota; Caudoviricetes; Caudovirales; Siphoviridae; Pahexavirus; unclassified Pahexavirus |
| JX570709 | Propionibacterium phage PHL067M10, complete genome | 29377 | Viruses; Duplodnaviria; Heunggongvirae; Uroviricota; Caudoviricetes; Caudovirales; Siphoviridae; Pahexavirus; unclassified Pahexavirus |
| JX570710 | Propionibacterium phage PHL071N05, complete genome | 29467 | Viruses; Duplodnaviria; Heunggongvirae; Uroviricota; Caudoviricetes; Caudovirales; Siphoviridae; Pahexavirus |
| JX570711 | Propionibacterium phage PHL066M04, complete genome | 29512 | Viruses; Duplodnaviria; Heunggongvirae; Uroviricota; Caudoviricetes; Caudovirales; Siphoviridae; Pahexavirus; unclassified Pahexavirus |
| JX570712 | Propionibacterium phage PHL114L00, complete genome | 29464 | Viruses; Duplodnaviria; Heunggongvirae; Uroviricota; Caudoviricetes; Caudovirales; Siphoviridae; Pahexavirus |
| JX570713 | Propionibacterium phage PHL113M01, complete genome | 29200 | Viruses; Duplodnaviria; Heunggongvirae; Uroviricota; Caudoviricetes; Caudovirales; Siphoviridae; Pahexavirus |
| JX570714 | Propionibacterium phage PHL112N00, complete genome | 29266 | Viruses; Duplodnaviria; Heunggongvirae; Uroviricota; Caudoviricetes; Caudovirales; Siphoviridae; Pahexavirus |
| KJ578758 | Propionibacterium phage PHL009M11, complete genome | 29503 | Viruses; Duplodnaviria; Heunggongvirae; Uroviricota; Caudoviricetes; Caudovirales; Siphoviridae; Pahexavirus |
| KJ578759 | Propionibacterium phage PHL025M00, complete genome | 29496 | Viruses; Duplodnaviria; Heunggongvirae; Uroviricota; Caudoviricetes; Caudovirales; Siphoviridae; Pahexavirus |
| KJ578760 | Propionibacterium phage PHL030N00, complete genome | 29423 | Viruses; Duplodnaviria; Heunggongvirae; Uroviricota; Caudoviricetes; Caudovirales; Siphoviridae; Pahexavirus; unclassified Pahexavirus |
| KJ578761 | Propionibacterium phage PHL041M10, complete genome | 29412 | Viruses; Duplodnaviria; Heunggongvirae; Uroviricota; Caudoviricetes; Caudovirales; Siphoviridae; Pahexavirus |
| KJ578762 | Propionibacterium phage PHL055N00, complete genome | 29264 | Viruses; Duplodnaviria; Heunggongvirae; Uroviricota; Caudoviricetes; Caudovirales; Siphoviridae; Pahexavirus; unclassified Pahexavirus |
| KJ578763 | Propionibacterium phage PHL064M01, complete genome | 29424 | Viruses; Duplodnaviria; Heunggongvirae; Uroviricota; Caudoviricetes; Caudovirales; Siphoviridae; Pahexavirus; unclassified Pahexavirus |
| KJ578764 | Propionibacterium phage PHL064M02, complete genome | 29407 | Viruses; Duplodnaviria; Heunggongvirae; Uroviricota; Caudoviricetes; Caudovirales; Siphoviridae; Pahexavirus; unclassified Pahexavirus |
| KJ578765 | Propionibacterium phage PHL067M01, complete genome | 29377 | Viruses; Duplodnaviria; Heunggongvirae; Uroviricota; Caudoviricetes; Caudovirales; Siphoviridae; Pahexavirus |
| KJ578766 | Propionibacterium phage PHL067M09, complete genome | 29386 | Viruses; Duplodnaviria; Heunggongvirae; Uroviricota; Caudoviricetes; Caudovirales; Siphoviridae; Pahexavirus; unclassified Pahexavirus |
| KJ578767 | Propionibacterium phage PHL070N00, complete genome | 29421 | Viruses; Duplodnaviria; Heunggongvirae; Uroviricota; Caudoviricetes; Caudovirales; Siphoviridae; Pahexavirus |
| KJ578768 | Propionibacterium phage PHL082M00, complete genome | 29491 | Viruses; Duplodnaviria; Heunggongvirae; Uroviricota; Caudoviricetes; Caudovirales; Siphoviridae; Pahexavirus; unclassified Pahexavirus |
| KJ578769 | Propionibacterium phage PHL082M02, complete genome | 29491 | Viruses; Duplodnaviria; Heunggongvirae; Uroviricota; Caudoviricetes; Caudovirales; Siphoviridae; Pahexavirus; unclassified Pahexavirus |
| KJ578770 | Propionibacterium phage PHL082M03, complete genome | 29491 | Viruses; Duplodnaviria; Heunggongvirae; Uroviricota; Caudoviricetes; Caudovirales; Siphoviridae; Pahexavirus |
| KJ578771 | Propionibacterium phage PHL082M04, complete genome | 29491 | Viruses; Duplodnaviria; Heunggongvirae; Uroviricota; Caudoviricetes; Caudovirales; Siphoviridae; Pahexavirus; unclassified Pahexavirus |
| KJ578772 | Propionibacterium phage PHL085N00, complete genome | 29454 | Viruses; Duplodnaviria; Heunggongvirae; Uroviricota; Caudoviricetes; Caudovirales; Siphoviridae; Pahexavirus; unclassified Pahexavirus |
| KJ578773 | Propionibacterium phage PHL092M00, complete genome | 29261 | Viruses; Duplodnaviria; Heunggongvirae; Uroviricota; Caudoviricetes; Caudovirales; Siphoviridae; Pahexavirus |
| KJ578774 | Propionibacterium phage PHL095N00, complete genome | 29751 | Viruses; Duplodnaviria; Heunggongvirae; Uroviricota; Caudoviricetes; Caudovirales; Siphoviridae; Pahexavirus |
| KJ578775 | Propionibacterium phage PHL114N00, complete genome | 29464 | Viruses; Duplodnaviria; Heunggongvirae; Uroviricota; Caudoviricetes; Caudovirales; Siphoviridae; Pahexavirus |
| KJ578776 | Propionibacterium phage PHL116M00, complete genome | 29394 | Viruses; Duplodnaviria; Heunggongvirae; Uroviricota; Caudoviricetes; Caudovirales; Siphoviridae; Pahexavirus |
| KJ578777 | Propionibacterium phage PHL116M10, complete genome | 29396 | Viruses; Duplodnaviria; Heunggongvirae; Uroviricota; Caudoviricetes; Caudovirales; Siphoviridae; Pahexavirus; unclassified Pahexavirus |
| KJ578778 | Propionibacterium phage PHL117M00, complete genome | 29255 | Viruses; Duplodnaviria; Heunggongvirae; Uroviricota; Caudoviricetes; Caudovirales; Siphoviridae; Pahexavirus |
| KJ578779 | Propionibacterium phage PHL117M01, complete genome | 29422 | Viruses; Duplodnaviria; Heunggongvirae; Uroviricota; Caudoviricetes; Caudovirales; Siphoviridae; Pahexavirus |
| KJ578780 | Propionibacterium phage PHL132N00, complete genome | 29003 | Viruses; Duplodnaviria; Heunggongvirae; Uroviricota; Caudoviricetes; Caudovirales; Siphoviridae; Pahexavirus |
| KJ578781 | Propionibacterium phage PHL141N00, complete genome | 29494 | Viruses; Duplodnaviria; Heunggongvirae; Uroviricota; Caudoviricetes; Caudovirales; Siphoviridae; Pahexavirus |
| KJ578782 | Propionibacterium phage PHL150M00, complete genome | 29438 | Viruses; Duplodnaviria; Heunggongvirae; Uroviricota; Caudoviricetes; Caudovirales; Siphoviridae; Pahexavirus; unclassified Pahexavirus |
| KJ578783 | Propionibacterium phage PHL151M00, complete genome | 29511 | Viruses; Duplodnaviria; Heunggongvirae; Uroviricota; Caudoviricetes; Caudovirales; Siphoviridae; Pahexavirus |
| KJ578784 | Propionibacterium phage PHL151N00, complete genome | 29511 | Viruses; Duplodnaviria; Heunggongvirae; Uroviricota; Caudoviricetes; Caudovirales; Siphoviridae; Pahexavirus |
| KJ578785 | Propionibacterium phage PHL152M00, complete genome | 29247 | Viruses; Duplodnaviria; Heunggongvirae; Uroviricota; Caudoviricetes; Caudovirales; Siphoviridae; Pahexavirus |
| KJ578786 | Propionibacterium phage PHL163M00, complete genome | 29264 | Viruses; Duplodnaviria; Heunggongvirae; Uroviricota; Caudoviricetes; Caudovirales; Siphoviridae; Pahexavirus |
| KJ578787 | Propionibacterium phage PHL171M01, complete genome | 29327 | Viruses; Duplodnaviria; Heunggongvirae; Uroviricota; Caudoviricetes; Caudovirales; Siphoviridae; Pahexavirus |
| KJ578788 | Propionibacterium phage PHL179M00, complete genome | 29428 | Viruses; Duplodnaviria; Heunggongvirae; Uroviricota; Caudoviricetes; Caudovirales; Siphoviridae; Pahexavirus |
| KJ578789 | Propionibacterium phage PHL194M00, complete genome | 29264 | Viruses; Duplodnaviria; Heunggongvirae; Uroviricota; Caudoviricetes; Caudovirales; Siphoviridae; Pahexavirus |
| KJ578790 | Propionibacterium phage PHL199M00, complete genome | 29806 | Viruses; Duplodnaviria; Heunggongvirae; Uroviricota; Caudoviricetes; Caudovirales; Siphoviridae; Pahexavirus |
| KJ578791 | Propionibacterium phage PHL301M00, complete genome | 29323 | Viruses; Duplodnaviria; Heunggongvirae; Uroviricota; Caudoviricetes; Caudovirales; Siphoviridae; Pahexavirus |
| KJ578792 | Propionibacterium phage PHL308M00, complete genome | 29442 | Viruses; Duplodnaviria; Heunggongvirae; Uroviricota; Caudoviricetes; Caudovirales; Siphoviridae; Pahexavirus |
| KJ722067 | Propionibacterium phage Pacnes 2012-15, complete genome | 29741 | Viruses; Duplodnaviria; Heunggongvirae; Uroviricota; Caudoviricetes; Caudovirales; Siphoviridae; Pahexavirus |
| KR337643 | Propionibacterium phage MrAK, complete genome | 29726 | Viruses; Duplodnaviria; Heunggongvirae; Uroviricota; Caudoviricetes; Caudovirales; Siphoviridae; Pahexavirus |
| KR337644 | Propionibacterium phage Procrass1, complete genome | 29347 | Viruses; Duplodnaviria; Heunggongvirae; Uroviricota; Caudoviricetes; Caudovirales; Siphoviridae; Pahexavirus |
| KR337645 | Propionibacterium phage Kubed, complete genome | 29461 | Viruses; Duplodnaviria; Heunggongvirae; Uroviricota; Caudoviricetes; Caudovirales; Siphoviridae; Pahexavirus |
| KR337646 | Propionibacterium phage Wizzo, complete genome | 29463 | Viruses; Duplodnaviria; Heunggongvirae; Uroviricota; Caudoviricetes; Caudovirales; Siphoviridae; Pahexavirus |
| KR337647 | Propionibacterium phage Solid, complete genome | 29440 | Viruses; Duplodnaviria; Heunggongvirae; Uroviricota; Caudoviricetes; Caudovirales; Siphoviridae; Pahexavirus |
| KR337648 | Propionibacterium phage SKKY, complete genome | 29594 | Viruses; Duplodnaviria; Heunggongvirae; Uroviricota; Caudoviricetes; Caudovirales; Siphoviridae; Pahexavirus |
| KR337649 | Propionibacterium phage Keiki, complete genome | 29339 | Viruses; Duplodnaviria; Heunggongvirae; Uroviricota; Caudoviricetes; Caudovirales; Siphoviridae; Pahexavirus |
| KR337650 | Propionibacterium phage Lauchelly, complete genome | 29517 | Viruses; Duplodnaviria; Heunggongvirae; Uroviricota; Caudoviricetes; Caudovirales; Siphoviridae; Pahexavirus |
| KR337651 | Propionibacterium phage Attacne, complete genome | 28876 | Viruses; Duplodnaviria; Heunggongvirae; Uroviricota; Caudoviricetes; Caudovirales; Siphoviridae; Pahexavirus |
| KR337652 | Propionibacterium phage Stormborn, complete genome | 29330 | Viruses; Duplodnaviria; Heunggongvirae; Uroviricota; Caudoviricetes; Caudovirales; Siphoviridae; Pahexavirus |
| KR337653 | Propionibacterium phage Pirate, complete genome | 29324 | Viruses; Duplodnaviria; Heunggongvirae; Uroviricota; Caudoviricetes; Caudovirales; Siphoviridae; Pahexavirus |
| KR337654 | Propionibacterium phage Ouroboros, complete genome | 29506 | Viruses; Duplodnaviria; Heunggongvirae; Uroviricota; Caudoviricetes; Caudovirales; Siphoviridae; Pahexavirus |
| KR347352 | Propionibacterium phage BruceLethal, complete genome | 29249 | Viruses; Duplodnaviria; Heunggongvirae; Uroviricota; Caudoviricetes; Caudovirales; Siphoviridae; Pahexavirus; unclassified Pahexavirus |
| KR347353 | Propionibacterium phage Enoki, complete genome | 29347 | Viruses; Duplodnaviria; Heunggongvirae; Uroviricota; Caudoviricetes; Caudovirales; Siphoviridae; Pahexavirus; unclassified Pahexavirus |
| KR347354 | Propionibacterium phage Moyashi, complete genome | 29254 | Viruses; Duplodnaviria; Heunggongvirae; Uroviricota; Caudoviricetes; Caudovirales; Siphoviridae; Pahexavirus; unclassified Pahexavirus |
| KR347355 | Propionibacterium phage QueenBey, complete genome | 29338 | Viruses; Duplodnaviria; Heunggongvirae; Uroviricota; Caudoviricetes; Caudovirales; Siphoviridae; Pahexavirus; unclassified Pahexavirus |
| KR902978 | Propionibacterium phage PAC1, complete genome | 29605 | Viruses; Duplodnaviria; Heunggongvirae; Uroviricota; Caudoviricetes; Caudovirales; Siphoviridae; Pahexavirus; unclassified Pahexavirus |
| KR902979 | Propionibacterium phage PAC2, complete genome | 29602 | Viruses; Duplodnaviria; Heunggongvirae; Uroviricota; Caudoviricetes; Caudovirales; Siphoviridae; Pahexavirus; unclassified Pahexavirus |
| KR902980 | Propionibacterium phage PAC3, complete genome | 29526 | Viruses; Duplodnaviria; Heunggongvirae; Uroviricota; Caudoviricetes; Caudovirales; Siphoviridae; Pahexavirus; unclassified Pahexavirus |
| KR902981 | Propionibacterium phage PAC4, complete genome | 29581 | Viruses; Duplodnaviria; Heunggongvirae; Uroviricota; Caudoviricetes; Caudovirales; Siphoviridae; Pahexavirus; unclassified Pahexavirus |
| KR902982 | Propionibacterium phage PAC5, complete genome | 29428 | Viruses; Duplodnaviria; Heunggongvirae; Uroviricota; Caudoviricetes; Caudovirales; Siphoviridae; Pahexavirus; unclassified Pahexavirus |
| KR902983 | Propionibacterium phage PAC6, complete genome | 29609 | Viruses; Duplodnaviria; Heunggongvirae; Uroviricota; Caudoviricetes; Caudovirales; Siphoviridae; Pahexavirus; unclassified Pahexavirus |
| KR902984 | Propionibacterium phage PAC7, complete genome | 29551 | Viruses; Duplodnaviria; Heunggongvirae; Uroviricota; Caudoviricetes; Caudovirales; Siphoviridae; Pahexavirus; unclassified Pahexavirus |
| KR902985 | Propionibacterium phage PAC8, complete genome | 29488 | Viruses; Duplodnaviria; Heunggongvirae; Uroviricota; Caudoviricetes; Caudovirales; Siphoviridae; Pahexavirus; unclassified Pahexavirus |
| KR902986 | Propionibacterium phage PAC9, complete genome | 29786 | Viruses; Duplodnaviria; Heunggongvirae; Uroviricota; Caudoviricetes; Caudovirales; Siphoviridae; Pahexavirus; unclassified Pahexavirus |
| KR902987 | Propionibacterium phage PAC10, complete genome | 29704 | Viruses; Duplodnaviria; Heunggongvirae; Uroviricota; Caudoviricetes; Caudovirales; Siphoviridae; Pahexavirus; unclassified Pahexavirus |
| KT934381 | Propionibacterium phage PA1-14, complete genome | 29407 | Viruses; Duplodnaviria; Heunggongvirae; Uroviricota; Caudoviricetes; Caudovirales; Siphoviridae; Pahexavirus; unclassified Pahexavirus |
| KY926792 | Propionibacterium phage PacnesP1, complete genome | 29533 | Viruses; Duplodnaviria; Heunggongvirae; Uroviricota; Caudoviricetes; Caudovirales; Siphoviridae; Pahexavirus; unclassified Pahexavirus |
| KY926793 | Propionibacterium phage PacnesP2, complete genome | 30016 | Viruses; Duplodnaviria; Heunggongvirae; Uroviricota; Caudoviricetes; Caudovirales; Siphoviridae; Pahexavirus; unclassified Pahexavirus |
| MF919491 | Propionibacterium phage Aquarius, complete genome | 30112 | Viruses; Duplodnaviria; Heunggongvirae; Uroviricota; Caudoviricetes; Caudovirales; Siphoviridae; Pahexavirus; unclassified Pahexavirus |
| MF919505 | Propionibacterium phage DrParker, complete genome | 29742 | Viruses; Duplodnaviria; Heunggongvirae; Uroviricota; Caudoviricetes; Caudovirales; Siphoviridae; Pahexavirus; unclassified Pahexavirus |
| MF919515 | Propionibacterium phage Leviosa, complete genome | 29451 | Viruses; Duplodnaviria; Heunggongvirae; Uroviricota; Caudoviricetes; Caudovirales; Siphoviridae; Pahexavirus; unclassified Pahexavirus |
| MF919516 | Propionibacterium phage LilBandit, complete genome | 29041 | Viruses; Duplodnaviria; Heunggongvirae; Uroviricota; Caudoviricetes; Caudovirales; Siphoviridae; Pahexavirus; unclassified Pahexavirus |
| MF919522 | Propionibacterium phage MEAK, complete genome | 29223 | Viruses; Duplodnaviria; Heunggongvirae; Uroviricota; Caudoviricetes; Caudovirales; Siphoviridae; Pahexavirus; unclassified Pahexavirus |
| MF919533 | Propionibacterium phage Supernova, complete genome | 29217 | Viruses; Duplodnaviria; Heunggongvirae; Uroviricota; Caudoviricetes; Caudovirales; Siphoviridae; Pahexavirus; unclassified Pahexavirus |
| MG820632 | Propionibacterium phage pa310, complete genome | 29508 | Viruses; Duplodnaviria; Heunggongvirae; Uroviricota; Caudoviricetes; Caudovirales; Siphoviridae; Pahexavirus; unclassified Pahexavirus |
| MG820633 | Propionibacterium phage pa59, complete genome | 29507 | Viruses; Duplodnaviria; Heunggongvirae; Uroviricota; Caudoviricetes; Caudovirales; Siphoviridae; Pahexavirus; unclassified Pahexavirus |
| MG820634 | Propionibacterium phage pa27, complete genome | 29593 | Viruses; Duplodnaviria; Heunggongvirae; Uroviricota; Caudoviricetes; Caudovirales; Siphoviridae; Pahexavirus; unclassified Pahexavirus |
| MG820635 | Propionibacterium phage pa29399-1-D_1, complete genome | 29370 | Viruses; Duplodnaviria; Heunggongvirae; Uroviricota; Caudoviricetes; Caudovirales; Siphoviridae; Pahexavirus; unclassified Pahexavirus |
| MG820636 | Propionibacterium phage pa29399-1-D_2, complete genome | 28892 | Viruses; Duplodnaviria; Heunggongvirae; Uroviricota; Caudoviricetes; Caudovirales; Siphoviridae; Pahexavirus; unclassified Pahexavirus |
| MG820637 | Propionibacterium phage pa63, complete genome | 29446 | Viruses; Duplodnaviria; Heunggongvirae; Uroviricota; Caudoviricetes; Caudovirales; Siphoviridae; Pahexavirus; unclassified Pahexavirus |
| MG820638 | Propionibacterium phage pa6919-4, complete genome | 29784 | Viruses; Duplodnaviria; Heunggongvirae; Uroviricota; Caudoviricetes; Caudovirales; Siphoviridae; Pahexavirus; unclassified Pahexavirus |
| MG820639 | Propionibacterium phage pa9-6919-4, complete genome | 29784 | Viruses; Duplodnaviria; Heunggongvirae; Uroviricota; Caudoviricetes; Caudovirales; Siphoviridae; Pahexavirus; unclassified Pahexavirus |
| MG820640 | Propionibacterium phage pa15, complete genome | 29309 | Viruses; Duplodnaviria; Heunggongvirae; Uroviricota; Caudoviricetes; Caudovirales; Siphoviridae; Pahexavirus; unclassified Pahexavirus |
| MG820641 | Propionibacterium phage pa615, complete genome | 29307 | Viruses; Duplodnaviria; Heunggongvirae; Uroviricota; Caudoviricetes; Caudovirales; Siphoviridae; Pahexavirus; unclassified Pahexavirus |
| MG820642 | Propionibacterium phage pa28, complete genome | 29733 | Viruses; Duplodnaviria; Heunggongvirae; Uroviricota; Caudoviricetes; Caudovirales; Siphoviridae; Pahexavirus; unclassified Pahexavirus |
| MG820643 | Propionibacterium phage pa35, complete genome | 29491 | Viruses; Duplodnaviria; Heunggongvirae; Uroviricota; Caudoviricetes; Caudovirales; Siphoviridae; Pahexavirus; unclassified Pahexavirus |
| MG820644 | Propionibacterium phage pa33, complete genome | 29135 | Viruses; Duplodnaviria; Heunggongvirae; Uroviricota; Caudoviricetes; Caudovirales; Siphoviridae; Pahexavirus; unclassified Pahexavirus |
| MG820645 | Propionibacterium phage pa3-SS3, complete genome | 29137 | Viruses; Duplodnaviria; Heunggongvirae; Uroviricota; Caudoviricetes; Caudovirales; Siphoviridae; Pahexavirus; unclassified Pahexavirus |
| MN329678 | Propionibacterium phage Cota, complete genome | 29549 | Viruses; Duplodnaviria; Heunggongvirae; Uroviricota; Caudoviricetes; Caudovirales; Siphoviridae; Pahexavirus; unclassified Pahexavirus |
| MN813675 | Cutibacterium phage P104B, complete genome | 29330 | Viruses; Duplodnaviria; Heunggongvirae; Uroviricota; Caudoviricetes; Caudovirales; Siphoviridae; Pahexavirus; unclassified Pahexavirus |
| MN813677 | Cutibacterium phage P107C, complete genome | 29516 | Viruses; Duplodnaviria; Heunggongvirae; Uroviricota; Caudoviricetes; Caudovirales; Siphoviridae; Pahexavirus; unclassified Pahexavirus |
| MN813679 | Cutibacterium phage P107A, complete genome | 29465 | Viruses; Duplodnaviria; Heunggongvirae; Uroviricota; Caudoviricetes; Caudovirales; Siphoviridae; Pahexavirus; unclassified Pahexavirus |
| MN813683 | Cutibacterium phage P108C, complete genome | 29494 | Viruses; Duplodnaviria; Heunggongvirae; Uroviricota; Caudoviricetes; Caudovirales; Siphoviridae; Pahexavirus; unclassified Pahexavirus |
| MN813688 | Cutibacterium phage P106I, complete genome | 29470 | Viruses; Duplodnaviria; Heunggongvirae; Uroviricota; Caudoviricetes; Caudovirales; Siphoviridae; Pahexavirus; unclassified Pahexavirus |
| MN813689 | Cutibacterium phage P106L, complete genome | 29641 | Viruses; Duplodnaviria; Heunggongvirae; Uroviricota; Caudoviricetes; Caudovirales; Siphoviridae; Pahexavirus; unclassified Pahexavirus |
| MN813690 | Cutibacterium phage P106M, complete genome | 29641 | Viruses; Duplodnaviria; Heunggongvirae; Uroviricota; Caudoviricetes; Caudovirales; Siphoviridae; Pahexavirus; unclassified Pahexavirus |
| MN813691 | Cutibacterium phage P106A, complete genome | 29554 | Viruses; Duplodnaviria; Heunggongvirae; Uroviricota; Caudoviricetes; Caudovirales; Siphoviridae; Pahexavirus; unclassified Pahexavirus |
| MN813692 | Cutibacterium phage P106C, complete genome | 29641 | Viruses; Duplodnaviria; Heunggongvirae; Uroviricota; Caudoviricetes; Caudovirales; Siphoviridae; Pahexavirus; unclassified Pahexavirus |
| MT647607 | Propionibacterium phage phiPA50S, complete genome | 29590 | Viruses; Duplodnaviria; Heunggongvirae; Uroviricota; Caudoviricetes; Caudovirales; Siphoviridae; Pahexavirus; unclassified Pahexavirus |
| MW161461 | Cutibacterium phage FD1, complete genome | 29774 | Viruses; Duplodnaviria; Heunggongvirae; Uroviricota; Caudoviricetes; Caudovirales; Siphoviridae; Pahexavirus; unclassified Pahexavirus |
| MW161462 | Cutibacterium phage FD2, complete genome | 29768 | Viruses; Duplodnaviria; Heunggongvirae; Uroviricota; Caudoviricetes; Caudovirales; Siphoviridae; Pahexavirus; unclassified Pahexavirus |
| MW161463 | Cutibacterium phage FD3, complete genome | 29638 | Viruses; Duplodnaviria; Heunggongvirae; Uroviricota; Caudoviricetes; Caudovirales; Siphoviridae; Pahexavirus; unclassified Pahexavirus |
| MW161464 | Cutibacterium phage PAVL20, complete genome | 29800 | Viruses; Duplodnaviria; Heunggongvirae; Uroviricota; Caudoviricetes; Caudovirales; Siphoviridae; Pahexavirus; unclassified Pahexavirus |
| MW161465 | Cutibacterium phage PAVL21, complete genome | 30034 | Viruses; Duplodnaviria; Heunggongvirae; Uroviricota; Caudoviricetes; Caudovirales; Siphoviridae; Pahexavirus; unclassified Pahexavirus |

| Accession | Description | Sequence Length | Taxonomy |
|---|---|---|---|
| MW161466 | Cutibacterium phage PAVL33, complete genome | 29627 | Viruses; Duplodnaviria; Heunggongvirae; Uroviricota; Caudoviricetes; Caudovirales; Siphoviridae; Pahexavirus; unclassified Pahexavirus |
| MW161467 | Cutibacterium phage PAVL34, complete genome | 29535 | Viruses; Duplodnaviria; Heunggongvirae; Uroviricota; Caudoviricetes; Caudovirales; Siphoviridae; Pahexavirus; unclassified Pahexavirus |
| MW161468 | Cutibacterium phage PAVL45, complete genome | 29772 | Viruses; Duplodnaviria; Heunggongvirae; Uroviricota; Caudoviricetes; Caudovirales; Siphoviridae; Pahexavirus; unclassified Pahexavirus |
| NC_009541 | Propionibacterium phage PA6, complete genome | 29739 | Viruses; Duplodnaviria; Heunggongvirae; Uroviricota; Caudoviricetes; Caudovirales; Siphoviridae; Pahexavirus |
| NC_015453 | Propionibacterium phage PAS50 endogenous virus, complete genome | 29017 | Viruses; Duplodnaviria; Heunggongvirae; Uroviricota; Caudoviricetes; Caudovirales; Siphoviridae; Pahexavirus |
| NC_015454 | Propionibacterium phage PAD20 endogenous virus, complete genome | 29074 | Viruses; Duplodnaviria; Heunggongvirae; Uroviricota; Caudoviricetes; Caudovirales; Siphoviridae; Pahexavirus |
| NC_018834 | Propionibacterium phage P9.1, complete genome | 29214 | Viruses; Duplodnaviria; Heunggongvirae; Uroviricota; Caudoviricetes; Caudovirales; Siphoviridae; Pahexavirus |
| NC_018838 | Propionibacterium phage P100_A, complete genome | 29505 | Viruses; Duplodnaviria; Heunggongvirae; Uroviricota; Caudoviricetes; Caudovirales; Siphoviridae; Pahexavirus |
| NC_018839 | Propionibacterium phage P14.4, complete genome | 29729 | Viruses; Duplodnaviria; Heunggongvirae; Uroviricota; Caudoviricetes; Caudovirales; Siphoviridae; Pahexavirus |
| NC_018840 | Propionibacterium phage P100_1, complete genome | 29612 | Viruses; Duplodnaviria; Heunggongvirae; Uroviricota; Caudoviricetes; Caudovirales; Siphoviridae; Pahexavirus |
| NC_018841 | Propionibacterium phage P101A, complete genome | 29574 | Viruses; Duplodnaviria; Heunggongvirae; Uroviricota; Caudoviricetes; Caudovirales; Siphoviridae; Pahexavirus |
| NC_018842 | Propionibacterium phage P1.1, complete genome | 29348 | Viruses; Duplodnaviria; Heunggongvirae; Uroviricota; Caudoviricetes; Caudovirales; Siphoviridae; Pahexavirus |
| NC_018845 | Propionibacterium phage P104A, complete genome | 29371 | Viruses; Duplodnaviria; Heunggongvirae; Uroviricota; Caudoviricetes; Caudovirales; Siphoviridae; Pahexavirus |
| NC_018847 | Propionibacterium phage ATCC29399B_T, complete genome | 29516 | Viruses; Duplodnaviria; Heunggongvirae; Uroviricota; Caudoviricetes; Caudovirales; Siphoviridae; Pahexavirus |
| NC_018849 | Propionibacterium phage P105, complete genome | 29202 | Viruses; Duplodnaviria; Heunggongvirae; Uroviricota; Caudoviricetes; Caudovirales; Siphoviridae; Pahexavirus |
| NC_018851 | Propionibacterium phage ATCC29399B_C, complete genome | 29516 | Viruses; Duplodnaviria; Heunggongvirae; Uroviricota; Caudoviricetes; Caudovirales; Siphoviridae; Pahexavirus |
| NC_018852 | Propionibacterium phage P100D, complete genome | 29506 | Viruses; Duplodnaviria; Heunggongvirae; Uroviricota; Caudoviricetes; Caudovirales; Siphoviridae; Pahexavirus |
| NC_022334 | Propionibacterium phage PHL112N00, complete genome | 29266 | Viruses; Duplodnaviria; Heunggongvirae; Uroviricota; Caudoviricetes; Caudovirales; Siphoviridae; Pahexavirus |
| NC_022335 | Propionibacterium phage PHL067M10, complete genome | 29377 | Viruses; Duplodnaviria; Heunggongvirae; Uroviricota; Caudoviricetes; Caudovirales; Siphoviridae; Pahexavirus; unclassified Pahexavirus |
| NC_022336 | Propionibacterium phage PHL010M04, complete genome | 29511 | Viruses; Duplodnaviria; Heunggongvirae; Uroviricota; Caudoviricetes; Caudovirales; Siphoviridae; Pahexavirus; unclassified Pahexavirus |
| NC_022337 | Propionibacterium phage PHL071N05, complete genome | 29467 | Viruses; Duplodnaviria; Heunggongvirae; Uroviricota; Caudoviricetes; Caudovirales; Siphoviridae; Pahexavirus |
| NC_022338 | Propionibacterium phage PHL060L00, complete genome | 29514 | Viruses; Duplodnaviria; Heunggongvirae; Uroviricota; Caudoviricetes; Caudovirales; Siphoviridae; Pahexavirus |
| NC_022339 | Propionibacterium phage PHL037M02, complete genome | 29443 | Viruses; Duplodnaviria; Heunggongvirae; Uroviricota; Caudoviricetes; Caudovirales; Siphoviridae; Pahexavirus |
| NC_022340 | Propionibacterium phage PHL114L00, complete genome | 29464 | Viruses; Duplodnaviria; Heunggongvirae; Uroviricota; Caudoviricetes; Caudovirales; Siphoviridae; Pahexavirus |
| NC_022341 | Propionibacterium phage PHL113M01, complete genome | 29200 | Viruses; Duplodnaviria; Heunggongvirae; Uroviricota; Caudoviricetes; Caudovirales; Siphoviridae; Pahexavirus |
| NC_022342 | Propionibacterium phage PHL111M01, complete genome | 29140 | Viruses; Duplodnaviria; Heunggongvirae; Uroviricota; Caudoviricetes; Caudovirales; Siphoviridae; Pahexavirus |
| NC_027294 | Propionibacterium phage PHL150M00, complete genome | 29438 | Viruses; Duplodnaviria; Heunggongvirae; Uroviricota; Caudoviricetes; Caudovirales; Siphoviridae; Pahexavirus; unclassified Pahexavirus |
| NC_027295 | Propionibacterium phage PHL199M00, complete genome | 29806 | Viruses; Duplodnaviria; Heunggongvirae; Uroviricota; Caudoviricetes; Caudovirales; Siphoviridae; Pahexavirus |
| NC_027333 | Propionibacterium phage PHL070N00, complete genome | 29421 | Viruses; Duplodnaviria; Heunggongvirae; Uroviricota; Caudoviricetes; Caudovirales; Siphoviridae; Pahexavirus |
| NC_027336 | Propionibacterium phage PHL009M11, complete genome | 29503 | Viruses; Duplodnaviria; Heunggongvirae; Uroviricota; Caudoviricetes; Caudovirales; Siphoviridae; Pahexavirus |
| NC_027346 | Propionibacterium phage PHL171M01, complete genome | 29327 | Viruses; Duplodnaviria; Heunggongvirae; Uroviricota; Caudoviricetes; Caudovirales; Siphoviridae; Pahexavirus |
| NC_027347 | Propionibacterium phage PHL151M00, complete genome | 29511 | Viruses; Duplodnaviria; Heunggongvirae; Uroviricota; Caudoviricetes; Caudovirales; Siphoviridae; Pahexavirus |
| NC_027354 | Propionibacterium phage PHL301M00, complete genome | 29323 | Viruses; Duplodnaviria; Heunggongvirae; Uroviricota; Caudoviricetes; Caudovirales; Siphoviridae; Pahexavirus |
| NC_027357 | Propionibacterium phage PHL025M00, complete genome | 29496 | Viruses; Duplodnaviria; Heunggongvirae; Uroviricota; Caudoviricetes; Caudovirales; Siphoviridae; Pahexavirus |
| NC_027359 | Propionibacterium phage PHL082M00, complete genome | 29491 | Viruses; Duplodnaviria; Heunggongvirae; Uroviricota; Caudoviricetes; Caudovirales; Siphoviridae; Pahexavirus; unclassified Pahexavirus |
| NC_027361 | Propionibacterium phage PHL085N00, complete genome | 29454 | Viruses; Duplodnaviria; Heunggongvirae; Uroviricota; Caudoviricetes; Caudovirales; Siphoviridae; Pahexavirus; unclassified Pahexavirus |
| NC_027362 | Propionibacterium phage PHL116M00, complete genome | 29394 | Viruses; Duplodnaviria; Heunggongvirae; Uroviricota; Caudoviricetes; Caudovirales; Siphoviridae; Pahexavirus |
| NC_027367 | Propionibacterium phage PHL132N00, complete genome | 29003 | Viruses; Duplodnaviria; Heunggongvirae; Uroviricota; Caudoviricetes; Caudovirales; Siphoviridae; Pahexavirus |
| NC_027370 | Propionibacterium phage PHL179M00, complete genome | 29428 | Viruses; Duplodnaviria; Heunggongvirae; Uroviricota; Caudoviricetes; Caudovirales; Siphoviridae; Pahexavirus |
| NC_027371 | Propionibacterium phage Pacnes 2012-15, complete genome | 29741 | Viruses; Duplodnaviria; Heunggongvirae; Uroviricota; Caudoviricetes; Caudovirales; Siphoviridae; Pahexavirus |
| NC_027373 | Propionibacterium phage PHL030N00, complete genome | 29423 | Viruses; Duplodnaviria; Heunggongvirae; Uroviricota; Caudoviricetes; Caudovirales; Siphoviridae; Pahexavirus; unclassified Pahexavirus |
| NC_027376 | Propionibacterium phage PHL308M00, complete genome | 29442 | Viruses; Duplodnaviria; Heunggongvirae; Uroviricota; Caudoviricetes; Caudovirales; Siphoviridae; Pahexavirus |
| NC_027380 | Propionibacterium phage PHL067M01, complete genome | 29377 | Viruses; Duplodnaviria; Heunggongvirae; Uroviricota; Caudoviricetes; Caudovirales; Siphoviridae; Pahexavirus |
| NC_027385 | Propionibacterium phage PHL092M00, complete genome | 29261 | Viruses; Duplodnaviria; Heunggongvirae; Uroviricota; Caudoviricetes; Caudovirales; Siphoviridae; Pahexavirus |
| NC_027386 | Propionibacterium phage PHL152M00, complete genome | 29247 | Viruses; Duplodnaviria; Heunggongvirae; Uroviricota; Caudoviricetes; Caudovirales; Siphoviridae; Pahexavirus |
| NC_027389 | Propionibacterium phage PHL141N00, complete genome | 29494 | Viruses; Duplodnaviria; Heunggongvirae; Uroviricota; Caudoviricetes; Caudovirales; Siphoviridae; Pahexavirus |
| NC_027391 | Propionibacterium phage PHL041M10, complete genome | 29412 | Viruses; Duplodnaviria; Heunggongvirae; Uroviricota; Caudoviricetes; Caudovirales; Siphoviridae; Pahexavirus |
| NC_027392 | Propionibacterium phage PHL194M00, complete genome | 29264 | Viruses; Duplodnaviria; Heunggongvirae; Uroviricota; Caudoviricetes; Caudovirales; Siphoviridae; Pahexavirus |
| NC_027400 | Propionibacterium phage PHL055N00, complete genome | 29264 | Viruses; Duplodnaviria; Heunggongvirae; Uroviricota; Caudoviricetes; Caudovirales; Siphoviridae; Pahexavirus; unclassified Pahexavirus |
| NC_027401 | Propionibacterium phage PHL095N00, complete genome | 29751 | Viruses; Duplodnaviria; Heunggongvirae; Uroviricota; Caudoviricetes; Caudovirales; Siphoviridae; Pahexavirus |
| NC_027403 | Propionibacterium phage PHL117M00, complete genome | 29255 | Viruses; Duplodnaviria; Heunggongvirae; Uroviricota; Caudoviricetes; Caudovirales; Siphoviridae; Pahexavirus |
| NC_027405 | Propionibacterium phage PHL163M00, complete genome | 29264 | Viruses; Duplodnaviria; Heunggongvirae; Uroviricota; Caudoviricetes; Caudovirales; Siphoviridae; Pahexavirus |
| NC_027620 | Propionibacterium phage MrAK, complete genome | 29726 | Viruses; Duplodnaviria; Heunggongvirae; Uroviricota; Caudoviricetes; Caudovirales; Siphoviridae; Pahexavirus |
| NC_027621 | Propionibacterium phage Wizzo, complete genome | 29463 | Viruses; Duplodnaviria; Heunggongvirae; Uroviricota; Caudoviricetes; Caudovirales; Siphoviridae; Pahexavirus |
| NC_027622 | Propionibacterium phage Stormborn, complete genome | 29330 | Viruses; Duplodnaviria; Heunggongvirae; Uroviricota; Caudoviricetes; Caudovirales; Siphoviridae; Pahexavirus |
| NC_027623 | Propionibacterium phage Pirate, complete genome | 29324 | Viruses; Duplodnaviria; Heunggongvirae; Uroviricota; Caudoviricetes; Caudovirales; Siphoviridae; Pahexavirus |
| NC_027624 | Propionibacterium phage SKKY, complete genome | 29594 | Viruses; Duplodnaviria; Heunggongvirae; Uroviricota; Caudoviricetes; Caudovirales; Siphoviridae; Pahexavirus |
| NC_027625 | Propionibacterium phage Kubed, complete genome | 29461 | Viruses; Duplodnaviria; Heunggongvirae; Uroviricota; Caudoviricetes; Caudovirales; Siphoviridae; Pahexavirus |
| NC_027626 | Propionibacterium phage Procrass1, complete genome | 29347 | Viruses; Duplodnaviria; Heunggongvirae; Uroviricota; Caudoviricetes; Caudovirales; Siphoviridae; Pahexavirus |
| NC_027627 | Propionibacterium phage Solid, complete genome | 29440 | Viruses; Duplodnaviria; Heunggongvirae; Uroviricota; Caudoviricetes; Caudovirales; Siphoviridae; Pahexavirus |
| NC_027628 | Propionibacterium phage Lauchelly, complete genome | 29517 | Viruses; Duplodnaviria; Heunggongvirae; Uroviricota; Caudoviricetes; Caudovirales; Siphoviridae; Pahexavirus |
| NC_027629 | Propionibacterium phage Attacne, complete genome | 28876 | Viruses; Duplodnaviria; Heunggongvirae; Uroviricota; Caudoviricetes; Caudovirales; Siphoviridae; Pahexavirus |
| NC_027630 | Propionibacterium phage Ouroboros, complete genome | 29506 | Viruses; Duplodnaviria; Heunggongvirae; Uroviricota; Caudoviricetes; Caudovirales; Siphoviridae; Pahexavirus |
| NC_028694 | Propionibacterium phage PA1-14, complete genome | 29407 | Viruses; Duplodnaviria; Heunggongvirae; Uroviricota; Caudoviricetes; Caudovirales; Siphoviridae; Pahexavirus; unclassified Pahexavirus |
| NC_028967 | Propionibacterium phage PAC1, complete genome | 29605 | Viruses; Duplodnaviria; Heunggongvirae; Uroviricota; Caudoviricetes; Caudovirales; Siphoviridae; Pahexavirus; unclassified Pahexavirus |
| NC_031003 | Propionibacterium phage Moyashi, complete genome | 29254 | Viruses; Duplodnaviria; Heunggongvirae; Uroviricota; Caudoviricetes; Caudovirales; Siphoviridae; Pahexavirus; unclassified Pahexavirus |
| NC_031005 | Propionibacterium phage QueenBey, complete genome | 29338 | Viruses; Duplodnaviria; Heunggongvirae; Uroviricota; Caudoviricetes; Caudovirales; Siphoviridae; Pahexavirus; unclassified Pahexavirus |
| NC_031084 | Propionibacterium phage BruceLethal, complete genome | 29249 | Viruses; Duplodnaviria; Heunggongvirae; Uroviricota; Caudoviricetes; Caudovirales; Siphoviridae; Pahexavirus; unclassified Pahexavirus |
| NC_031119 | Propionibacterium phage Enoki, complete genome | 29347 | Viruses; Duplodnaviria; Heunggongvirae; Uroviricota; Caudoviricetes; Caudovirales; Siphoviridae; Pahexavirus; unclassified Pahexavirus |
| NC_041954 | Propionibacterium phage Keiki, complete genome | 29339 | Viruses; Duplodnaviria; Heunggongvirae; Uroviricota; Caudoviricetes; Caudovirales; Siphoviridae; Pahexavirus |
| NC_041955 | Propionibacterium phage PHL082M03, complete genome | 29491 | Viruses; Duplodnaviria; Heunggongvirae; Uroviricota; Caudoviricetes; Caudovirales; Siphoviridae; Pahexavirus |
| NC_041956 | Propionibacterium phage PHL117M01, complete genome | 29422 | Viruses; Duplodnaviria; Heunggongvirae; Uroviricota; Caudoviricetes; Caudovirales; Siphoviridae; Pahexavirus |
| NC_041957 | Propionibacterium phage PHL151N00, complete genome | 29511 | Viruses; Duplodnaviria; Heunggongvirae; Uroviricota; Caudoviricetes; Caudovirales; Siphoviridae; Pahexavirus |

| # | Fd1 | Fd2 | Fd3 | PAVL20 | PAVL21 | PAVL33 | PAVL34 | PAVL45 | Erythromycin | Clindamycin | Tetracycline | Doxycycline | Minocycline |
|---|---|---|---|---|---|---|---|---|---|---|---|---|---|
| **1** | S | S | S | S | S | S | S | S | S | S | S | S | S |
| **2** | S | S | S | S | S | S | S | S | R | R | I | R | I |
| **4** | R | R | R | R | R | R | R | R | S | S | S | S | S |
| **5** | S | S | S | S | S | S | S | S | R | R | R | R | S |
| **6** | S | S | S | S | S | S | S | S | S | S | S | S | S |
| **7** | S | S | S | S | S | S | S | S | R | I | I | I | S |
| **8** | S | S | S | S | S | S | S | S | S | S | S | S | S |
| **9** | S | S | S | S | S | S | S | S | S | S | S | S | S |
| **10** | S | S | S | S | S | S | S | S | S | S | S | S | R |
| **11** | S | S | S | S | S | S | S | S | S | S | S | S | S |
| **13** | S | S | S | S | S | S | S | S | S | S | I | S | S |
| **14** | S | S | S | S | S | S | S | S | R | R | R | R | R |
| **15** | S | S | S | S | S | S | S | S | S | S | S | S | S |
| **18** | S | S | S | S | S | S | S | S | S | S | S | S | S |
| **19** | S | S | S | S | S | S | S | S | S | S | S | I | S |
| **21** | R | R | R | R | R | R | R | R | S | S | S | S | S |
| **22** | S | S | S | S | S | S | S | S | S | S | S | S | S |
| **23** | S | S | S | S | S | S | S | S | S | S | S | S | R |
| **24** | S | S | S | S | S | S | S | S | R | I | S | S | S |
| **25** | S | S | S | S | S | S | S | S | R | I | I | R | I |
| **27** | S | S | S | S | S | S | S | S | S | S | S | S | S |
| **28** | S | S | S | S | S | S | S | S | R | R | I | R | S |
| **30** | S | S | S | S | S | S | S | S | S | S | S | S | S |
| **31** | S | S | S | S | S | S | S | S | R | R | I | R | I |
| **32** | S | S | S | S | S | S | S | S | S | S | S | S | S |
| **33** | R | R | R | R | R | R | R | R | S | S | S | S | S |
| **35** | S | S | S | S | S | S | S | S | R | R | R | R | R |
| **36** | S | S | S | S | S | S | S | S | S | S | S | S | S |
| **37** | S | S | S | S | S | S | S | S | S | S | S | S | S |
| **40** | S | S | S | S | S | S | S | S | S | S | S | S | S |
| **41** | S | S | S | S | S | S | S | S | S | S | S | S | S |
| **43** | S | S | S | S | S | S | S | S | S | S | S | S | S |
| **44** | S | S | S | S | S | S | S | S | S | S | S | S | S |
| **47** | S | S | S | S | S | S | S | S | S | S | S | S | S |
| **48** | S | R | R | R | R | R | R | R | S | S | S | S | S |
| **49** | S | S | S | S | S | S | S | S | S | S | S | S | S |

### Control

### Animal : 1

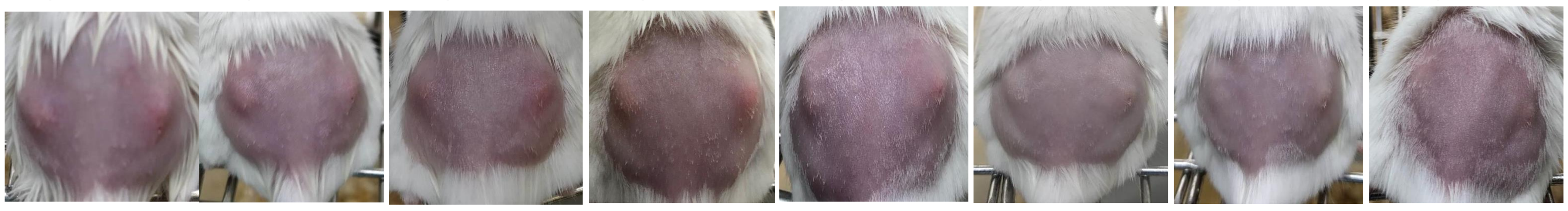

| | | Day 1 | | Day 2 | | Day 3 | | Day 4 | | Day 5 | | Day 6 | | Day 7 | | Day 8 | |
|---|---|---|---|---|---|---|---|---|---|---|---|---|---|---|---|---|---|
| | | L | R | L | R | L | R | L | R | L | R | L | R | L | R | L | R |
| **Diameter Score** | measured | 2 | 2 | 1 | 1 | 2 | 2 | 2 | 2 | 2 | 2 | 2 | 2 | 2 | 1 | 1 | 2 |
| **Elevation Score** | first scorer | 1 | 1 | 1 | 1 | 1 | 1 | 2 | 2 | 1 | 1 | 1 | 1 | 1 | 1 | 0 | 1 |
| | second scorer | 2 | 2 | 1 | 1 | 2 | 1 | 1 | 2 | 1 | 1 | 1 | 1 | 0 | 1 | 0 | 1 |
| **Eschar Score** | first scorer | 0 | 0 | 0 | 1 | 0 | 0 | 0 | 0 | 0 | 1 | 0 | 0 | 0 | 1 | 0 | 0 |
| | second scorer | 0 | 0 | 0 | 1 | 0 | 0 | 0 | 0 | 0 | 1 | 0 | 1 | 0 | 0 | 0 | 0 |
| **Total Score** | averaged | 3.5 | 3.5 | 2 | 3 | 3.5 | 3 | 3.5 | 4 | 3 | 4 | 3 | 3.5 | 2.5 | 2.5 | 1 | 3 |

### Animal : 2

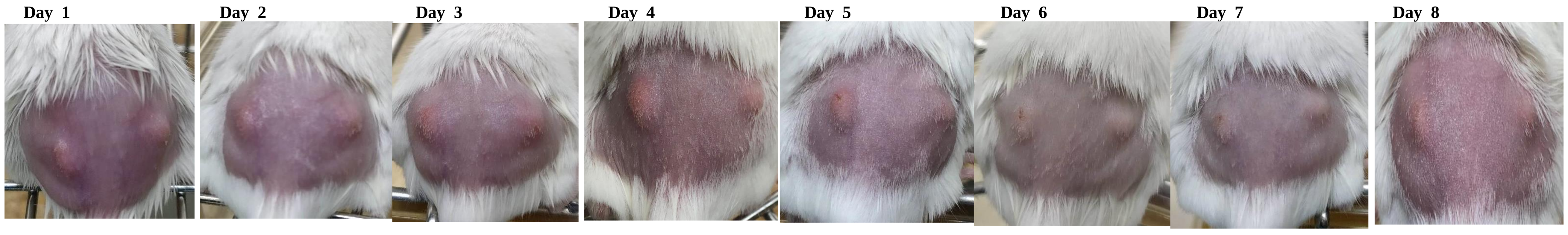

| | | Day 1 | | Day 2 | | Day 3 | | Day 4 | | Day 5 | | Day 6 | | Day 7 | | Day 8 | |
|---|---|---|---|---|---|---|---|---|---|---|---|---|---|---|---|---|---|
| | | L | R | L | R | L | R | L | R | L | R | L | R | L | R | L | R |
| **Diameter Score** | measured | 2 | 3 | 2 | 2 | 3 | 2 | 3 | 1 | 3 | 2 | 2 | 1 | 2 | 2 | 1 | 1 |
| **Elevation Score** | first scorer | 2 | 2 | 2 | 2 | 2 | 2 | 2 | 1 | 2 | 1 | 1 | 1 | 1 | 1 | 1 | 1 |
| | second scorer | 2 | 2 | 2 | 2 | 2 | 2 | 2 | 2 | 1 | 1 | 1 | 1 | 2 | 1 | 2 | 1 |
| **Eschar Score** | first scorer | 0 | 0 | 0 | 0 | 0 | 0 | 0 | 0 | 1 | 0 | 1 | 0 | 1 | 0 | 0 | 0 |
| | second scorer | 0 | 0 | 0 | 0 | 0 | 0 | 0 | 0 | 1 | 0 | 1 | 0 | 1 | 0 | 0 | 0 |
| **Total Score** | averaged | 4 | 5 | 4 | 4 | 5 | 4 | 5 | 2.5 | 5.5 | 3 | 4 | 2 | 4.5 | 3 | 2.5 | 2 |

### Animal : 3

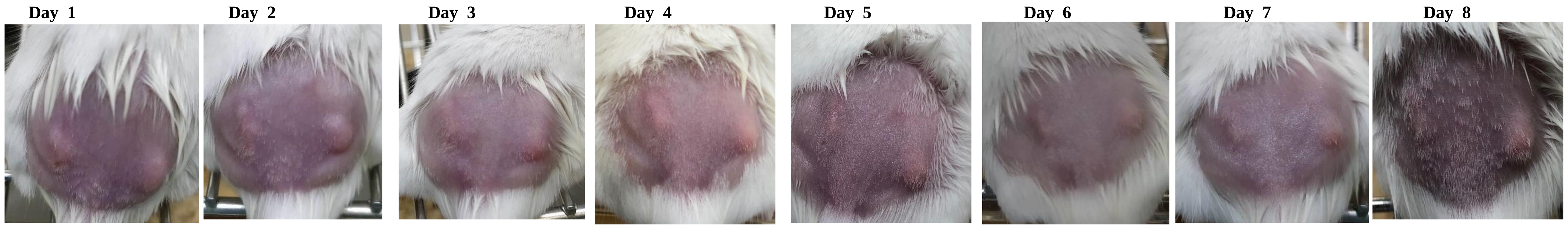

| | | Day 1 | | Day 2 | | Day 3 | | Day 4 | | Day 5 | | Day 6 | | Day 7 | | Day 8 | |
|---|---|---|---|---|---|---|---|---|---|---|---|---|---|---|---|---|---|
| | | L | R | L | R | L | R | L | R | L | R | L | R | L | R | L | R |
| **Diameter Score** | measured | 2 | 2 | 2 | 2 | 2 | 2 | 2 | 2 | 2 | 3 | 2 | 2 | 2 | 1 | 1 | 2 |
| **Elevation Score** | first scorer | 1 | 2 | 1 | 2 | 1 | 2 | 1 | 2 | 0 | 1 | 1 | 1 | 1 | 1 | 1 | 1 |
| | second scorer | 1 | 2 | 1 | 2 | 1 | 2 | 1 | 2 | 0 | 2 | 1 | 2 | 2 | 2 | 0 | 2 |
| **Eschar Score** | first scorer | 0 | 0 | 0 | 0 | 0 | 0 | 0 | 0 | 1 | 0 | 0 | 0 | 0 | 0 | 0 | 0 |
| | second scorer | 0 | 0 | 0 | 0 | 0 | 0 | 0 | 0 | 1 | 0 | 0 | 0 | 0 | 0 | 0 | 0 |
| **Total Score** | averaged | 3 | 4 | 3 | 4 | 3 | 4 | 3 | 4 | 3 | 4.5 | 3 | 3.5 | 3.5 | 2.5 | 1.5 | 3.5 |

### Animal : 4

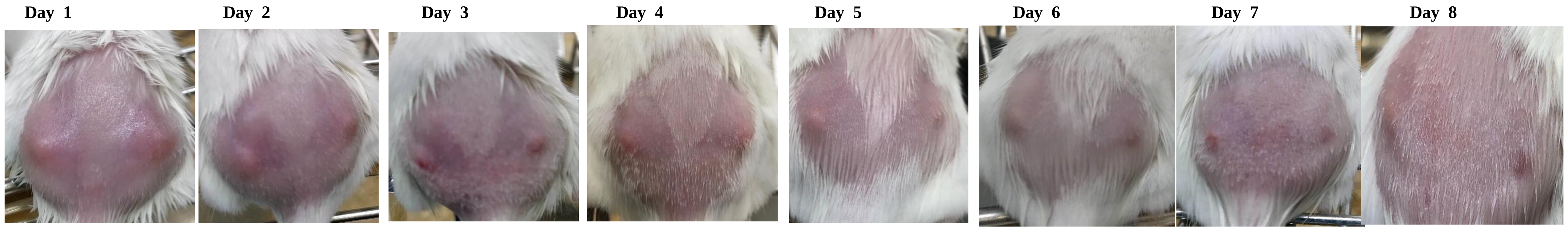

| | | Day 1 | | Day 2 | | Day 3 | | Day 4 | | Day 5 | | Day 6 | | Day 7 | | Day 8 | |
|---|---|---|---|---|---|---|---|---|---|---|---|---|---|---|---|---|---|
| | | L | R | L | R | L | R | L | R | L | R | L | R | L | R | L | R |
| **Diameter Score** | measured | 2 | 2 | 2 | 2 | 2 | 2 | 2 | 2 | 2 | 2 | 2 | 2 | 2 | 2 | 1 | 1 |
| **Elevation Score** | first scorer | 2 | 2 | 1 | 1 | 1 | 1 | 1 | 1 | 1 | 1 | 1 | 1 | 0 | 1 | 0 | 0 |
| | second scorer | 2 | 1 | 1 | 1 | 1 | 1 | 1 | 1 | 1 | 1 | 1 | 1 | 1 | 0 | 1 | 1 |
| **Eschar Score** | first scorer | 0 | 0 | 0 | 0 | 0 | 0 | 0 | 0 | 0 | 0 | 0 | 0 | 0 | 0 | 0 | 0 |
| | second scorer | 0 | 0 | 0 | 0 | 0 | 0 | 0 | 0 | 0 | 0 | 0 | 0 | 0 | 0 | 0 | 0 |
| **Total Score** | averaged | 4 | 3.5 | 3 | 3 | 3 | 3 | 3 | 3 | 3 | 3 | 3 | 3 | 2.5 | 2.5 | 1.5 | 1.5 |

### Animal : 5

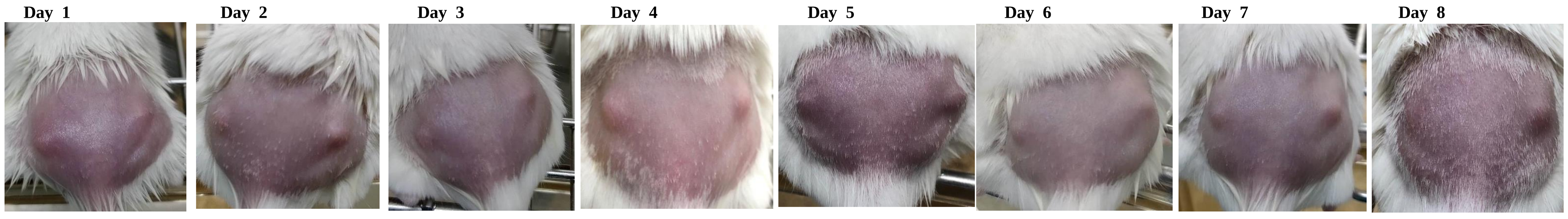

| | | Day 1 | | Day 2 | | Day 3 | | Day 4 | | Day 5 | | Day 6 | | Day 7 | | Day 8 | |
|---|---|---|---|---|---|---|---|---|---|---|---|---|---|---|---|---|---|
| | | L | R | L | R | L | R | L | R | L | R | L | R | L | R | L | R |
| **Diameter Score** | measured | 2 | 2 | 2 | 2 | 1 | 1 | 1 | 1 | 2 | 2 | 2 | 2 | 2 | 2 | 2 | 2 |
| **Elevation Score** | first scorer | 1 | 1 | 1 | 2 | 1 | 2 | 2 | 2 | 1 | 2 | 1 | 1 | 2 | 2 | 1 | 1 |
| | second scorer | 2 | 1 | 1 | 1 | 1 | 1 | 2 | 2 | 2 | 1 | 2 | 2 | 1 | 1 | 1 | 2 |
| **Eschar Score** | first scorer | 0 | 0 | 0 | 0 | 0 | 0 | 0 | 0 | 0 | 0 | 0 | 0 | 0 | 0 | 0 | 0 |
| | second scorer | 0 | 0 | 0 | 0 | 0 | 0 | 0 | 0 | 0 | 0 | 0 | 0 | 0 | 0 | 0 | 0 |
| **Total Score** | averaged | 3.5 | 3 | 3 | 3.5 | 2 | 2.5 | 3 | 3 | 3.5 | 3.5 | 3.5 | 3.5 | 3.5 | 3.5 | 3 | 3.5 |

**# Animal :** **6**

Day 1 | Day 2 | Day 3 | Day 4 | Day 5 | Day 6 | Day 7 | Day 8

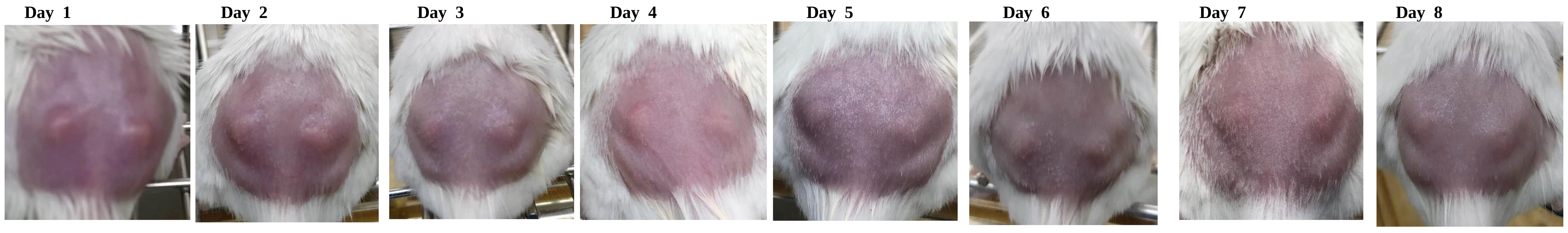

| | | L | R | L | R | L | R | L | R | L | R | L | R | L | R | L | R |
|---|---|---|---|---|---|---|---|---|---|---|---|---|---|---|---|---|---|
| **Diameter Score** | measured | 1 | 1 | 2 | 1 | 1 | 1 | 1 | 1 | 2 | 2 | 2 | 1 | 1 | 1 | 2 | 2 |
| **Elevation Score** | first scorer | 1 | 2 | 2 | 2 | 2 | 2 | 2 | 2 | 2 | 1 | 2 | 3 | 1 | 2 | 1 | 1 |
| | second scorer | 2 | 1 | 2 | 2 | 2 | 2 | 2 | 2 | 1 | 1 | 2 | 2 | 1 | 2 | 2 | 2 |
| **Eschar Score** | first scorer | 0 | 0 | 0 | 0 | 0 | 0 | 0 | 0 | 0 | 0 | 0 | 1 | 0 | 1 | 0 | 1 |
| | second scorer | 0 | 0 | 0 | 0 | 0 | 0 | 0 | 0 | 0 | 0 | 0 | 0 | 0 | 0 | 0 | 1 |
| **Total Score** | averaged | **2.5** | **2.5** | **4** | **3** | **3** | **3** | **3** | **3** | **3.5** | **3** | **4** | **4** | **2** | **3.5** | **3.5** | **4.5** |

**# Animal :** **7**

Day 1 | Day 2 | Day 3 | Day 4 | Day 5 | Day 6 | Day 7 | Day 8

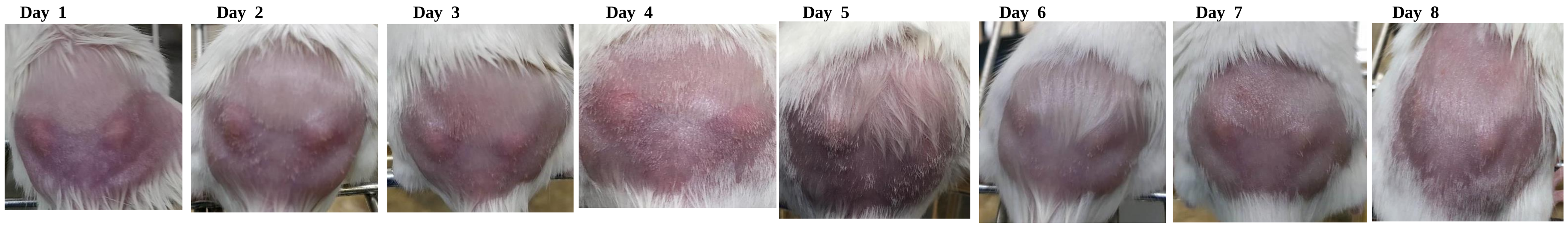

| | | L | R | L | R | L | R | L | R | L | R | L | R | L | R | L | R |
|---|---|---|---|---|---|---|---|---|---|---|---|---|---|---|---|---|---|
| **Diameter Score** | measured | 2 | 1 | 3 | 2 | 3 | 2 | 3 | 3 | 2 | 1 | 2 | 2 | 1 | 2 | 1 | 1 |
| **Elevation Score** | first scorer | 1 | 1 | 1 | 2 | 2 | 1 | 1 | 0 | 2 | 1 | 1 | 1 | 1 | 1 | 1 | 1 |
| | second scorer | 1 | 2 | 1 | 2 | 2 | 1 | 0 | 1 | 2 | 0 | 1 | 2 | 2 | 2 | 1 | 2 |
| **Eschar Score** | first scorer | 0 | 0 | 0 | 0 | 0 | 0 | 1 | 0 | 0 | 0 | 0 | 0 | 0 | 0 | 1 | 0 |
| | second scorer | 0 | 0 | 0 | 0 | 0 | 0 | 0 | 0 | 0 | 0 | 0 | 0 | 0 | 0 | 0 | 0 |
| **Total Score** | averaged | **3** | **2.5** | **4** | **4** | **5** | **3** | **4** | **3.5** | **4** | **1.5** | **3** | **3.5** | **2.5** | **3.5** | **2.5** | **2.5** |

**# Animal :** **8**

Day 1 | Day 2 | Day 3 | Day 4 | Day 5 | Day 6 | Day 7 | Day 8

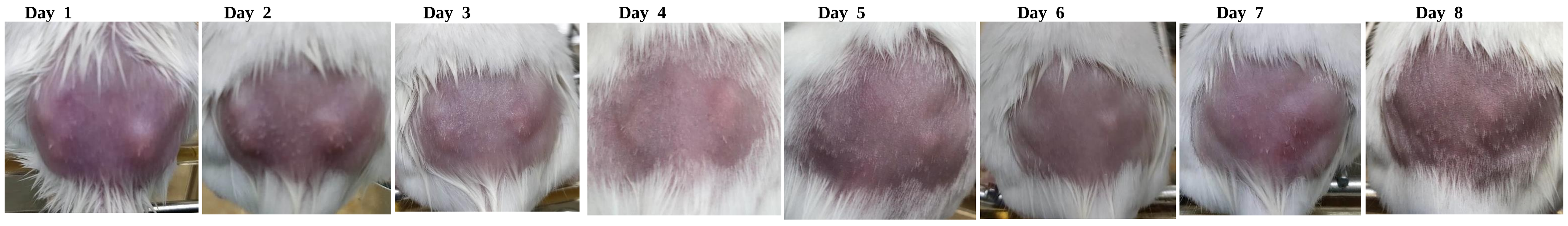

| | | L | R | L | R | L | R | L | R | L | R | L | R | L | R | L | R |
|---|---|---|---|---|---|---|---|---|---|---|---|---|---|---|---|---|---|
| **Diameter Score** | measured | 2 | 2 | 2 | 2 | 3 | 3 | 3 | 2 | 2 | 1 | 2 | 2 | 2 | 2 | 2 | 2 |
| **Elevation Score** | first scorer | 1 | 2 | 2 | 2 | 1 | 1 | 1 | 1 | 0 | 1 | 0 | 1 | 1 | 1 | 1 | 1 |
| | second scorer | 1 | 2 | 2 | 2 | 1 | 1 | 0 | 1 | 0 | 1 | 1 | 2 | 1 | 2 | 1 | 2 |
| **Eschar Score** | first scorer | 0 | 0 | 0 | 0 | 0 | 0 | 0 | 0 | 0 | 0 | 0 | 0 | 0 | 0 | 0 | 0 |
| | second scorer | 0 | 0 | 0 | 0 | 0 | 0 | 0 | 0 | 0 | 0 | 0 | 0 | 0 | 0 | 0 | 0 |
| **Total Score** | averaged | **3** | **4** | **4** | **4** | **4** | **4** | **3.5** | **3** | **2** | **2** | **2.5** | **3.5** | **3** | **3.5** | **3** | **3.5** |

**# Animal :** **9**

Day 1 | Day 2 | Day 3 | Day 4 | Day 5 | Day 6 | Day 7 | Day 8

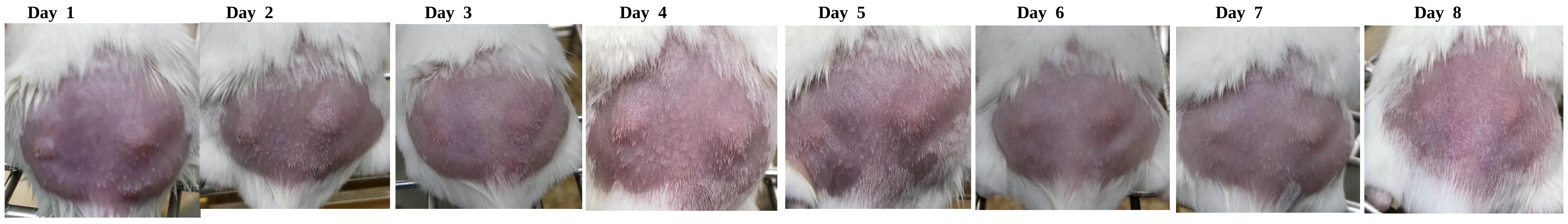

| | | L | R | L | R | L | R | L | R | L | R | L | R | L | R | L | R |
|---|---|---|---|---|---|---|---|---|---|---|---|---|---|---|---|---|---|
| **Diameter Score** | measured | 1 | 2 | 1 | 2 | 2 | 2 | 2 | 3 | 1 | 2 | 2 | 1 | 2 | 2 | 2 | 1 |
| **Elevation Score** | first scorer | 1 | 2 | 1 | 2 | 0 | 0 | 2 | 1 | 0 | 2 | 1 | 1 | 1 | 1 | 1 | 1 |
| | second scorer | 2 | 2 | 2 | 2 | 0 | 0 | 2 | 1 | 0 | 1 | 1 | 1 | 1 | 2 | 1 | 2 |
| **Eschar Score** | first scorer | 0 | 0 | 0 | 0 | 0 | 0 | 0 | 0 | 0 | 0 | 0 | 0 | 0 | 0 | 0 | 0 |
| | second scorer | 0 | 0 | 0 | 0 | 0 | 0 | 0 | 0 | 0 | 0 | 0 | 0 | 0 | 0 | 0 | 0 |
| **Total Score** | averaged | **2.5** | **4** | **2.5** | **4** | **2** | **2** | **4** | **4** | **1** | **3.5** | **3** | **2** | **3** | **3.5** | **3** | **2.5** |

**# Animal :** **10**

Day 1 | Day 2 | Day 3 | Day 4 | Day 5 | Day 6 | Day 7 | Day 8

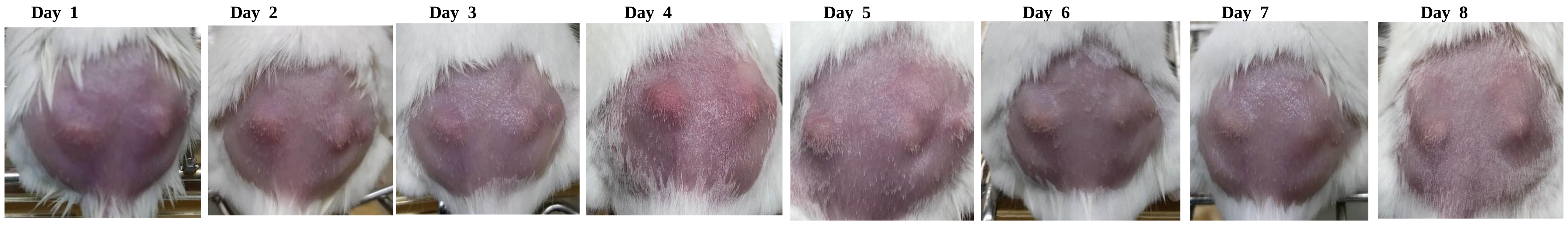

| | | L | R | L | R | L | R | L | R | L | R | L | R | L | R | L | R |
|---|---|---|---|---|---|---|---|---|---|---|---|---|---|---|---|---|---|
| **Diameter Score** | measured | 2 | 2 | 2 | 1 | 2 | 1 | 2 | 2 | 1 | 1 | 2 | 2 | 2 | 2 | 2 | 2 |
| **Elevation Score** | first scorer | 2 | 2 | 2 | 2 | 1 | 2 | 1 | 1 | 2 | 2 | 2 | 2 | 2 | 2 | 2 | 2 |
| | second scorer | 2 | 2 | 2 | 2 | 2 | 2 | 1 | 1 | 2 | 2 | 2 | 2 | 2 | 2 | 2 | 2 |
| **Eschar Score** | first scorer | 0 | 0 | 0 | 0 | 0 | 0 | 0 | 0 | 0 | 0 | 0 | 0 | 0 | 0 | 1 | 0 |
| | second scorer | 0 | 0 | 0 | 0 | 0 | 0 | 0 | 0 | 0 | 0 | 0 | 0 | 0 | 0 | 0 | 0 |
| **Total Score** | averaged | **4** | **4** | **4** | **3** | **3.5** | **3** | **3** | **3** | **3** | **3** | **4** | **4** | **4** | **4** | **4.5** | **4** |

Day 1 | Day 2 | Day 3 | Day 4 | Day 5 | Day 6 | Day 7 | Day 8

**# Animal :** **11**

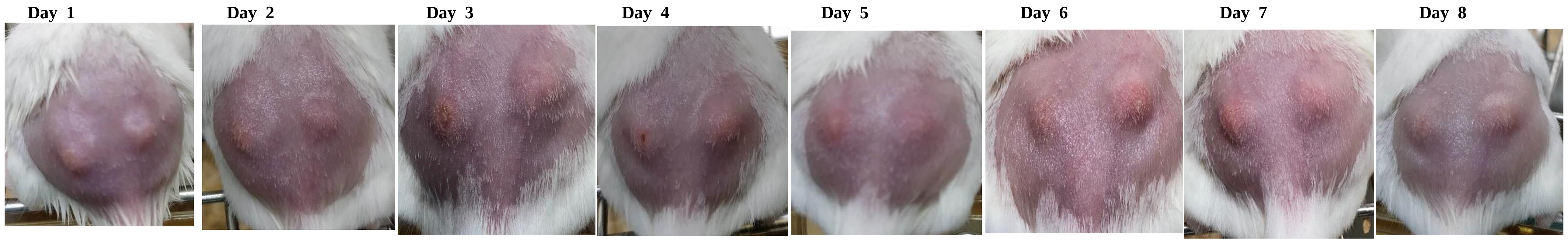

| | | Day 1 | | Day 2 | | Day 3 | | Day 4 | | Day 5 | | Day 6 | | Day 7 | | Day 8 | |
|---|---|---|---|---|---|---|---|---|---|---|---|---|---|---|---|---|---|
| | | **L** | **R** | **L** | **R** | **L** | **R** | **L** | **R** | **L** | **R** | **L** | **R** | **L** | **R** | **L** | **R** |
| **Diameter Score** | measured | 2 | 2 | 2 | 2 | 2 | 2 | 2 | 3 | 2 | 2 | 2 | 2 | 2 | 2 | 2 | 3 |
| **Elevation Score** | first scorer | 2 | 3 | 1 | 2 | 2 | 2 | 2 | 3 | 2 | 2 | 2 | 2 | 2 | 2 | 2 | 3 |
| | second scorer | 3 | 3 | 2 | 1 | 2 | 2 | 2 | 3 | 1 | 2 | 1 | 2 | 2 | 2 | 2 | 2 |
| **Eschar Score** | first scorer | 0 | 0 | 0 | 0 | 1 | 0 | 1 | 0 | 0 | 0 | 0 | 0 | 0 | 0 | 1 | 0 |
| | second scorer | 0 | 0 | 0 | 0 | 1 | 0 | 1 | 0 | 0 | 0 | 0 | 0 | 0 | 0 | 0 | 0 |
| **Total Score** | averaged | **4.5** | **5** | **3.5** | **3.5** | **5** | **4** | **5** | **6** | **3.5** | **4** | **3.5** | **4** | **4** | **4** | **4.5** | **5.5** |

Day 1 | Day 2 | Day 3 | Day 4 | Day 5 | Day 6 | Day 7 | Day 8

**# Animal :** **12**

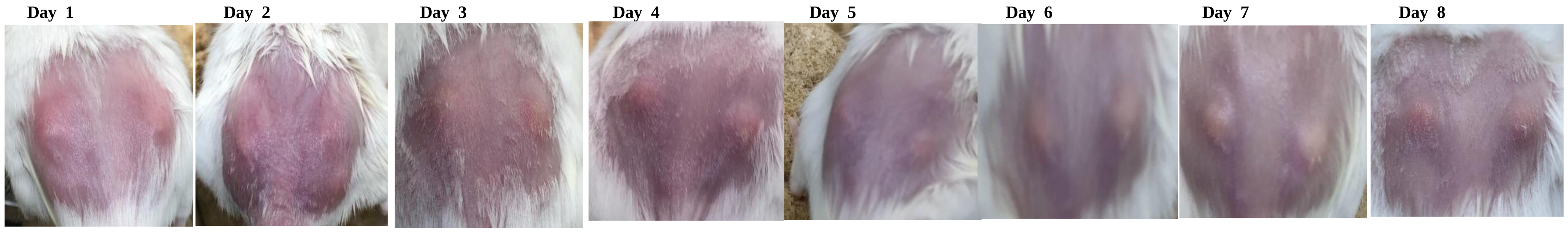

| | | Day 1 | | Day 2 | | Day 3 | | Day 4 | | Day 5 | | Day 6 | | Day 7 | | Day 8 | |
|---|---|---|---|---|---|---|---|---|---|---|---|---|---|---|---|---|---|
| | | **L** | **R** | **L** | **R** | **L** | **R** | **L** | **R** | **L** | **R** | **L** | **R** | **L** | **R** | **L** | **R** |
| **Diameter Score** | measured | 2 | 2 | 3 | 2 | 3 | 2 | 2 | 2 | 2 | 2 | 2 | 2 | 2 | 2 | 2 | 2 |
| **Elevation Score** | first scorer | 2 | 1 | 1 | 2 | 1 | 2 | 2 | 2 | 2 | 1 | 1 | 1 | 3 | 2 | 2 | 1 |
| | second scorer | 1 | 1 | 1 | 2 | 1 | 2 | 1 | 2 | 2 | 2 | 2 | 2 | 2 | 1 | 2 | 1 |
| **Eschar Score** | first scorer | 0 | 0 | 0 | 0 | 0 | 0 | 0 | 0 | 0 | 0 | 0 | 0 | 0 | 0 | 0 | 0 |
| | second scorer | 0 | 0 | 0 | 0 | 0 | 0 | 0 | 0 | 0 | 0 | 0 | 0 | 0 | 0 | 0 | 0 |
| **Total Score** | averaged | **3.5** | **3** | **4** | **4** | **4** | **4** | **3.5** | **4** | **4** | **3.5** | **3.5** | **3.5** | **4.5** | **3.5** | **4** | **3** |

Day 1 | Day 2 | Day 3 | Day 4 | Day 5 | Day 6 | Day 7 | Day 8

**# Animal :** **13**

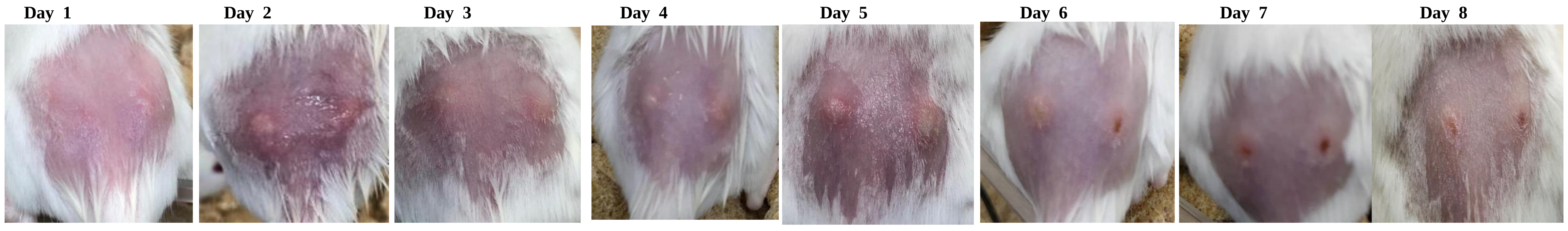

| | | Day 1 | | Day 2 | | Day 3 | | Day 4 | | Day 5 | | Day 6 | | Day 7 | | Day 8 | |
|---|---|---|---|---|---|---|---|---|---|---|---|---|---|---|---|---|---|
| | | **L** | **R** | **L** | **R** | **L** | **R** | **L** | **R** | **L** | **R** | **L** | **R** | **L** | **R** | **L** | **R** |
| **Diameter Score** | measured | 2 | 2 | 2 | 2 | 2 | 2 | 2 | 2 | 2 | 2 | 2 | 2 | 2 | 2 | 1 | 2 |
| **Elevation Score** | first scorer | 2 | 2 | 2 | 1 | 2 | 2 | 2 | 1 | 2 | 2 | 2 | 1 | 1 | 1 | 1 | 1 |
| | second scorer | 2 | 1 | 1 | 1 | 2 | 2 | 1 | 1 | 3 | 2 | 2 | 1 | 1 | 1 | 1 | 1 |
| **Eschar Score** | first scorer | 0 | 0 | 0 | 0 | 0 | 0 | 0 | 0 | 0 | 0 | 0 | 1 | 1 | 2 | 1 | 2 |
| | second scorer | 0 | 0 | 0 | 0 | 0 | 0 | 0 | 0 | 0 | 0 | 0 | 1 | 1 | 2 | 1 | 2 |
| **Total Score** | averaged | **4** | **3.5** | **3.5** | **3** | **4** | **4** | **3.5** | **3** | **4.5** | **4** | **4** | **4** | **4** | **5** | **3** | **5** |

Day 1 | Day 2 | Day 3 | Day 4 | Day 5 | Day 6 | Day 7 | Day 8

**# Animal :** **14**

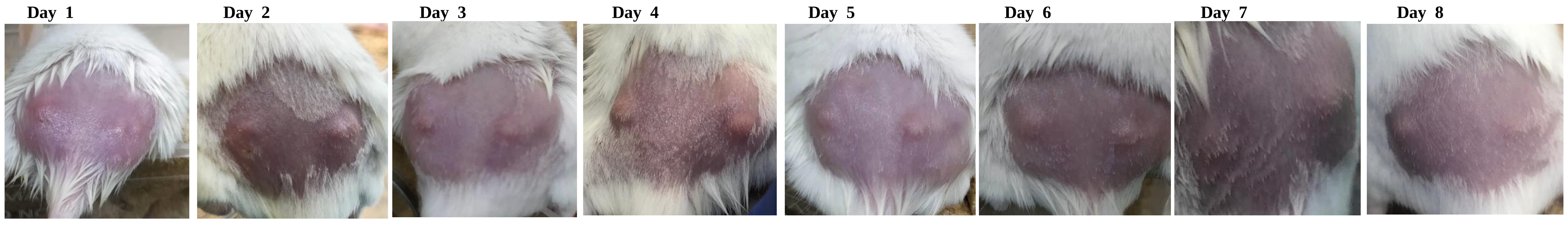

| | | Day 1 | | Day 2 | | Day 3 | | Day 4 | | Day 5 | | Day 6 | | Day 7 | | Day 8 | |
|---|---|---|---|---|---|---|---|---|---|---|---|---|---|---|---|---|---|
| | | **L** | **R** | **L** | **R** | **L** | **R** | **L** | **R** | **L** | **R** | **L** | **R** | **L** | **R** | **L** | **R** |
| **Diameter Score** | measured | 2 | 2 | 2 | 2 | 2 | 3 | 2 | 3 | 2 | 3 | 3 | 3 | 2 | 3 | 2 | 3 |
| **Elevation Score** | first scorer | 0 | 0 | 1 | 1 | 1 | 1 | 2 | 1 | 1 | 1 | 2 | 1 | 0 | 1 | 1 | 0 |
| | second scorer | 0 | 1 | 1 | 2 | 1 | 1 | 1 | 2 | 2 | 2 | 1 | 1 | 1 | 1 | 1 | 1 |
| **Eschar Score** | first scorer | 0 | 0 | 0 | 0 | 0 | 0 | 0 | 0 | 0 | 0 | 0 | 0 | 0 | 0 | 0 | 0 |
| | second scorer | 0 | 0 | 0 | 0 | 0 | 0 | 0 | 0 | 0 | 0 | 0 | 0 | 0 | 0 | 0 | 0 |
| **Total Score** | averaged | **2** | **2.5** | **3** | **3.5** | **3** | **4** | **3.5** | **4.5** | **3.5** | **4.5** | **4.5** | **4** | **2.5** | **4** | **3** | **3.5** |

Day 1 | Day 2 | Day 3 | Day 4 | Day 5 | Day 6 | Day 7 | Day 8

**# Animal :** **15**

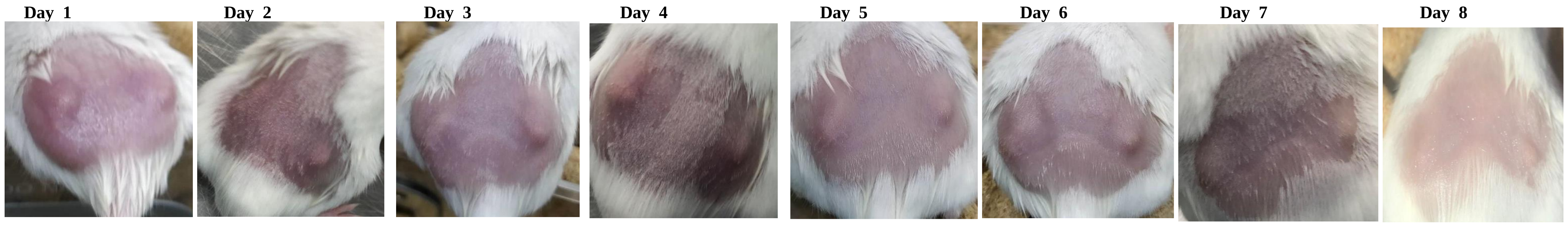

| | | Day 1 | | Day 2 | | Day 3 | | Day 4 | | Day 5 | | Day 6 | | Day 7 | | Day 8 | |
|---|---|---|---|---|---|---|---|---|---|---|---|---|---|---|---|---|---|
| | | **L** | **R** | **L** | **R** | **L** | **R** | **L** | **R** | **L** | **R** | **L** | **R** | **L** | **R** | **L** | **R** |
| **Diameter Score** | measured | 1 | 2 | 2 | 2 | 2 | 2 | 2 | 2 | 1 | 1 | 1 | 2 | 1 | 2 | 1 | 2 |
| **Elevation Score** | first scorer | 2 | 2 | 1 | 1 | 1 | 2 | 1 | 1 | 2 | 1 | 2 | 1 | 1 | 1 | 1 | 1 |
| | second scorer | 2 | 2 | 0 | 1 | 1 | 2 | 2 | 1 | 2 | 2 | 2 | 2 | 1 | 1 | 1 | 1 |
| **Eschar Score** | first scorer | 0 | 0 | 0 | 0 | 0 | 0 | 0 | 0 | 0 | 0 | 0 | 0 | 0 | 0 | 0 | 0 |
| | second scorer | 0 | 0 | 0 | 0 | 0 | 0 | 0 | 0 | 0 | 0 | 0 | 0 | 0 | 0 | 0 | 0 |
| **Total Score** | averaged | **3** | **4** | **2.5** | **3** | **3** | **4** | **3.5** | **3** | **3** | **2.5** | **3** | **3.5** | **2** | **3** | **2** | **3** |

Day 1 | Day 2 | Day 3 | Day 4 | Day 5 | Day 6 | Day 7 | Day 8

**# Animal :** **16**

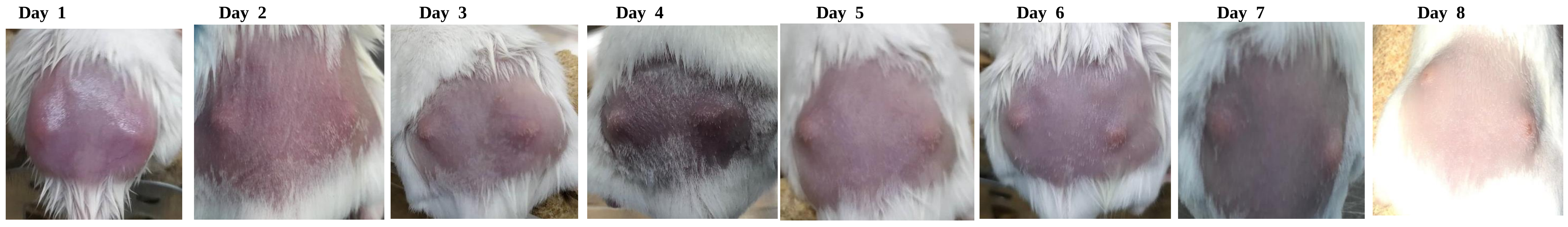

| | | L | R | L | R | L | R | L | R | L | R | L | R | L | R | L | R |
|---|---|---|---|---|---|---|---|---|---|---|---|---|---|---|---|---|---|
| **Diameter Score** | measured | 2 | 2 | 2 | 1 | 2 | 2 | 2 | 2 | 2 | 2 | 1 | 2 | 2 | 2 | 1 | 2 |
| **Elevation Score** | first scorer | 1 | 2 | 1 | 1 | 1 | 2 | 1 | 2 | 1 | 1 | 1 | 1 | 1 | 1 | 1 | 1 |
| | second scorer | 2 | 2 | 1 | 1 | 1 | 1 | 2 | 1 | 1 | 1 | 1 | 1 | 2 | 2 | 0 | 1 |
| **Eschar Score** | first scorer | 0 | 0 | 0 | 0 | 0 | 0 | 0 | 0 | 0 | 0 | 0 | 0 | 0 | 0 | 1 | 2 |
| | second scorer | 0 | 0 | 0 | 0 | 0 | 0 | 0 | 0 | 0 | 0 | 0 | 0 | 0 | 0 | 1 | 1 |
| **Total Score** | averaged | **3.5** | **4** | **3** | **2** | **3** | **3.5** | **3.5** | **3.5** | **3** | **3** | **2** | **3** | **3.5** | **3.5** | **2.5** | **4.5** |

Day 1 | Day 2 | Day 3 | Day 4 | Day 5 | Day 6 | Day 7 | Day 8

**# Animal :** **17**

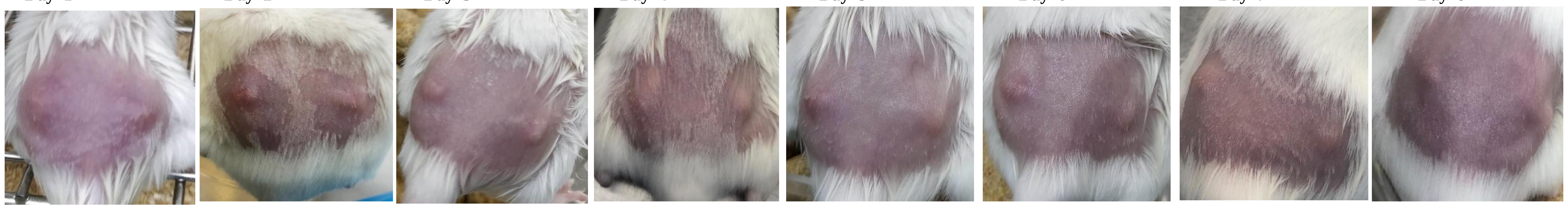

| | | L | R | L | R | L | R | L | R | L | R | L | R | L | R | L | R |
|---|---|---|---|---|---|---|---|---|---|---|---|---|---|---|---|---|---|
| **Diameter Score** | measured | 2 | 1 | 3 | 2 | 2 | 2 | 3 | 2 | 2 | 2 | 2 | 3 | 1 | 2 | 2 | 2 |
| **Elevation Score** | first scorer | 1 | 1 | 1 | 0 | 1 | 1 | 1 | 1 | 2 | 2 | 2 | 2 | 1 | 2 | 1 | 2 |
| | second scorer | 1 | 2 | 2 | 1 | 2 | 2 | 1 | 1 | 1 | 1 | 1 | 1 | 2 | 2 | 2 | 2 |
| **Eschar Score** | first scorer | 0 | 1 | 1 | 0 | 0 | 0 | 0 | 0 | 0 | 0 | 0 | 0 | 0 | 0 | 0 | 0 |
| | second scorer | 0 | 0 | 0 | 0 | 0 | 0 | 0 | 0 | 0 | 0 | 0 | 0 | 0 | 0 | 0 | 0 |
| **Total Score** | averaged | **3** | **3** | **5** | **2.5** | **3.5** | **3.5** | **4** | **3** | **3.5** | **3.5** | **3.5** | **4.5** | **2.5** | **4** | **3.5** | **4** |

### Phage-treated group

Day 1 | Day 2 | Day 3 | Day 4 | Day 5 | Day 6 | Day 7 | Day 8

### Animal : 1

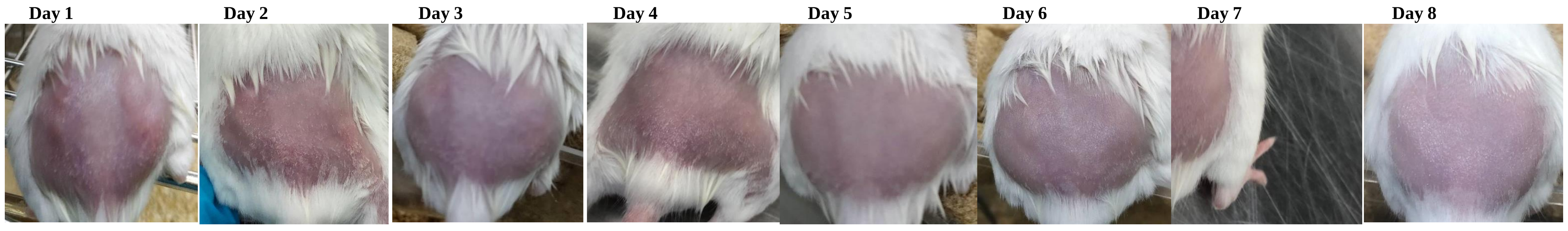

| | | Day 1 | | Day 2 | | Day 3 | | Day 4 | | Day 5 | | Day 6 | | Day 7 | | Day 8 | |
|---|---|---|---|---|---|---|---|---|---|---|---|---|---|---|---|---|---|
| | | **L** | **R** | **L** | **R** | **L** | **R** | **L** | **R** | **L** | **R** | **L** | **R** | **L** | **R** | **L** | **R** |
| **Diameter Score** | measured | 1 | 2 | 2 | 1 | 2 | 2 | 1 | 1 | 1 | 0 | 1 | 0 | 1 | 0 | 1 | 0 |
| **Elevation Score** | first scorer | 1 | 1 | 2 | 0 | 1 | 0 | 1 | 1 | 0 | 0 | 1 | 1 | 1 | 0 | 1 | 0 |
| | second scorer | 2 | 2 | 1 | 0 | 1 | 0 | 0 | 0 | 1 | 0 | 1 | 1 | 1 | 0 | 1 | 1 |
| **Eschar Score** | first scorer | 0 | 0 | 0 | 0 | 0 | 0 | 0 | 0 | 0 | 0 | 0 | 0 | 0 | 0 | 0 | 0 |
| | second scorer | 0 | 0 | 0 | 0 | 0 | 0 | 0 | 0 | 0 | 0 | 0 | 0 | 0 | 0 | 0 | 0 |
| **Total Score** | averaged | **2.5** | **3.5** | **3.5** | **1** | **3** | **2** | **1.5** | **1.5** | **1.5** | **0** | **2** | **1** | **2** | **0** | **2** | **0.5** |

### Animal : 2

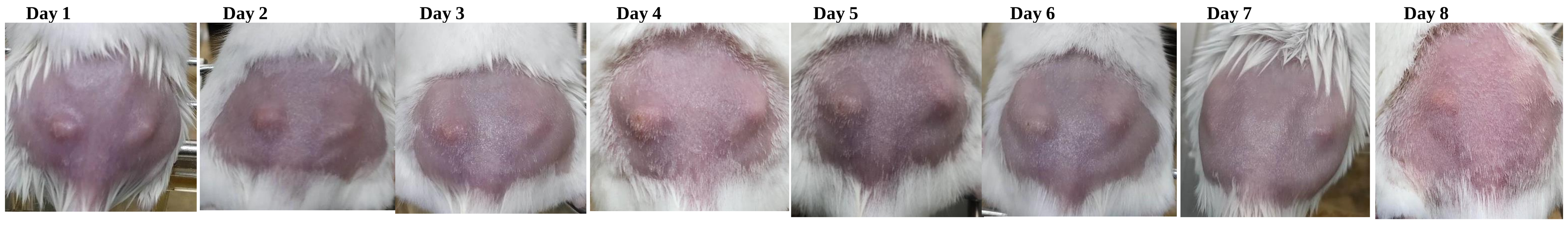

| | | Day 1 | | Day 2 | | Day 3 | | Day 4 | | Day 5 | | Day 6 | | Day 7 | | Day 8 | |
|---|---|---|---|---|---|---|---|---|---|---|---|---|---|---|---|---|---|
| | | **L** | **R** | **L** | **R** | **L** | **R** | **L** | **R** | **L** | **R** | **L** | **R** | **L** | **R** | **L** | **R** |
| **Diameter Score** | measured | 3 | 1 | 2 | 1 | 2 | 1 | 2 | 1 | 2 | 1 | 2 | 1 | 1 | 1 | 1 | 1 |
| **Elevation Score** | first scorer | 2 | 2 | 2 | 1 | 2 | 2 | 2 | 1 | 2 | 1 | 2 | 1 | 1 | 2 | 1 | 1 |
| | second scorer | 2 | 2 | 2 | 2 | 2 | 1 | 2 | 2 | 2 | 2 | 2 | 2 | 1 | 1 | 1 | 1 |
| **Eschar Score** | first scorer | 0 | 0 | 0 | 0 | 0 | 0 | 1 | 0 | 1 | 0 | 0 | 0 | 0 | 0 | 0 | 0 |
| | second scorer | 0 | 0 | 0 | 0 | 0 | 0 | 0 | 0 | 0 | 0 | 0 | 0 | 0 | 0 | 0 | 0 |
| **Total Score** | averaged | **5** | **3** | **4** | **2.5** | **4** | **2.5** | **4.5** | **2.5** | **4.5** | **2.5** | **4** | **2.5** | **2** | **2.5** | **2** | **2** |

### Animal : 3

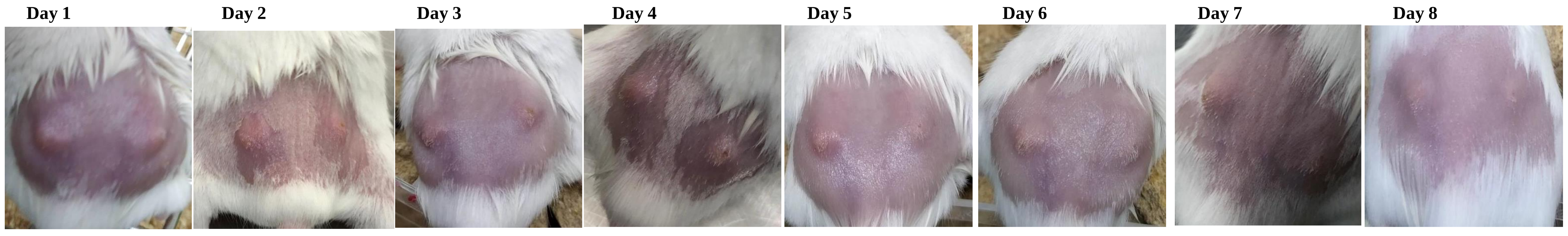

| | | Day 1 | | Day 2 | | Day 3 | | Day 4 | | Day 5 | | Day 6 | | Day 7 | | Day 8 | |
|---|---|---|---|---|---|---|---|---|---|---|---|---|---|---|---|---|---|
| | | **L** | **R** | **L** | **R** | **L** | **R** | **L** | **R** | **L** | **R** | **L** | **R** | **L** | **R** | **L** | **R** |
| **Diameter Score** | measured | 3 | 2 | 2 | 1 | 2 | 1 | 1 | 1 | 1 | 1 | 1 | 1 | 1 | 1 | 1 | 0 |
| **Elevation Score** | first scorer | 2 | 2 | 1 | 1 | 1 | 1 | 1 | 1 | 1 | 0 | 1 | 1 | 1 | 1 | 1 | 0 |
| | second scorer | 2 | 2 | 0 | 1 | 1 | 0 | 0 | 1 | 1 | 1 | 1 | 1 | 1 | 0 | 1 | 1 |
| **Eschar Score** | first scorer | 0 | 0 | 0 | 0 | 0 | 0 | 0 | 0 | 0 | 0 | 0 | 0 | 0 | 0 | 0 | 0 |
| | second scorer | 0 | 0 | 0 | 0 | 0 | 0 | 0 | 1 | 0 | 0 | 1 | 1 | 0 | 0 | 0 | 0 |
| **Total Score** | averaged | **5** | **4** | **2.5** | **2** | **3** | **1.5** | **1.5** | **2.5** | **2** | **1.5** | **2.5** | **2.5** | **2** | **1.5** | **2** | **0.5** |

### Animal : 4

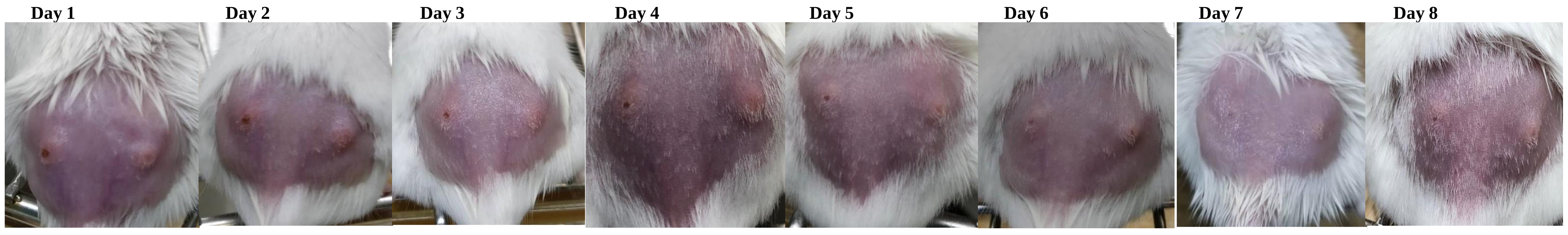

| | | Day 1 | | Day 2 | | Day 3 | | Day 4 | | Day 5 | | Day 6 | | Day 7 | | Day 8 | |
|---|---|---|---|---|---|---|---|---|---|---|---|---|---|---|---|---|---|
| | | **L** | **R** | **L** | **R** | **L** | **R** | **L** | **R** | **L** | **R** | **L** | **R** | **L** | **R** | **L** | **R** |
| **Diameter Score** | measured | 2 | 2 | 2 | 2 | 1 | 2 | 1 | 2 | 1 | 1 | 2 | 2 | 1 | 1 | 1 | 1 |
| **Elevation Score** | first scorer | 1 | 1 | 0 | 2 | 1 | 1 | 1 | 1 | 2 | 1 | 1 | 2 | 1 | 1 | 1 | 1 |
| | second scorer | 2 | 1 | 1 | 2 | 0 | 1 | 1 | 1 | 1 | 1 | 1 | 2 | 1 | 1 | 0 | 1 |
| **Eschar Score** | first scorer | 2 | 1 | 1 | 0 | 1 | 0 | 1 | 0 | 1 | 0 | 2 | 0 | 0 | 0 | 1 | 1 |
| | second scorer | 1 | 1 | 2 | 0 | 2 | 0 | 1 | 0 | 2 | 1 | 1 | 1 | 1 | 0 | 0 | 0 |
| **Total Score** | averaged | **5** | **4** | **4** | **4** | **3** | **3** | **3** | **3** | **4** | **2.5** | **4.5** | **4.5** | **2.5** | **2** | **2** | **2.5** |

### Animal : 5

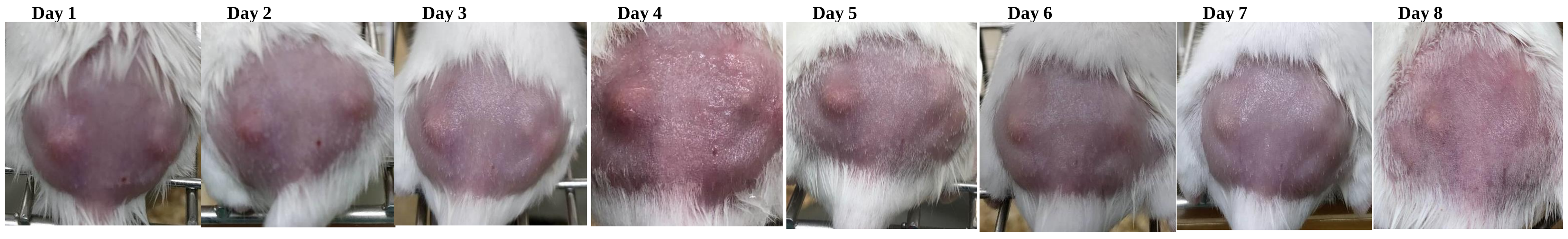

| | | Day 1 | | Day 2 | | Day 3 | | Day 4 | | Day 5 | | Day 6 | | Day 7 | | Day 8 | |
|---|---|---|---|---|---|---|---|---|---|---|---|---|---|---|---|---|---|
| | | **L** | **R** | **L** | **R** | **L** | **R** | **L** | **R** | **L** | **R** | **L** | **R** | **L** | **R** | **L** | **R** |
| **Diameter Score** | measured | 2 | 1 | 2 | 1 | 2 | 2 | 2 | 1 | 2 | 1 | 2 | 2 | 2 | 1 | 2 | 0 |
| **Elevation Score** | first scorer | 2 | 2 | 2 | 1 | 2 | 1 | 2 | 1 | 2 | 1 | 2 | 2 | 1 | 1 | 1 | 1 |
| | second scorer | 2 | 2 | 1 | 1 | 2 | 1 | 1 | 0 | 2 | 2 | 2 | 1 | 1 | 1 | 0 | 0 |
| **Eschar Score** | first scorer | 0 | 0 | 0 | 0 | 0 | 0 | 0 | 0 | 0 | 0 | 0 | 0 | 0 | 0 | 0 | 0 |
| | second scorer | 0 | 0 | 0 | 0 | 0 | 0 | 0 | 0 | 0 | 0 | 0 | 0 | 0 | 0 | 0 | 0 |
| **Total Score** | averaged | **4** | **3** | **3.5** | **2** | **4** | **3** | **3.5** | **1.5** | **4** | **2.5** | **4** | **3.5** | **3** | **2** | **2.5** | **0.5** |

### Animal : 6

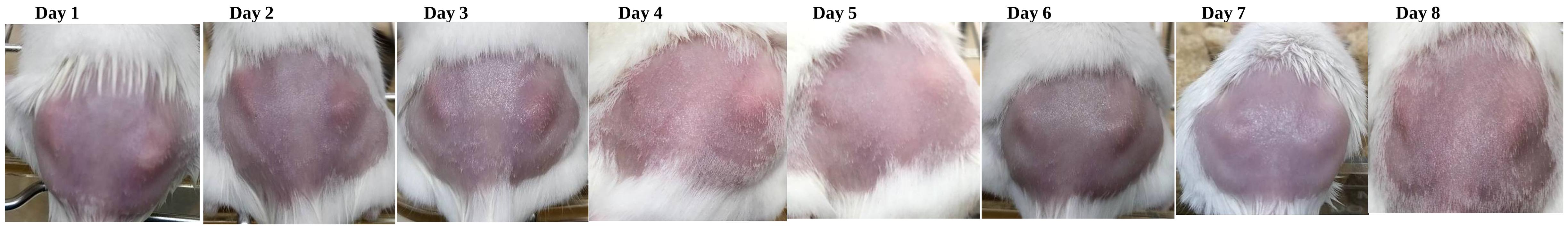

| | | Day 1 | | Day 2 | | Day 3 | | Day 4 | | Day 5 | | Day 6 | | Day 7 | | Day 8 | |
|---|---|---|---|---|---|---|---|---|---|---|---|---|---|---|---|---|---|
| | | **L** | **R** | **L** | **R** | **L** | **R** | **L** | **R** | **L** | **R** | **L** | **R** | **L** | **R** | **L** | **R** |
| **Diameter Score** | measured | 2 | 2 | 2 | 2 | 2 | 3 | 1 | 2 | 1 | 2 | 1 | 2 | 1 | 2 | 1 | 2 |
| **Elevation Score** | first scorer | 2 | 1 | 1 | 2 | 0 | 2 | 1 | 2 | 2 | 2 | 1 | 1 | 0 | 1 | 1 | 0 |
| | second scorer | 1 | 2 | 0 | 1 | 1 | 1 | 0 | 1 | 1 | 1 | 1 | 2 | 1 | 2 | 0 | 1 |
| **Eschar Score** | first scorer | 0 | 0 | 0 | 0 | 0 | 0 | 0 | 0 | 0 | 0 | 0 | 0 | 0 | 0 | 0 | 0 |
| | second scorer | 0 | 0 | 0 | 0 | 0 | 0 | 0 | 0 | 0 | 0 | 0 | 0 | 0 | 0 | 0 | 0 |
| **Total Score** | averaged | **3.5** | **3.5** | **2.5** | **3.5** | **2.5** | **4.5** | **1.5** | **3.5** | **2.5** | **3.5** | **2** | **3.5** | **1.5** | **3.5** | **1.5** | **2.5** |

Day 1 | Day 2 | Day 3 | Day 4 | Day 5 | Day 6 | Day 7 | Day 8

**# Animal :** **7**

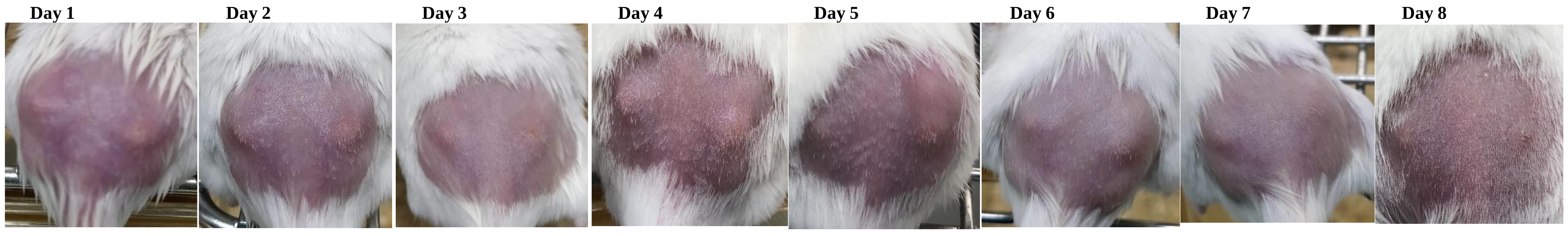

| | | Day 1 L | Day 1 R | Day 2 L | Day 2 R | Day 3 L | Day 3 R | Day 4 L | Day 4 R | Day 5 L | Day 5 R | Day 6 L | Day 6 R | Day 7 L | Day 7 R | Day 8 L | Day 8 R |
|---|---|---|---|---|---|---|---|---|---|---|---|---|---|---|---|---|---|
| **Diameter Score** | measured | 2 | 1 | 2 | 1 | 2 | 1 | 1 | 1 | 1 | 1 | 1 | 2 | 1 | 2 | 1 | 1 |
| **Elevation Score** | first scorer | 2 | 2 | 0 | 2 | 1 | 2 | 0 | 1 | 1 | 1 | 1 | 1 | 1 | 0 | 1 | 1 |
| | second scorer | 1 | 1 | 1 | 1 | 1 | 1 | 1 | 2 | 1 | 1 | 1 | 2 | 1 | 0 | 1 | 0 |
| **Eschar Score** | first scorer | 0 | 0 | 0 | 0 | 0 | 0 | 0 | 0 | 0 | 0 | 0 | 0 | 0 | 0 | 0 | 0 |
| | second scorer | 0 | 0 | 0 | 0 | 0 | 0 | 0 | 0 | 0 | 0 | 0 | 0 | 0 | 0 | 0 | 0 |
| **Total Score** | averaged | **3.5** | **2.5** | **2.5** | **2.5** | **3** | **2.5** | **1.5** | **2.5** | **2** | **2** | **2** | **3.5** | **2** | **2** | **2** | **1.5** |

Day 1 | Day 2 | Day 3 | Day 4 | Day 5 | Day 6 | Day 7 | Day 8

**# Animal :** **8**

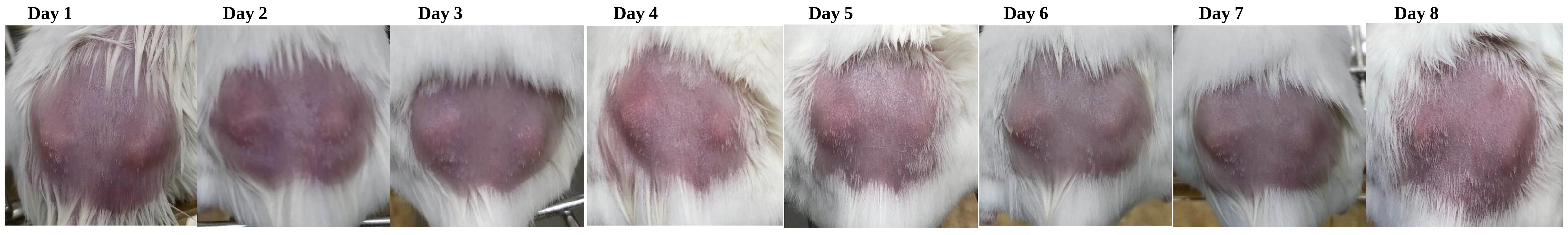

| | | Day 1 L | Day 1 R | Day 2 L | Day 2 R | Day 3 L | Day 3 R | Day 4 L | Day 4 R | Day 5 L | Day 5 R | Day 6 L | Day 6 R | Day 7 L | Day 7 R | Day 8 L | Day 8 R |
|---|---|---|---|---|---|---|---|---|---|---|---|---|---|---|---|---|---|
| **Diameter Score** | measured | 1 | 2 | 2 | 2 | 2 | 2 | 2 | 2 | 1 | 1 | 1 | 2 | 1 | 1 | 1 | 1 |
| **Elevation Score** | first scorer | 1 | 2 | 2 | 1 | 2 | 1 | 1 | 2 | 1 | 1 | 1 | 1 | 1 | 1 | 1 | 1 |
| | second scorer | 1 | 2 | 1 | 1 | 1 | 1 | 1 | 1 | 1 | 0 | 1 | 2 | 0 | 1 | 1 | 2 |
| **Eschar Score** | first scorer | 0 | 0 | 0 | 0 | 0 | 0 | 0 | 0 | 0 | 0 | 0 | 0 | 0 | 0 | 0 | 0 |
| | second scorer | 0 | 0 | 0 | 0 | 0 | 0 | 0 | 0 | 0 | 0 | 0 | 0 | 0 | 0 | 0 | 0 |
| **Total Score** | averaged | **2** | **4** | **3.5** | **3** | **3.5** | **3** | **3** | **3.5** | **2** | **1.5** | **2** | **3.5** | **1.5** | **2** | **2** | **2.5** |

Day 1 | Day 2 | Day 3 | Day 4 | Day 5 | Day 6 | Day 7 | Day 8

**# Animal :** **9**

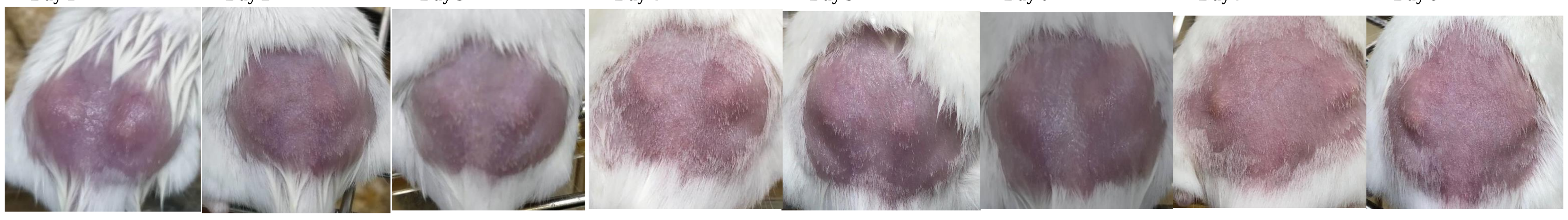

| | | Day 1 L | Day 1 R | Day 2 L | Day 2 R | Day 3 L | Day 3 R | Day 4 L | Day 4 R | Day 5 L | Day 5 R | Day 6 L | Day 6 R | Day 7 L | Day 7 R | Day 8 L | Day 8 R |
|---|---|---|---|---|---|---|---|---|---|---|---|---|---|---|---|---|---|
| **Diameter Score** | measured | 1 | 2 | 2 | 2 | 1 | 3 | 1 | 2 | 1 | 1 | 1 | 2 | 1 | 2 | 1 | 2 |
| **Elevation Score** | first scorer | 2 | 2 | 0 | 2 | 1 | 2 | 1 | 1 | 1 | 0 | 1 | 1 | 1 | 1 | 1 | 1 |
| | second scorer | 1 | 1 | 0 | 1 | 1 | 1 | 2 | 2 | 2 | 1 | 0 | 1 | 1 | 1 | 1 | 1 |
| **Eschar Score** | first scorer | 0 | 0 | 0 | 0 | 0 | 0 | 0 | 0 | 0 | 0 | 0 | 0 | 0 | 0 | 0 | 0 |
| | second scorer | 0 | 0 | 0 | 0 | 0 | 0 | 0 | 0 | 0 | 0 | 0 | 0 | 0 | 0 | 0 | 0 |
| **Total Score** | averaged | **2.5** | **3.5** | **2** | **3.5** | **2** | **4.5** | **2.5** | **3.5** | **2.5** | **1.5** | **1.5** | **3** | **2** | **3** | **2** | **3** |

Day 1 | Day 2 | Day 3 | Day 4 | Day 5 | Day 6 | Day 7 | Day 8

**# Animal :** **10**

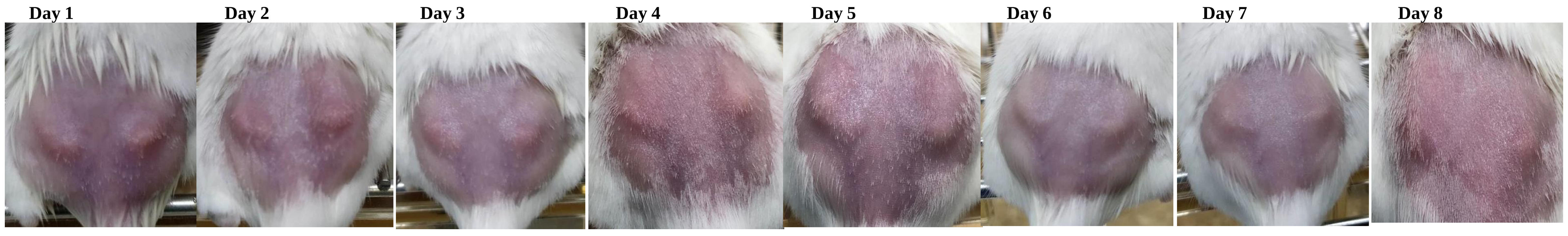

| | | Day 1 L | Day 1 R | Day 2 L | Day 2 R | Day 3 L | Day 3 R | Day 4 L | Day 4 R | Day 5 L | Day 5 R | Day 6 L | Day 6 R | Day 7 L | Day 7 R | Day 8 L | Day 8 R |
|---|---|---|---|---|---|---|---|---|---|---|---|---|---|---|---|---|---|
| **Diameter Score** | measured | 2 | 3 | 1 | 2 | 2 | 2 | 1 | 2 | 2 | 2 | 1 | 2 | 1 | 2 | 1 | 2 |
| **Elevation Score** | first scorer | 2 | 1 | 2 | 2 | 1 | 1 | 1 | 1 | 1 | 1 | 1 | 1 | 1 | 1 | 1 | 1 |
| | second scorer | 2 | 1 | 2 | 2 | 1 | 1 | 1 | 1 | 1 | 0 | 1 | 1 | 1 | 1 | 1 | 1 |
| **Eschar Score** | first scorer | 0 | 0 | 0 | 0 | 0 | 0 | 0 | 0 | 0 | 0 | 0 | 0 | 0 | 0 | 0 | 0 |
| | second scorer | 0 | 0 | 0 | 0 | 0 | 0 | 0 | 0 | 0 | 0 | 0 | 0 | 0 | 0 | 0 | 0 |
| **Total Score** | averaged | **4** | **4** | **3** | **4** | **3** | **3** | **2** | **3** | **3** | **2.5** | **2** | **3** | **2** | **3** | **2** | **3** |

Day 1 | Day 2 | Day 3 | Day 4 | Day 5 | Day 6 | Day 7 | Day 8

**# Animal :** **11**

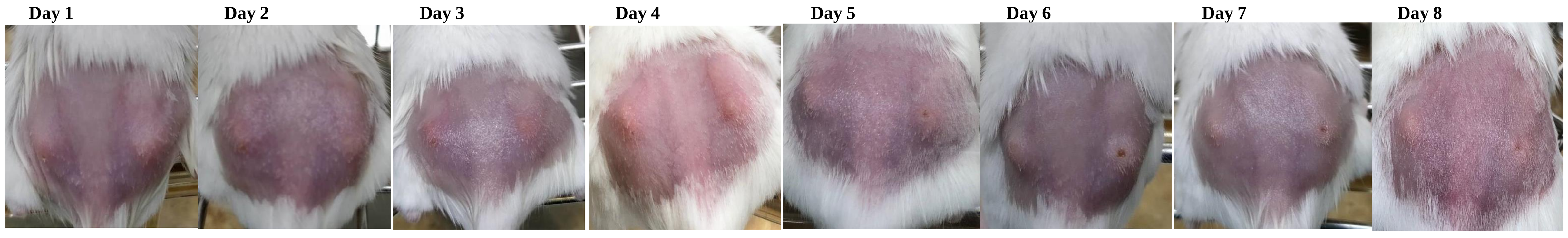

| | | Day 1 L | Day 1 R | Day 2 L | Day 2 R | Day 3 L | Day 3 R | Day 4 L | Day 4 R | Day 5 L | Day 5 R | Day 6 L | Day 6 R | Day 7 L | Day 7 R | Day 8 L | Day 8 R |
|---|---|---|---|---|---|---|---|---|---|---|---|---|---|---|---|---|---|
| **Diameter Score** | measured | 2 | 2 | 2 | 2 | 2 | 2 | 1 | 1 | 1 | 0 | 2 | 1 | 1 | 1 | 1 | 0 |
| **Elevation Score** | first scorer | 1 | 0 | 1 | 0 | 0 | 1 | 1 | 0 | 0 | 0 | 1 | 1 | 0 | 1 | 1 | 0 |
| | second scorer | 2 | 1 | 0 | 1 | 1 | 1 | 1 | 1 | 1 | 1 | 1 | 1 | 1 | 1 | 1 | 0 |
| **Eschar Score** | first scorer | 0 | 0 | 0 | 0 | 0 | 0 | 0 | 0 | 0 | 0 | 0 | 1 | 0 | 0 | 0 | 0 |
| | second scorer | 0 | 0 | 0 | 0 | 1 | 0 | 0 | 0 | 0 | 1 | 0 | 1 | 0 | 1 | 0 | 1 |
| **Total Score** | averaged | **3.5** | **2.5** | **2.5** | **2.5** | **3** | **3** | **2** | **1.5** | **1.5** | **1** | **3** | **3** | **1.5** | **2.5** | **2** | **0.5** |

Day 1 | Day 2 | Day 3 | Day 4 | Day 5 | Day 6 | Day 7 | Day 8

**# Animal :** **12**

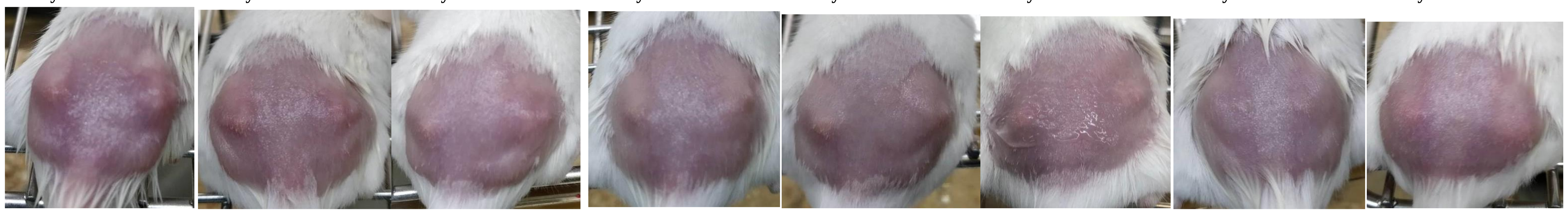

| | | Day 1 L | Day 1 R | Day 2 L | Day 2 R | Day 3 L | Day 3 R | Day 4 L | Day 4 R | Day 5 L | Day 5 R | Day 6 L | Day 6 R | Day 7 L | Day 7 R | Day 8 L | Day 8 R |
|---|---|---|---|---|---|---|---|---|---|---|---|---|---|---|---|---|---|
| **Diameter Score** | measured | 3 | 2 | 2 | 2 | 3 | 2 | 2 | 1 | 2 | 1 | 2 | 1 | 1 | 0 | 1 | 1 |
| **Elevation Score** | first scorer | 2 | 2 | 2 | 1 | 2 | 1 | 1 | 2 | 2 | 2 | 2 | 1 | 2 | 1 | 1 | 1 |
| | second scorer | 3 | 3 | 2 | 1 | 2 | 1 | 2 | 1 | 2 | 1 | 2 | 1 | 1 | 0 | 1 | 1 |
| **Eschar Score** | first scorer | 0 | 0 | 0 | 0 | 0 | 0 | 0 | 0 | 0 | 0 | 0 | 0 | 0 | 0 | 0 | 0 |
| | second scorer | 0 | 0 | 0 | 0 | 0 | 0 | 0 | 0 | 0 | 0 | 0 | 0 | 0 | 0 | 0 | 0 |
| **Total Score** | averaged | **5.5** | **4.5** | **4** | **3** | **5** | **3** | **3.5** | **2.5** | **4** | **2.5** | **4** | **2** | **2.5** | **0.5** | **2** | **2** |

### Animal : 13

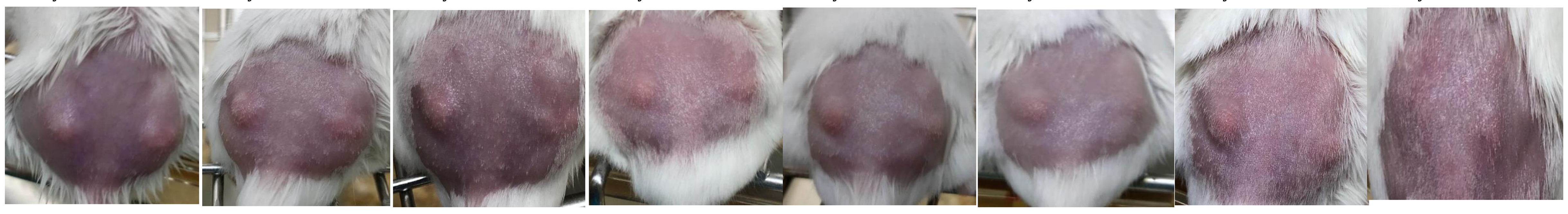

| | | Day 1 | | Day 2 | | Day 3 | | Day 4 | | Day 5 | | Day 6 | | Day 7 | | Day 8 | |
|---|---|---|---|---|---|---|---|---|---|---|---|---|---|---|---|---|---|
| | | **L** | **R** | **L** | **R** | **L** | **R** | **L** | **R** | **L** | **R** | **L** | **R** | **L** | **R** | **L** | **R** |
| **Diameter Score** | measured | 2 | 2 | 2 | 2 | 3 | 2 | 2 | 1 | 2 | 1 | 2 | 0 | 2 | 0 | 2 | 0 |
| **Elevation Score** | first scorer | 1 | 1 | 1 | 1 | 2 | 1 | 2 | 1 | 2 | 1 | 1 | 1 | 1 | 1 | 1 | 0 |
| | second scorer | 2 | 2 | 1 | 1 | 2 | 1 | 2 | 1 | 2 | 1 | 1 | 0 | 1 | 0 | 0 | 1 |
| **Eschar Score** | first scorer | 0 | 0 | 0 | 0 | 0 | 0 | 0 | 0 | 0 | 0 | 0 | 0 | 0 | 0 | 0 | 0 |
| | second scorer | 0 | 0 | 0 | 0 | 0 | 0 | 0 | 0 | 0 | 0 | 0 | 0 | 0 | 0 | 0 | 0 |
| **Total Score** | averaged | **3.5** | **3.5** | **3** | **3** | **5** | **3** | **4** | **2** | **4** | **2** | **3** | **0.5** | **3** | **0.5** | **2.5** | **0.5** |

### Animal : 14

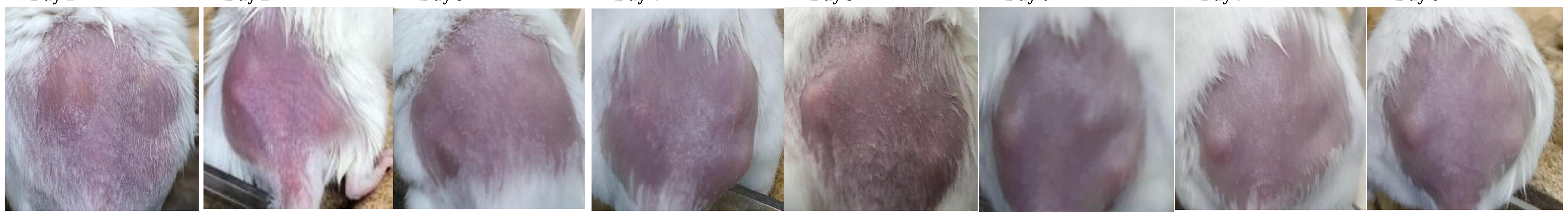

| | | Day 1 | | Day 2 | | Day 3 | | Day 4 | | Day 5 | | Day 6 | | Day 7 | | Day 8 | |
|---|---|---|---|---|---|---|---|---|---|---|---|---|---|---|---|---|---|
| | | **L** | **R** | **L** | **R** | **L** | **R** | **L** | **R** | **L** | **R** | **L** | **R** | **L** | **R** | **L** | **R** |
| **Diameter Score** | measured | 1 | 2 | 1 | 2 | 1 | 0 | 1 | 0 | 2 | 1 | 2 | 2 | 2 | 1 | 2 | 1 |
| **Elevation Score** | first scorer | 1 | 0 | 1 | 0 | 0 | 0 | 1 | 1 | 2 | 1 | 2 | 1 | 2 | 1 | 2 | 0 |
| | second scorer | 0 | 0 | 1 | 0 | 0 | 0 | 0 | 0 | 1 | 1 | 1 | 1 | 2 | 1 | 2 | 1 |
| **Eschar Score** | first scorer | 0 | 0 | 0 | 0 | 0 | 0 | 0 | 0 | 0 | 0 | 0 | 0 | 0 | 0 | 0 | 0 |
| | second scorer | 0 | 0 | 0 | 0 | 0 | 0 | 0 | 0 | 0 | 0 | 0 | 0 | 0 | 0 | 0 | 0 |
| **Total Score** | averaged | **1.5** | **2** | **2** | **2** | **1** | **0** | **1.5** | **0.5** | **3.5** | **2** | **3.5** | **3** | **4** | **2** | **4** | **1.5** |

### Animal : 15

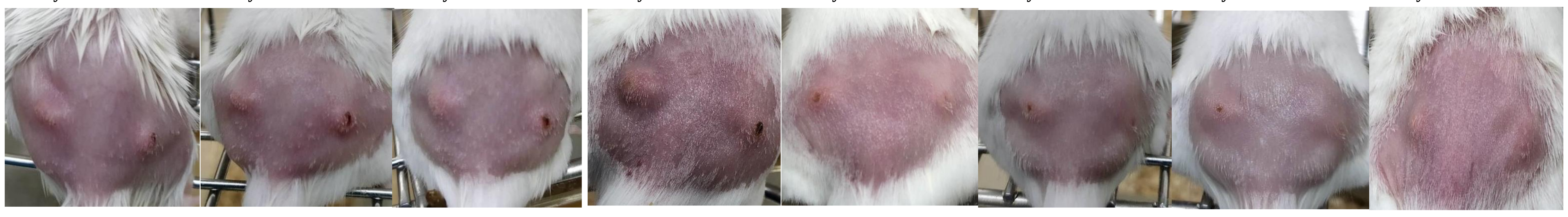

| | | Day 1 | | Day 2 | | Day 3 | | Day 4 | | Day 5 | | Day 6 | | Day 7 | | Day 8 | |
|---|---|---|---|---|---|---|---|---|---|---|---|---|---|---|---|---|---|
| | | **L** | **R** | **L** | **R** | **L** | **R** | **L** | **R** | **L** | **R** | **L** | **R** | **L** | **R** | **L** | **R** |
| **Diameter Score** | measured | 2 | 2 | 2 | 2 | 3 | 1 | 2 | 1 | 2 | 1 | 2 | 2 | 2 | 1 | 2 | 1 |
| **Elevation Score** | first scorer | 2 | 1 | 2 | 2 | 2 | 1 | 2 | 1 | 1 | 1 | 1 | 1 | 1 | 0 | 1 | 0 |
| | second scorer | 2 | 1 | 2 | 2 | 2 | 1 | 2 | 1 | 1 | 1 | 1 | 1 | 1 | 1 | 1 | 1 |
| **Eschar Score** | first scorer | 0 | 1 | 0 | 2 | 0 | 2 | 0 | 2 | 1 | 1 | 1 | 0 | 1 | 0 | 0 | 0 |
| | second scorer | 0 | 2 | 0 | 2 | 0 | 2 | 0 | 2 | 2 | 0 | 2 | 0 | 1 | 0 | 0 | 0 |
| **Total Score** | averaged | **4** | **4.5** | **4** | **6** | **5** | **4** | **4** | **4** | **4.5** | **2.5** | **4.5** | **3** | **4** | **1.5** | **3** | **1.5** |

### Animal : 16

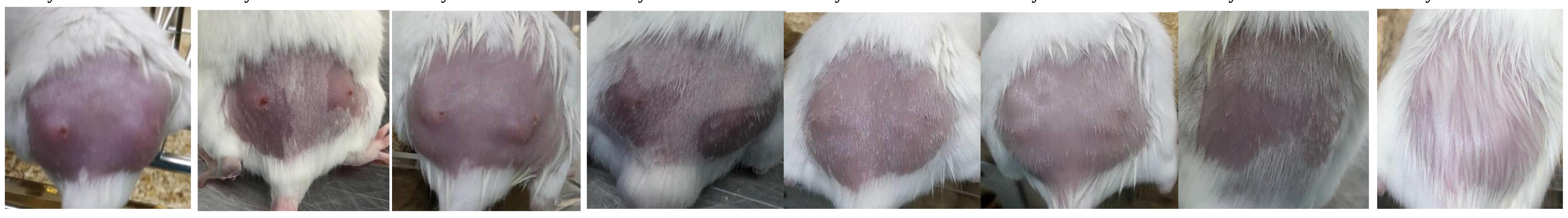

| | | Day 1 | | Day 2 | | Day 3 | | Day 4 | | Day 5 | | Day 6 | | Day 7 | | Day 8 | |
|---|---|---|---|---|---|---|---|---|---|---|---|---|---|---|---|---|---|
| | | **L** | **R** | **L** | **R** | **L** | **R** | **L** | **R** | **L** | **R** | **L** | **R** | **L** | **R** | **L** | **R** |
| **Diameter Score** | measured | 3 | 2 | 2 | 1 | 2 | 1 | 0 | 0 | 1 | 1 | 1 | 0 | 0 | 0 | 0 | 0 |
| **Elevation Score** | first scorer | 1 | 1 | 1 | 1 | 1 | 1 | 0 | 0 | 0 | 0 | 0 | 0 | 0 | 0 | 0 | 0 |
| | second scorer | 1 | 1 | 0 | 1 | 1 | 1 | 0 | 0 | 0 | 0 | 0 | 0 | 0 | 0 | 0 | 0 |
| **Eschar Score** | first scorer | 2 | 0 | 2 | 1 | 1 | 1 | 1 | 0 | 0 | 0 | 0 | 0 | 0 | 0 | 0 | 0 |
| | second scorer | 2 | 0 | 2 | 2 | 1 | 0 | 1 | 0 | 0 | 0 | 0 | 0 | 0 | 0 | 0 | 0 |
| **Total Score** | averaged | **6** | **3** | **4.5** | **3.5** | **4** | **2.5** | **1** | **0** | **1** | **1** | **1** | **0** | **0** | **0** | **0** | **0** |

### Animal : 17

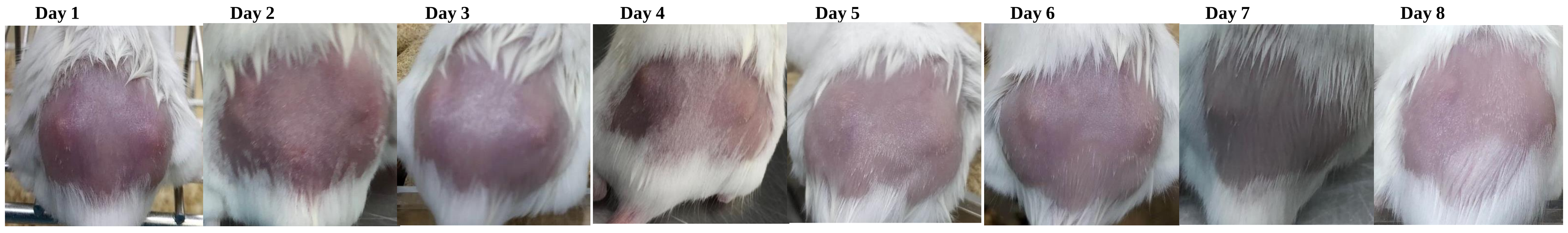

| | | Day 1 | | Day 2 | | Day 3 | | Day 4 | | Day 5 | | Day 6 | | Day 7 | | Day 8 | |
|---|---|---|---|---|---|---|---|---|---|---|---|---|---|---|---|---|---|
| | | **L** | **R** | **L** | **R** | **L** | **R** | **L** | **R** | **L** | **R** | **L** | **R** | **L** | **R** | **L** | **R** |
| **Diameter Score** | measured | 2 | 2 | 2 | 2 | 2 | 2 | 2 | 1 | 2 | 1 | 2 | 1 | 1 | 1 | 1 | 1 |
| **Elevation Score** | first scorer | 2 | 2 | 2 | 1 | 1 | 1 | 1 | 1 | 1 | 1 | 2 | 1 | 1 | 1 | 1 | 0 |
| | second scorer | 2 | 2 | 1 | 1 | 1 | 1 | 1 | 1 | 1 | 1 | 1 | 1 | 1 | 1 | 1 | 0 |
| **Eschar Score** | first scorer | 0 | 0 | 0 | 0 | 0 | 0 | 0 | 0 | 0 | 0 | 0 | 0 | 0 | 0 | 0 | 0 |
| | second scorer | 0 | 0 | 0 | 0 | 0 | 0 | 0 | 0 | 0 | 0 | 0 | 0 | 0 | 0 | 0 | 0 |
| **Total Score** | averaged | **4** | **4** | **3.5** | **3** | **3** | **3** | **3** | **2** | **3** | **2** | **3.5** | **2** | **2** | **2** | **2** | **1** |
